# Receptor-like cytoplasmic kinases belonging to different subfamilies mediate immune responses downstream of the Cf-4 resistance protein in *Nicotiana benthamiana*

**DOI:** 10.1101/2023.04.25.538242

**Authors:** Wen R.H. Huang, Ciska Braam, Carola Kretschmer, Huan Liu, Filiz Ferik, Aranka M. van der Burgh, Max Pluis, Agnes Omabour Hagan, Sergio Landeo Villanueva, Jinbin Wu, Amber van Loosbroek, Yulu Wang, Michael F. Seidl, Johannes Stuttmann, Matthieu H.A.J. Joosten

**Affiliations:** Laboratory of Phytopathology, Wageningen University, Droevendaalsesteeg 1, 6708 PB Wageningen, The Netherlands; Institute of Biology, Department of Genetics, Martin- Luther- University, Halle-Wittenberg, Weinbergweg 10, D- 06120 Halle (Saale), Germany; Plant Breeding, Wageningen University & Research, Droevendaalsesteeg 1, 6708 PB Wageningen, The Netherlands; Laboratory of Biomanufacturing and Food Engineering, Institute of Food Science and Technology, Chinese Academy of Agricultural Sciences, Beijing 100193, China; Theoretical Biology & Bioinformatics, Department of Biology, Utrecht University, Utrecht, the Netherlands

## Abstract

Cell-surface receptors, which are either receptor-like proteins (RLPs) or receptor-like kinases (RLKs), form the first layer of the plant innate immune system. The LRR-RLKs SUPPRESSOR OF BIR1-1 (SOBIR1) and BRASSINOSTEROID INSENSITIVE 1-ASSOCIATED KINASE 1 (BAK1) are the common regulatory co-receptors of LRR-RLPs. The tomato LRR-RLP Cf-4, which specifically detects the apoplastic effector Avr4 that is secreted by the pathogenic intercellular fungus *Cladosporium fulvum*, requires SOBIR1 and BAK1 to mediate resistance against this fungus. It is proposed that phosphorylation events take place between the cytoplasmic kinase domains of SOBIR1 and BAK1, which lead to the initiation of various downstream signaling outputs such as the rapid phosphorylation of receptor-like cytoplasmic kinases (RLCKs), the accumulation of reactive oxygen species (ROS), the activation of mitogen-activated protein kinase (MAPK) cascades, and in some cases, the hypersensitive response (HR). Here, by performing *in vitro* phosphorylation assays, we show that SOBIR1 directly phosphorylates BAK1, whereas BAK1, in its turn, directly phosphorylates SOBIR1, which results in the full activation of the Cf-4-containing immune complex. We found that threonine 522, present in the activation segment of the SOBIR1 kinase domain of *Nicotiana benthamiana* (*Nb*), is required for its intrinsic kinase activity, and is essential for the Avr4/Cf-4/SOBIR1/BAK1-triggered ROS burst, MAPK activation and the HR. Tyrosine residue Y469 was found to be crucial for the Avr4/Cf-4- triggered activation of MAPKs and the HR, but not for ROS production and *Nb*SOBIR1 intrinsic kinase activity. RLCKs are well-recognized to act as the initial cytoplasmic transducers, bridging cell-surface receptor complexes with their downstream signaling partners. By knocking out multiple genes belonging to different RLCK-VII subfamilies in *N. benthamiana* plants stably expressing *Cf-4*, we show that members of the RLCK-VII subfamilies 6, 7 and 8 are required for the Avr4/Cf-4-triggered ROS production. Typically, the Avr4 protein triggers a biphasic ROS burst in leaf discs obtained from *N. benthamiana:Cf-4* plants, in which the first transitory response is followed by a second, sustained ROS burst. Interestingly, the intensities of the first and second phase of the ROS burst are affected in different ways in the various *rlck-vii* knock-out plants, indicating that there are different downstream ROS regulatory mechanisms, involving different RLCKs in *N. benthamiana*. Further studies show that members from RLCK-VII-6, 7 and 8 also play an essential role in regulating the ROS production triggered by flg22, chitin, and the nlp20/RLP23 and the pg13/RLP42 combinations.

## INTRODUCTION

Plants have evolved a two-layered innate immune system to fend off invading microbes, of which the first layer is mediated by cell-surface receptors (van der Burgh and Joosten, 2019). The two largest families of such receptors are formed by receptor-like proteins (RLPs) and receptor-like kinases (RLKs) (DeFalco and Zipfel, 2021; Macho and Zipfel, 2014; Monaghan and Zipfel, 2012; Ranf, 2017). Both RLKs and RLPs detect extracellular immunogenic patterns (ExIPs), leading to extracellularly-triggered immunity (ExTI) (van der Burgh and Joosten, 2019). These ExIPs can be any extracellular danger signal, including pathogen-derived immunogenic patterns and effectors, and host-derived damage-associated molecular patterns (van der Burgh and Joosten, 2019). The second layer of plant innate immunity is mediated by intracellular receptors, which specifically recognize intracellular immunogenic patterns (InIPs), leading to intracellularly-triggered immunity (InTI) (van der Burgh and Joosten, 2019). Both ExTI and InTI are associated with various downstream immune outputs, including the rapid phosphorylation of downstream receptor-like cytoplasmic kinases (RLCKs), the swift production of apoplastic reactive oxygen species (ROS), the activation of mitogen-activated protein kinase (MAPK) cascades, large-scale transcriptional reprogramming, and in some cases, the induction of programmed cell death, referred to as the hypersensitive response (HR) (Lu and Tsuda, 2021; Tsuda and Katagiri, 2010; Yuan et al., 2021).

RLKs and RLPs share the same overall structure (Macho and Zipfel, 2014; Monaghan and Zipfel, 2012; Steinbrenner, 2020). They both are composed of an ectodomain that is potentially involved in ligand binding and a single-pass transmembrane domain. However, in contrast to RLKs, RLPs lack an intracellular kinase domain for downstream signaling (Chisholm et al., 2006; Lee et al., 2021; Monaghan and Zipfel, 2012; Wang and Chai, 2020). To date, the most extensively studied plant cell-surface receptors are the families of leucine-rich repeat (LRR)-RLKs and LRR-RLPs (Kong et al., 2021; Lee et al., 2021). Typically, LRR-RLKs form a complex with the regulatory LRR-RLK BRASSINOSTEROID INSENSITIVE 1-ASSOCIATED KINASE 1 (BAK1), also referred to as SOMATIC EMBRYOGENESIS RECEPTOR KINASE 3 (SERK3, further referred to as BAK1), in a ligand-dependent manner (Albert et al., 2020; Chinchilla et al., 2009; Heese et al., 2007; Ranf, 2017; Wan, W.-L. et al., 2019). Well-studied examples are the Arabidopsis (*Arabidopsis thaliana*, *At*) LRR-RLKs FLAGELLIN-SENSING 2 (FLS2) and EF-Tu RECEPTOR (EFR), which specifically recognize bacterial flagellin (or its N-terminal 22-amino acid epitope, flg22) and bacterial elongation factor TU (EF-Tu) (or its N-terminal 18-amino acid epitope, elf18), respectively, resulting in the rapid trans-phosphorylation of the cytoplasmic kinase domains of BAK1 and FLS2/EFR (Chinchilla et al., 2009; Chinchilla et al., 2007; Gómez-Gómez et al., 2001; Gómez-Gómez and Boller, 2000; Heese et al., 2007; Sun et al., 2013; Zipfel et al., 2006).

LRR-RLPs, which lack an intracellular kinase domain, have been found to constitutively associate with the LRR-RLK SUPPRESSOR OF BIR1-1 (SOBIR1), also referred to as EVERSHED (further referred to as SOBIR1), and to require SOBIR1 for their function in plant immunity (Gao et al., 2009; Liebrand et al., 2013; Liebrand et al., 2014). RLPs are omnipresent in plants, and SOBIR1 is also widely conserved in all sequenced plant genomes for which information is available (Liebrand et al., 2014). The tomato (*Solanum lycopersicum*, *Sl*) resistance protein Cf-4 is one of the best-studied LRR-RLPs and specifically recognizes the apoplastic effector Avr4 that is secreted by the pathogenic intercellular fungus *Cladosporium fulvum* (Joosten et al., 1994; Thomas et al., 1997). Cf- 4 forms a complex with SOBIR1 in a ligand-independent manner, which is essential for Cf- 4-mediated resistance to *C. fulvum* (Liebrand et al., 2013; Liebrand et al., 2014). In addition, Arabidopsis RLP23 requires SOBIR1 to mount resistance against various oomycete and fungal plant pathogens that secrete necrosis and ethylene-inducing peptide 1-like proteins (NLPs) (Albert et al., 2015). Consistent with LRR-RLKs, the LRR-RLP/SOBIR1 complex also requires the recruitment of BAK1 for the initiation of downstream signaling upon ligand perception (Albert et al., 2015; DeFalco and Zipfel, 2021; Postma et al., 2016).

Protein phosphorylation, which is a swift and reversible biochemical post-translational modification, plays an important role as a versatile molecular switch in various cellular activities, including the initiation of plant immune responses (Kong et al., 2021; Macho and Zipfel, 2014; Mithoe and Menke, 2018). According to whether a conserved arginine (Arg/R) is immediately preceding the highly conserved catalytic aspartate (Asp/D) in their catalytic loop, protein kinases can be subdivided into RD and non-RD kinases (Johnson et al., 1996). Generally, activation of RD kinases requires phosphorylation of one or more residues in their activation segment (Johnson et al., 1996; Nolen et al., 2004; Oliver et al., 2007). In contrast, non-RD kinases require a regulatory RD kinase, such as BAK1, to promote their phosphorylation and signaling (Dardick et al., 2012). Unlike the majority of mammalian receptor kinases that possess tyrosine (Tyr/Y) kinase activity in addition to serine (Ser/S)/threonine (Thr/T) kinase activity, plant RLKs were traditionally annotated as serine/threonine-specific kinases (Bleecker, 2001; G. Manning, 2002; Wendell A. Lim, 2010). However, this knowledge has been revised over the past twenty years, as emerging evidence shows that plant RLKs also exhibit tyrosine phosphorylation activity, in addition to phosphorylate at serine/threonine residues. A good example is Arabidopsis BAK1, which is a dual-specificity kinase that can phosphorylate at both serine/threonine and tyrosine residues (Lin et al., 2014; Wang et al., 2005; Wang et al., 2008). Phosphorylation of Thr455 in the activation segment of BAK1 is essential for obtaining its kinase activity (Wang et al., 2005; Wang et al., 2008), while phosphorylation of Tyr403 is required for BAK1 function in downstream immune signaling, as an amino acid change at Tyr403 strongly reduced flg22- and elf18-triggered ROS production and flg22-triggered MAPK activation, but does not impair the function of BAK1 in plant development (Perraki et al., 2018).

Activation of cell-surface receptor complexes subsequently triggers a suite of downstream signaling events. Increasing evidence suggests that RLCKs are the direct downstream cytoplasmic substrates of activated receptor complexes and that they fill the gap between receptors present at the cell surface and downstream signaling components (Bi et al., 2018; Lin et al., 2013; Macho and Zipfel, 2014; Tang et al., 2017; Zhou and Zhang, 2020). Arabidopsis BOTRYTIS-INDUCED KINASE 1 (BIK1) is one of the best-characterized RLCKs and is required for triggering ROS production upon the perception of multiple ExIPs, such as flg22, elf18, and chitin (Lu et al., 2010; Wan, W.L. et al., 2019; Zhang et al., 2010). In addition to BIK1, Arabidopsis AvrPphB SUSCEPTIBLE 1-LIKE 27 (PBL27) has been reported to connect chitin perception to MAPK cascade activation, as MAPKKK5 is directly phosphorylated by PBL27 (Yamada et al., 2016). BIK1, PBL27 and some other RLCKs that have been identified as central positive regulatory components that relay immune signaling from the cell surface to the intracellular space, all belong to Arabidopsis RLCK class VII (Lin et al., 2015; Luo et al., 2020; Pruitt et al., 2021; Zhang et al., 2010). RLCK-VII is composed of 46 members, and according to their sequence similarity, these members can be further divided into nine subfamilies, termed RLCK-VII-1 to RLCK-VII-9 (Rao et al., 2018). Interestingly, flg22-, elf18- and chitin-triggered ROS production is significantly reduced in *rlck-vii-5*, *-7*, and *-8* knock-out Arabidopsis plants, whereas in contrast, *rlck-vii-6* knock-out plants exhibit higher flg22-induced ROS accumulation levels when compared to wild-type (WT) plants (Rao et al., 2018).

The last two decades have witnessed remarkable progress in our understanding of the initiation and regulation of plant innate immunity. Nevertheless, it is largely unknown how SOBIR1 and BAK1 exactly trans-phosphorylate each other to initiate downstream signaling and whether tyrosine residues that are present in the kinase domain of SOBIR1 play an essential role in regulating LRR-RLP/SOBIR1-mediated signaling. Additionally, compared to the model plant Arabidopsis, little is known about the function of the various RLCK-VII subfamilies in Solanaceous plants, and it is thus far not known which RLCKs are involved in the LRR-RLP/SOBIR1-triggered immune signaling pathway. In this study, we performed a site-directed mutagenesis screen combined with a complementation study, in *Nicotiana benthamiana* plants stably expressing tomato *Cf-4* and in which the functional *SOBIR1* gene has been knocked out. *Nb*SOBIR1 Thr522, as well as its analogous residues in both tomato SOBIR1s, present in the activation segment of the kinase domain of SOBIR1, was found to play an essential role in Avr4/Cf-4-triggered immune signaling, including the initiation of the ROS burst, MAPK activation and the HR. Interestingly, *in vitro* phosphorylation assays demonstrate that this highly conserved Thr residue is required for SOBIR1(-like) intrinsic kinase activity. Besides, we show that SOBIR1 directly trans-phosphorylates BAK1, whereas, on the other hand, BAK1 is also able to directly trans-phosphorylate SOBIR1. These trans-phosphorylation events are proposed to eventually lead to the full activation of both SOBIR1 and BAK1, and their intrinsic kinase activity is required for these trans-phosphorylation events to take place and the initiation of downstream immune signaling. In addition to Thr522, *Nb*SOBIR1 Tyr469, as well as its analogous residues in *Sl*SOBIR1 and *Sl*SOBIR1-like, was identified to be essential for Avr4/Cf-4-triggered MAPK activation and the initiation of the HR, but not for activating ROS production. RLCKs are the initial cytoplasmic transducers of the extracellular signal that is perceived by cell-surface receptors. By knocking out multiple candidate genes belonging to different RLCK-VII subfamilies in transgenic *N. benthamiana* plants stably expressing *Cf-4*, we show that members from RLCK-VII-6, −7 and −8 are required for full Avr4/Cf-4- triggered biphasic ROS production, even though these RLCKs might be involved in different mechanisms actually regulating ROS production. Additionally, these RLCKs are also vital for initiating ROS production triggered by flg22, chitin, and the matching nlp20/RLP23 and pg13/RLP42 ligand/receptor combinations in *N. benthamiana*.

## RESULTS

### Thr522, present in the activation segment of the kinase domain of *Nb*SOBIR1, is essential for mounting the Avr4/Cf-4-triggered immune signaling

The regulatory LRR-RLK SOBIR1 appears to be present throughout the plant kingdom, including *N. benthamiana* (*Nb*), which is a versatile experimental host plant (Gao et al., 2009; Liebrand et al., 2013; Liebrand et al., 2014). SOBIR1 is an RD kinase, which suggests that likely SOBIR1 first requires phosphorylation of its activation segment to acquire the kinase-active conformation (Johnson et al., 1996; Oliver et al., 2007). It has been reported that the kinase domain of *At*SOBIR1 auto-phosphorylates at Ser, Thr, and Tyr residues *in vitro* (Leslie et al., 2010), and in the activation segment of SOBIR1 of tomato and *N. benthamiana* such residues are present. To investigate which residue(s) in the kinase domain of SOBIR1 is(are) essential for the activation of Cf-4/SOBIR1-triggered signaling pathway, we decided to zoom in on the activation segment of *Nb*SOBIR1 (Liebrand et al., 2013). Five potential phosphorylation sites are located in this loop of 30 amino acids, including one Ser and four Thr residues (Figure 1A). Strikingly, these residues are highly conserved in SOBIR1 from various plant species, including tomato (Figure S1 and S2A).

**Figure 1.**
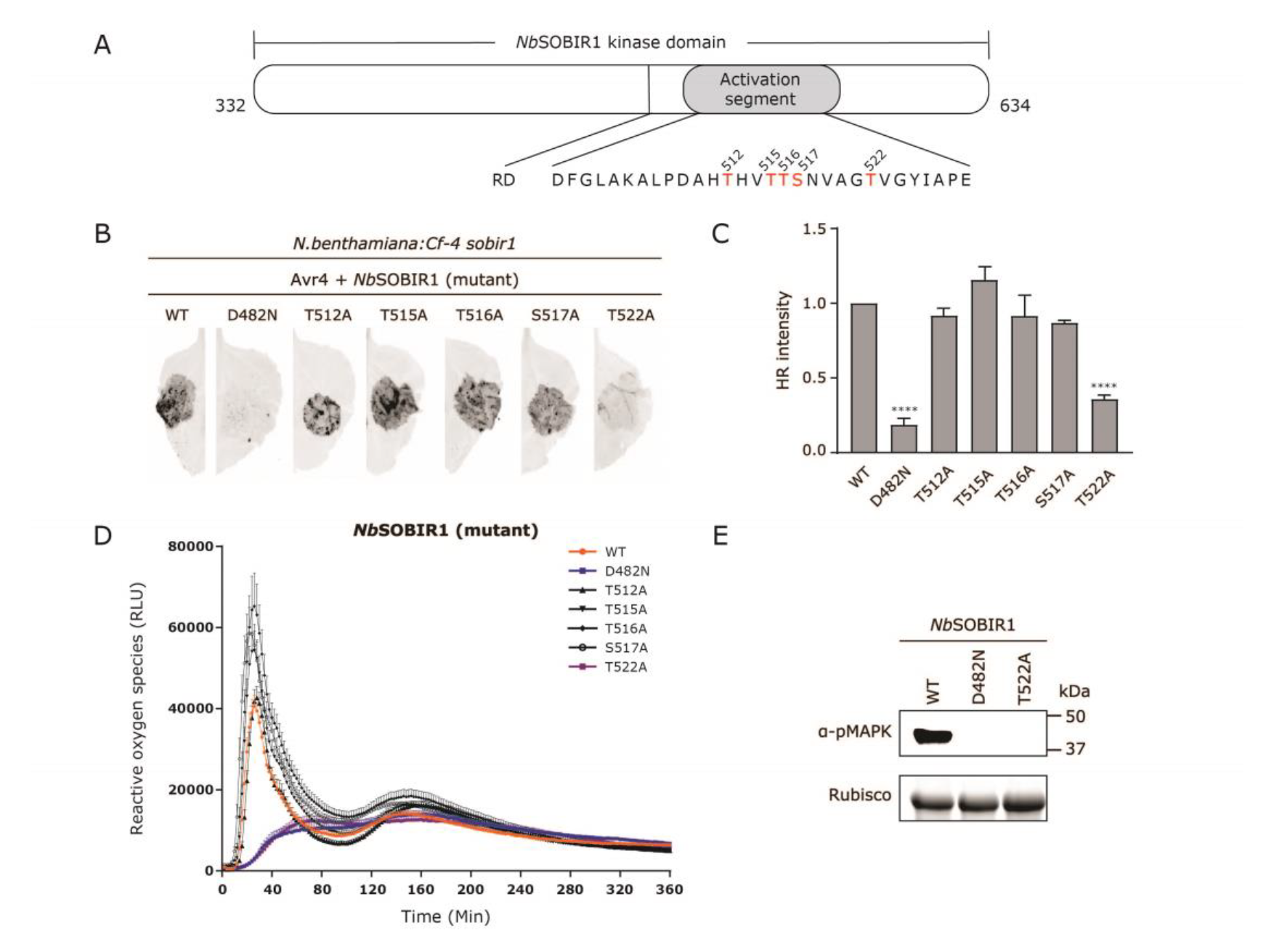
Thr522, Present in the Activation Segment of the *Nb*SOBIR1 Kinase Domain, Is Essential for Avr4/Cf-4/SOBIR1/BAK1-Triggered Immune Signaling. **(A)** Schematic diagram of the kinase domain of *Nb*SOBIR1, with the RD motif and the activation segment indicated. The amino acid sequence of the activation segment of *Nb*SOBIR1 is shown below the diagram. Possible phosphorylation sites are denoted in red. **(B** to **E)** Mutagenesis screen in combination with complementation of all potential phosphorylation sites in the activation segment of *Nb*SOBIR1 to determine their importance in Avr4/Cf-4-triggered immune signaling. **(B)** All SOBIR1 Thr and Ser mutants were transiently co-expressed with *Avr4* by agro-infiltration (OD600 = 0.8) in leaves of *N. benthamiana:Cf-4 sobir1* knock-out plants. Wild-type (WT) *Nb*SOBIR1 was used as a positive control, whereas its kinase-dead mutant *Nb*SOBIR1 D482N, was used as a negative control. Leaves were imaged using the ChemiDoc, with the Red Fluorescent Protein (RFP) channel (filter: 605/50; light: Green Epi Illumination; exposure time: 2 seconds), at 5 days post infiltration (dpi). **(C)** Image Lab quantification of the intensity of the HR. Data shown are the average relative intensities of the HR + standard error of the mean (SEM) (n≥6) that were triggered upon complementation with the various *Nb*SOBIR1 mutants, when compared to complementation with *Nb*SOBIR1 WT, of which the level of the HR was set to 1. Statistical significance was determined by a one-way ANOVA/Dunnett’s multiple comparison test, compared with *Nb*SOBIR1 WT. **** p < 0.0001. **(D)** Leaf discs of a *N. benthamiana:Cf-4 sobir1* knock-out line, transiently expressing the five different *Nb*SOBIR1 mutants, as well as their corresponding WT (the positive control) and kinase-dead mutant (D482N) (the negative control), were treated with 0.1 μM Avr4 protein, and ROS accumulation was monitored over time. ROS production is expressed as relative light units (RLUs), and the data are represented as mean + SEM. **(E)** The different *Nb*SOBIR1 mutants, as well as *Nb*SOBIR1 WT and D482N, were agro-infiltrated together with Avr4 in leaves of an *N. benthamiana:Cf-4 sobir1* knock-out line. Leaf samples were harvested at 2 dpi and total protein extracts were subjected to immunoblotting with a p42/p44- erk antibody to determine the activation of downstream MAPKs by phosphorylation. Experiments were repeated at least three times with similar results, and representative results are shown.

Cf-4 is functional in *N. benthamiana* and agro-infiltration of Avr4 in stable transgenic *N. benthamiana:Cf-4* plants triggers a typical HR (Gabriëls et al., 2006). Cf-4 function is completely abolished in *N. benthamiana:Cf-4 sobir1* knock-out lines, whereas complementation through transient expression of functional *NbSOBIR1* restores the Avr4/Cf-4-mediated HR in such knock-out lines (Huang et al., 2021). Based on these observations, we carried out site-directed mutagenesis of the activation segment of *Nb*SOBIR1 to substitute individual Ser/Thr residues with alanine (Ala/A) residue, which lacks the phosphorylatable hydroxyl group and thereby cannot be phosphorylated. Subsequently, we performed a complementation study with the five different *Nb*SOBIR1 mutants, taking along the *Nb*SOBIR1 WT and the corresponding D to N (asparagine/Asn) kinase-dead mutant (D482N), as a positive and a negative control, respectively (van der Burgh et al., 2019). Interestingly, in contrast to *Nb*SOBIR1 WT but similar to *Nb*SOBIR1 D482N, *Nb*SOBIR1 T522A failed to restore the Avr4/Cf-4-specific HR in *N. benthamiana:Cf- 4 sobir1* knock-out plants (Figure 1B). It is worth noting that this phenotype was not caused by a lack of accumulation of the *Nb*SOBIR1 T522A protein *in planta* (Figure S3). Quantification of the intensity of the HR, which was determined by employing red light imaging (Landeo Villanueva et al., 2021), showed that the intensity of the HR obtained upon transient co-expression of *NbSOBIR1 T522A* with *Avr4* was significantly lower than the intensity of the HR obtained when co-expressing *NbSOBIR1 WT* with *Avr4* (Figure 1C). The four additional mutants of *Nb*SOBIR1 showed a complementation capacity that was similar to *Nb*SOBIR1 WT.

The swift production of ROS is a hallmark of the plant immune response and apoplastic ROS are mainly produced by nicotinamide adenine dinucleotide phosphate (NADPH) oxidases, such as the RESPIRATORY BURST OXIDASE HOMOLOGUE B (RBOHB) from *N. benthamiana* and tomato, which localizes at the plasma membrane (Qi et al., 2017; Waszczak et al., 2018; Yu et al., 2017). We have shown that the Avr4 protein triggers a biphasic ROS burst in *N. benthamiana:Cf-4*, while this biphasic ROS burst is eliminated in an *N. benthamiana:Cf-4 sobir1* knock-out line (Huang et al., 2021). To examine whether *Nb*SOBIR1 activation segment phosphorylation is crucial for mediating the Avr4-triggered ROS burst, we transiently expressed each *Nb*SOBIR1 mutant in the leaves of *N. benthamiana:Cf-4 sobir1* plants, after which the ROS production of discs taken from these leaves was monitored upon adding the Avr4 protein. Intriguingly, in contrast to the other four *Nb*SOBIR1 mutants and the positive control (*Nb*SOBIR1 WT), complementation with neither *Nb*SOBIR1 T522A nor the negative control (*Nb*SOBIR1 D482N) restored the Avr4-triggered ROS burst in *N. benthamiana:Cf-4 sobir1* knock-out line (Figure 1D).

Rapid and transient activation of MAPK cascades is another critical downstream event in the resistance of plants to pathogens (DeFalco and Zipfel, 2021; Yu et al., 2017). To determine whether Avr4/Cf-4-triggered MAPK activation in *N. benthamiana* also requires *Nb*SOBIR1 Thr522, we transiently co-expressed the mutants *NbSOBIR1 T522A* (and *NbSOBIR1 WT* and *D482N* as a positive and negative control, respectively) with *Avr4* in the leaves of an *N. benthamiana:Cf-4 sobir1* knock-out line, and subsequently detected possible MAPK activation by incubating western blots of a total protein extract with p42/p44-erk antibodies. Similar to the negative kinase-dead control but in contrast to the positive WT control, complementation with *Nb*SOBIR1 T522A failed to restore the Avr4/Cf- 4-induced MAPK activation in the *sobir1* knock-out line (Figure 1E).

Consistently, the analogous residues of *Nb*SOBIR1 Thr522 in tomato SOBIR1, which are *Sl*SOBIR1 Thr513 and *Sl*SOBIR1-like Thr526, gave the same phenotype after complementation with their Thr-to-Ala mutants in *N. benthamiana:Cf-4 sobir1* knock-out plants (Figure S2B-S2I). These residues are overall highly conserved in SOBIR1 from various plant species (Figure 1A and S2A), which suggests that this specific Thr residue might be crucial for the functionality of SOBIR1 in all plant species.

### SOBIR1 and BAK1 trans-phosphorylate each other *in vitro*

Thr522 is located at the activation segment of the kinase domain of *Nb*SOBIR1, and plays an important role in Cf-4/SOBIR1-initiated plant immunity. That led us to hypothesize this residue is crucial for the intrinsic kinase activity of SOBIR1, and thereby for its auto-phosphorylation. This SOBIR1 auto-phosphorylation represents step 1 in the model that was proposed by van der Burgh et al (2019), in which SOBIR1 and BAK1 act together in immune signaling. To test whether this is the case, we employed *in vitro* phosphorylation assays. The N-terminally GST-tagged cytoplasmic kinase domain from *Nb*SOBIR1, as well as this domain from various *Nb*SOBIR1 mutants, were expressed in *Escherichia coli*, which was followed by SDS-PAGE of the *E. coli* lysates, Coomassie brilliant blue (CBB) staining (for determining the accumulation levels of the various mutant kinase domains) and Pro-Q staining (to determine the phosphorylation state of the different kinase domains) (Taylor et al., 2013). Successful production of the various *Nb*SOBIR1 kinase domains was confirmed by western blotting, using SOBIR1 antibodies (Figure 2A). *Nb*SOBIR1 WT exhibited strong auto-phosphorylation activity, similar to its homolog in Arabidopsis (Leslie et al., 2010). Interestingly, when compared to *Nb*SOBIR1 WT, the auto-phosphorylation activity of *Nb*SOBIR1 T522A was completely abolished, similar to that of the kinase-dead D482N mutant, suggesting that indeed *Nb*SOBIR1 Thr522 is essential for the intrinsic kinase activity, and thereby the auto-phosphorylation, of *Nb*SOBIR1 (Figure 2A). In line with this observation for *Nb*SOBIR1 T522A, for *Sl*SOBIR1 T513A and *Sl*SOBIR1-like T526A also a loss of their intrinsic kinase activity was observed (Figure S4A and S4B). Altogether, these results demonstrate that this specific Thr residue is essential for SOBIR1 intrinsic kinase activity.

**Figure 2.**
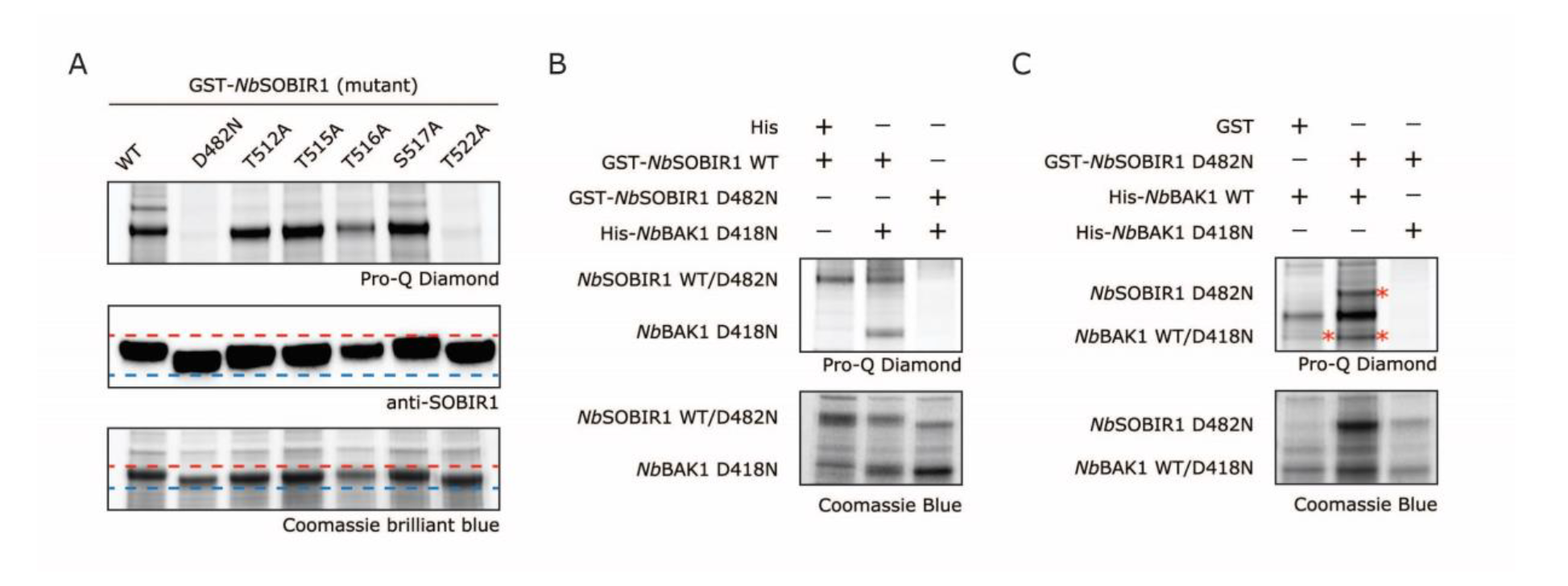
SOBIR1 and BAK1 Trans-Phosphorylate Each Other *In Vitro*. **(A)** Thr522 is required for the intrinsic kinase activity of *Nb*SOBIR1. The N-terminally GST-tagged cytoplasmic kinase domain of *Nb*SOBIR1, and its Ala substitution mutants, were produced in *E. coli*, in addition to kinase-dead D482N as a negative control. After SDS-PAGE of the *E. coli* lysates, the recombinant proteins were stained with Coomassie brilliant blue (bottom panel), whereas the phosphorylation status of the kinase domains was determined by performing a Pro-Q Diamond stain (top panel). Accumulation of the various *Nb*SOBIR1 kinase domains was detected by western blotting, using SOBIR1 antibodies (middle panel). The dotted lines are added in the middle and bottom panels to show the mobility shifts of the different SOBIR1 mutants, with the red dotted lines indicating the top of the highest protein bands, and the blue dotted lines indicating the base of the lowest protein bands. **(B)** *Nb*SOBIR1 WT directly phosphorylates kinase-dead *Nb*BAK1 D418N, and **(C)** *Nb*BAK1 WT directly phosphorylates kinase-dead *Nb*SOBIR1 D482N. The cytoplasmic kinase domains of *Nb*SOBIR1 WT and *Nb*SOBIR1 D482N were fused to a GST tag, whereas the cytoplasmic kinase domains of *Nb*BAK1 WT and its kinase-dead mutant (*Nb*BAK1 D418N), were fused to a His tag. Non-fused GST and His tags served as negative controls. After co-expressing the indicated recombinant proteins in *E. coli*, the proteins were separated by SDS-PAGE and stained with Coomassie brilliant blue, whereas their phosphorylation status was determined by performing a Pro-Q Diamond stain. Bands with the expected sizes are indicated with an asterisk. Experiments were repeated at least three times with similar results, and representative results are shown.

To elucidate whether SOBIR1 can directly phosphorylate BAK1, corresponding to the proposed SOBIR1 to BAK1 trans-phosphorylation step 2 in the model of van der Burgh et al (2019), we performed an *in vitro* phosphorylation assay. The GST-tagged *Nb*SOBIR1 WT cytoplasmic kinase domain, or its corresponding kinase-dead mutant, was co-expressed with the His-tagged cytoplasmic kinase domain of the *Nb*BAK1 kinase-dead mutant. The result showed that, first of all, the kinase-dead mutant of *Nb*BAK1, *Nb*BAK1 D418N, of which the conserved “RD” motif in the catalytic loop is changed into “RN”, did not have intrinsic auto-phosphorylation activity, as a Pro-Q stain was negative for this mutant when combined with *Nb*SOBIR1 D482N (Figure 2B). However, this mutant was properly phosphorylated by kinase-active *Nb*SOBIR1 WT, as visualized by a positive Pro-Q stain for *Nb*BAK1 D418N (Figure 2B). Similarly, the tomato homolog of *Nb*BAK1, *Sl*BAK1, was also directly phosphorylated by *Sl*SOBIR1 WT, as well as by *Sl*SOBIR1-like WT *in vitro*, as in both cases *Sl*BAK1 D418N was properly phosphorylated. Also here, this trans-phosphorylation fully depended on the intrinsic kinase activity of *Sl*SOBIR1 WT and *Sl*SOBIR1-like WT, as their corresponding “RN” kinase-dead mutants did not phosphorylate *Sl*BAK1 D418N (Figure S4C and S4D).

We next sought to determine whether BAK1 WT can directly phosphorylate SOBIR1, an event that corresponds to the proposed BAK1 to SOBIR1 trans-phosphorylation step 3 in the model of van der Burgh et al (2019). Earlier, we already showed that the strong auto-phosphorylation activity of *E. coli*-produced *Nb*SOBIR1 was eliminated in its “RN” kinase-dead mutant (Figure 2C). Therefore, we performed an additional *in vitro* phosphorylation assay by co-expressing *Nb*SOBIR1 D482N with either *Nb*BAK1 WT or its kinase-dead mutant. Importantly, *Nb*SOBIR1 D482N was phosphorylated when co-expressed with *Nb*BAK1 WT, but not when co-expressed with *Nb*BAK1 D418N, which demonstrates that indeed *Nb*SOBIR1 can be trans-phosphorylated by *Nb*BAK1 (Figure 2C). Consistently, *Sl*SOBIR1 D473N and *Sl*SOBIR1-like D486N were also directly phosphorylated by *Sl*BAK1 WT (Figure S4E and S4F). Notably, trans-phosphorylation of SOBIR1 by BAK1 also required intrinsic kinase activity of BAK1, as phosphorylation of kinase-dead SOBIR1 did not take place when co-expressed with the BAK1 kinase-dead mutant (Figure 2C, S4E and S4F).

### Tyr469, present in the kinase domain of *Nb*SOBIR1, is essential for mounting the Avr4/Cf-4-triggered HR and MAPK activation, but not required for ROS production and its intrinsic kinase activity

To determine whether particular Tyr residues that are present in the kinase domain of *Nb*SOBIR1 are specifically required for the Avr4/Cf-4-triggered HR, all eight Tyr residues present in the kinase domain of *Nb*SOBIR1 were selected to be studied (Figure 3A). Strikingly, most of these Tyr residues are highly conserved in SOBIR1 from many different plant species (Figure S5 and S6). We conducted a site-directed mutagenesis involving the substitution of each of the eight Tyr residues of *Nb*SOBIR1 by phenylalanine (Phe/F), which lacks the phosphorylatable hydroxyl group at the aromatic ring. Each *Nb*SOBIR1 mutant was co-expressed with *Avr4* in *N. benthamiana:Cf-4 sobir1* knock-out plants, including *WT* as a positive control and *D482N* as a negative control. Intriguingly, in contrast to the WT, but very similar to the kinase-dead mutant, transient expression of *NbSOBIR1 Y469F* failed to fully complement the Avr4-triggered HR in *N. benthamiana sobir1* knock-out plants (Figure 3B). Quantification of the HR intensity that was obtained upon complementation by red light imaging (Landeo Villanueva et al., 2021), showed that the differences in the HR intensity between *Nb*SOBIR1 WT and *Nb*SOBIR1 Y469F was significant (Figure 3C). Furthermore, the remaining Tyr mutants showed a similar complementation capacity in *N. benthamiana:Cf-4 sobir1* knock-out plants to the positive control (Figure 3C).

**Figure 3.**
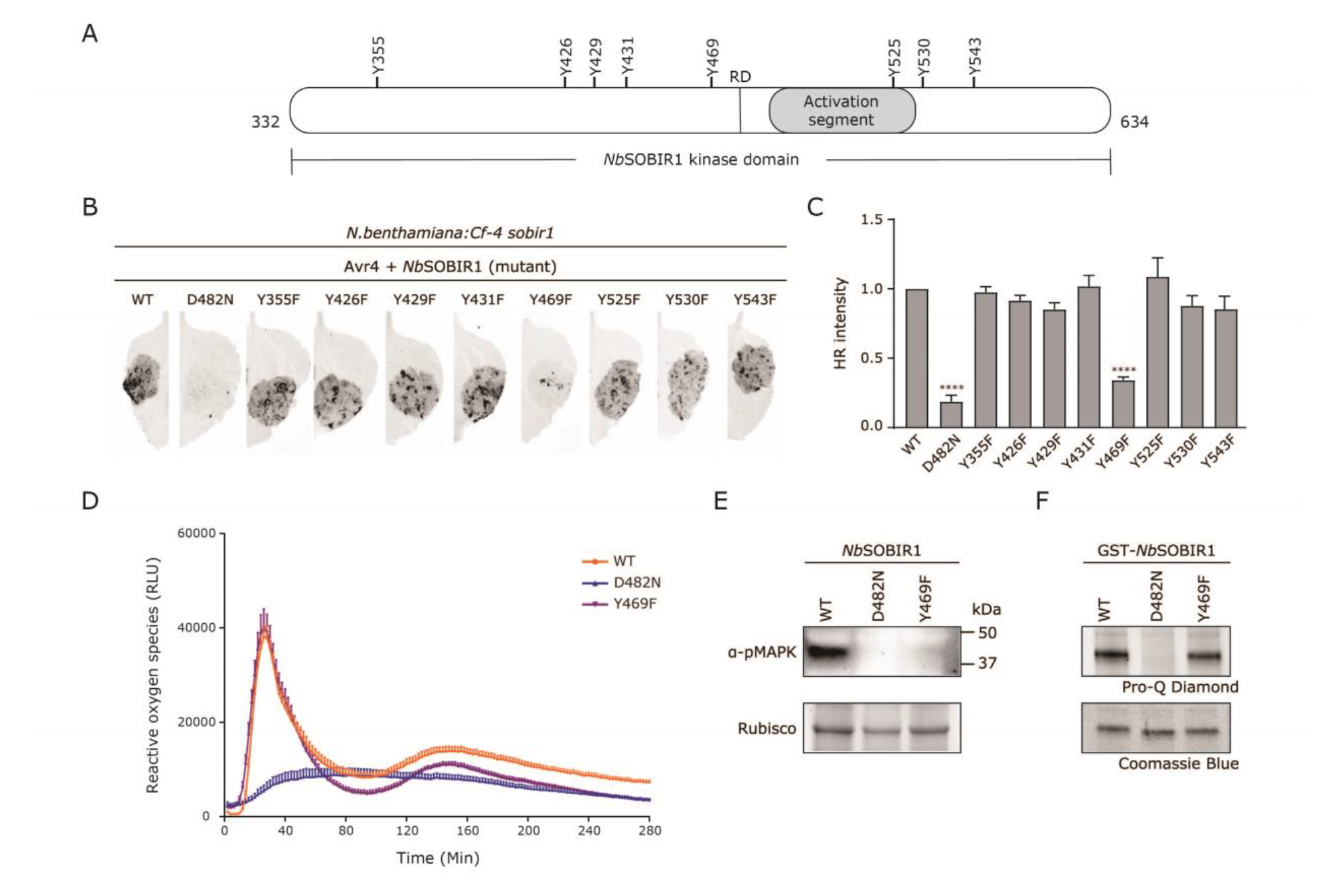
Tyr469, Present in the Kinase Domain of *Nb*SOBIR1, Is Crucial for the Avr4/Cf- 4/SOBIR1/BAK1-Triggered HR and MAPK Activation, but not for ROS Production and Intrinsic Kinase Activity. **(A)** Schematic diagram of the kinase domain of *Nb*SOBIR1, with the location of the activation segment, the RD motif, and all Tyr (Y) residues indicated. **(B** to **E)** Mutagenesis screen of all putative Tyr phosphorylation sites in *Nb*SOBIR1, to determine their importance in immune signaling by complementation. **(B)** All generated *Nb*SOBIR1 Tyr (Y)-to-Phe (F), the latter which is a mutant of Tyr that cannot be phosphorylated, mutants were transiently co-expressed with *Avr4* by agro-infiltration (both at an OD600 of 0.8) in the leaves of *N. benthamiana:Cf- 4 sobir1* knock-out plants, with WT as a positive control and kinase-dead D482N as a negative control. The presence of an HR was determined at 5 dpi, using the red-light imaging system (filter: 605/50; light: Green Epi Illumination; exposure time: 2 seconds). **(C)** Image Lab quantification of the intensity of the HR. Data shown are the average relative intensities of the HR + SEM (n≥6) that was triggered upon complementation with the various SOBIR1 mutants, when compared to complementation with WT *Nb*SOBIR1, of which the HR that was triggered was set to 1. Statistical significance was determined by a one-way ANOVA/Dunnett’s multiple comparison test, compared with *Nb*SOBIR1 WT. ****p < 0.0001. **(D)** The different Tyr mutants of *Nb*SOBIR1 were transiently expressed in leaves of the *N. benthamiana:Cf-4 sobir1* knock-out line, with WT as a positive control and D482N as a negative control. Leaf discs were taken from these plants at 24 hours after agro-infiltration, followed by adding 0.1 μM Avr4 protein and measuring ROS accumulation over time. ROS production is expressed as RLUs, and the data are represented as mean + SEM. All the tested *Nb*SOBIR1 Tyr mutants at least partially restored the Avr4/Cf-4-triggered ROS production in this complementation study, similar to *Nb*SOBIR1 WT. Only the results from *Nb*SOBIR1 WT, Y469F, and D482N are shown. (E) *NbSOBIR1 Y469F*, as well as *NbSOBIR1 WT* and *D482N*, were transiently co-expressed with *Avr4* in leaves of *N. benthamiana:Cf-4 sobir1* knock-out plants. Leaf samples were collected at 2 dpi and total protein extracts were subjected to immunoblotting with a p42/p44-erk antibody to determine the activation of downstream MAPKs by phosphorylation. (**F**) *Nb*SOBIR1 Y469F exhibits intrinsic kinase activity. N-terminally GST-tagged cytoplasmic kinase domains of *Nb*SOBIR1 WT, D482N and Y469F were produced in *E. coli*. After SDS-PAGE of the *E. coli* lysates, the recombinant proteins were stained with Coomassie brilliant blue (lower panel), whereas the phosphorylation status of the kinase domains was determined by performing a Pro-Q Diamond stain (upper panel). Experiments were repeated at least three times with similar results, and representative results are shown.

To investigate whether *Nb*SOBIR1 Tyr469 is also required for regulating the Avr4/Cf-4- triggered ROS burst, we monitored the ROS accumulation in the leaves of *N. benthamiana:Cf-4 sobir1* knock-out plants in which we transiently expressed the individual SOBIR1 mutants, upon adding Avr4 protein. Surprisingly, similar to *Nb*SOBIR1 WT and all other Tyr-to-Phe mutants, transient expression of *NbSOBIR1 Y469F* also partially restored the Avr4-triggered ROS burst in *N. benthamiana:Cf-4 sobir1* knock-out plants (Figure 3D). These results suggest that *Nb*SOBIR1 Tyr469 is not essential for the Avr4/Cf-4-triggered ROS burst.

To further explore the importance of *Nb*SOBIR1 Tyr469 in Avr4/Cf-4-mediated immune signaling, we determined the occurrence of Avr4/Cf-4-induced MAPK activation in *N. benthamiana:Cf-4 sobir1* knock-out plants, after co-expressing *NbSOBIR1 Y469F* with *Avr4*. *Nb*SOBIR1 WT was taken along as a positive control, and the kinase-dead mutant was taken along as a negative control. In contrast to the positive control, but similar to the negative control, co-expression of *NbSOBIR1 Y469F* with *Avr4* failed to restore the Avr4/Cf- 4-triggered MAPK activation in *N. benthamiana:Cf-4 sobir1* knock-out plants (Figure 3E). Again, the lack of MAPK activation was not correlated with its protein accumulation *in planta* (Figure S7).

Transient expression of *NbSOBIR1 Y469F* at least partially restored the Avr4-triggered ROS production in *N. benthamiana:Cf-4 sobir1* knock-out plants (Figure 3D), which led us to speculate that this Tyr-to-Phe mutant would still exhibit intrinsic kinase activity. As expected, the recombinant *Nb*SOBIR1 Y469F still showed strong auto-phosphorylation activity *in vitro* (Figure 3F). It is worth noting that we did not observe obvious differences between the staining intensities of the bands upon Pro-Q staining or between the protein mobilities of this mutant and *Nb*SOBIR1 WT (Figure 3F). Therefore, we conclude that *Nb*SOBIR1 Tyr469 plays no, or only a minor role, in determining the intrinsic kinase activity of *Nb*SOBIR1.

Consistently, the analogous residues of *Nb*SOBIR1 Tyr469 in tomato SOBIR1s, *Sl*SOBIR1 Tyr460 and *Sl*SOBIR1-like Tyr473, gave a similar phenotype after complementation with their Tyr to Phe mutants (Figure S6, S7 and S8). Taken together, these observations demonstrate that this particular conserved Tyr residue in the kinase domain of SOBIR1 plays an essential role in mediating specific Avr4/Cf-4-triggered plant immune responses.

### Members of *N. benthamiana* RLCK class VII subfamily 6, 7 and 8 appear all to be required for the Avr4/Cf-4/SOBIR1-triggered ROS burst

Increasing evidence has shown that RLCKs play an important role in regulating plant defense responses (DeFalco and Zipfel, 2021; Liang and Zhou, 2018; Lin et al., 2013). The genome of Arabidopsis contains 149 RLCK-encoding genes, which are divided into 17 classes (Shiu and Bleecker, 2001; Shiu et al., 2004). Some members from RLCK-VII, such as BIK1, PBL1, PBL13, RPM1-INDUCED PROTEIN KINASE (RIPK) and PBL27, have been well described (Li et al., 2021; Lin et al., 2015; Lu et al., 2010; Shinya et al., 2014; Zhang et al., 2010). The 46 members of RLCK-VII can be further classified into nine subfamilies, of which subfamilies 4, 5, 6, 7, and 8 were found to be involved in the ROS accumulation induced by different ExIPs (Luo et al., 2020; Pruitt et al., 2021; Rao et al., 2018). In addition to Arabidopsis, the Solanaceous plant *N. benthamiana* is another versatile experimental host plant (Bombarely et al., 2012; Goodin et al., 2008). In our previous studies, silencing of some individual RLCK-encoding genes in *N. benthamiana*, followed by monitoring the intensity of the Avr4/Cf-4-triggered HR, did not result in the identification of any promising individual RLCK candidate (van der Burgh et al., 2018). This could be the result of redundancy, as the family of RLCKs in *N. benthamiana* is large and thus multiple RLCK proteins might perform the same function (Goodin et al., 2008; Kourelis et al., 2019). Another reason might be that the selected candidates are only involved in regulating ROS production and/or MAPK activation, but not in the activation of the HR.

To search for RLCKs in Solanaceous plants, we, first of all, performed a phylogenetic analysis to identify Arabidopsis BIK1 homologs in Arabidopsis, tomato, and *N. benthamiana* (Figure S9A). All the RLCKs were further assigned to six subfamilies, which are referred to as subfamily 4, 5, 6, 7, 8, and 9, and are corresponding to the RLCK-VII subfamilies in Arabidopsis that were previously reported by Rao and colleagues (Figure S9B) (2018). To identify the RLCKs that play a role in cytoplasmic immune signaling downstream of the activated Avr4/Cf-4/SOBIR1/BAK1 complex, and to cope with the consequences of functional gene redundancy, we attempted to implement the CRISPR/Cas9 system in *N. benthamiana*:*Cf-4* to simultaneously target multiple RLCK-encoding genes belonging to the same subfamily (Table S1). *N. benthamiana* RLCK-VII-6 contains 14 members, which might be beyond the limit of the current multiplex CRISPR/Cas9 system (Figure S9B). Hence, the genes that are up-regulated or remain unchanged in their expression in *N. benthamiana:Cf-4* upon transient expression of *Avr4*, *NB-LRR PROTEIN REQUIRED FOR HR-ASSOCIATED CELL DEATH 1* (*NRC1*) *D481V* which triggers an elicitor-independent HR in *N. benthamiana*, and *COAT PROTEIN* (*CP*) which also triggers an HR in *N. benthamiana* when being overexpressed, were selected to be knocked out (Figure S10 and Table S1) (Gabriëls et al., 2007; Tameling et al., 2010).

We performed ROS burst assays on the transgenic plants of the T1 generation, in which all targeted *RLCK* members of each subfamily were anticipated to be knocked out, to identify the RLCK subfamilies that are required for the ROS production triggered upon activation of the Cf-4/SOBIR1 complex by Avr4. As we observed before, the Avr4 protein triggered a biphasic ROS burst in leaf discs obtained from *N. benthamiana:Cf-4* plants (Huang et al., 2021) (Figure 4). Strikingly, the intensity of the first, early phase of the ROS burst, which generally reaches its peak at around 20 minutes after the addition of the Avr4 protein, was only slightly reduced, whereas the second, more sustained phase of the ROS burst, generally reaching its peak around 140 minutes after Avr4 addition, was completely absent in all the independent *N. benthamiana:Cf-4* RLCK-VII-6 knock-out lines, referred to as *rlck-vii-6* (Figure 4C). This observation indicates that members of RLCK-VII-6 probably play an important role in regulating the Avr4-triggered ROS burst in Cf-4-transgenic *N. benthamiana*, especially concerning the second, more sustained phase of the ROS burst. Intriguingly, the Avr4-induced ROS production was overall strongly compromised in all the *rlck-vii-7* knock-out lines, whereas two of the *rlck-vii-8* knock-out lines showed a similar reduction of the overall ROS burst when treated with Avr4 (Figure 4D and 4E). These findings suggest that also the members of RLCK-VII-7 and −8 might be redundantly required for the Avr4/Cf-4-triggered ROS burst.

**Figure 4.**
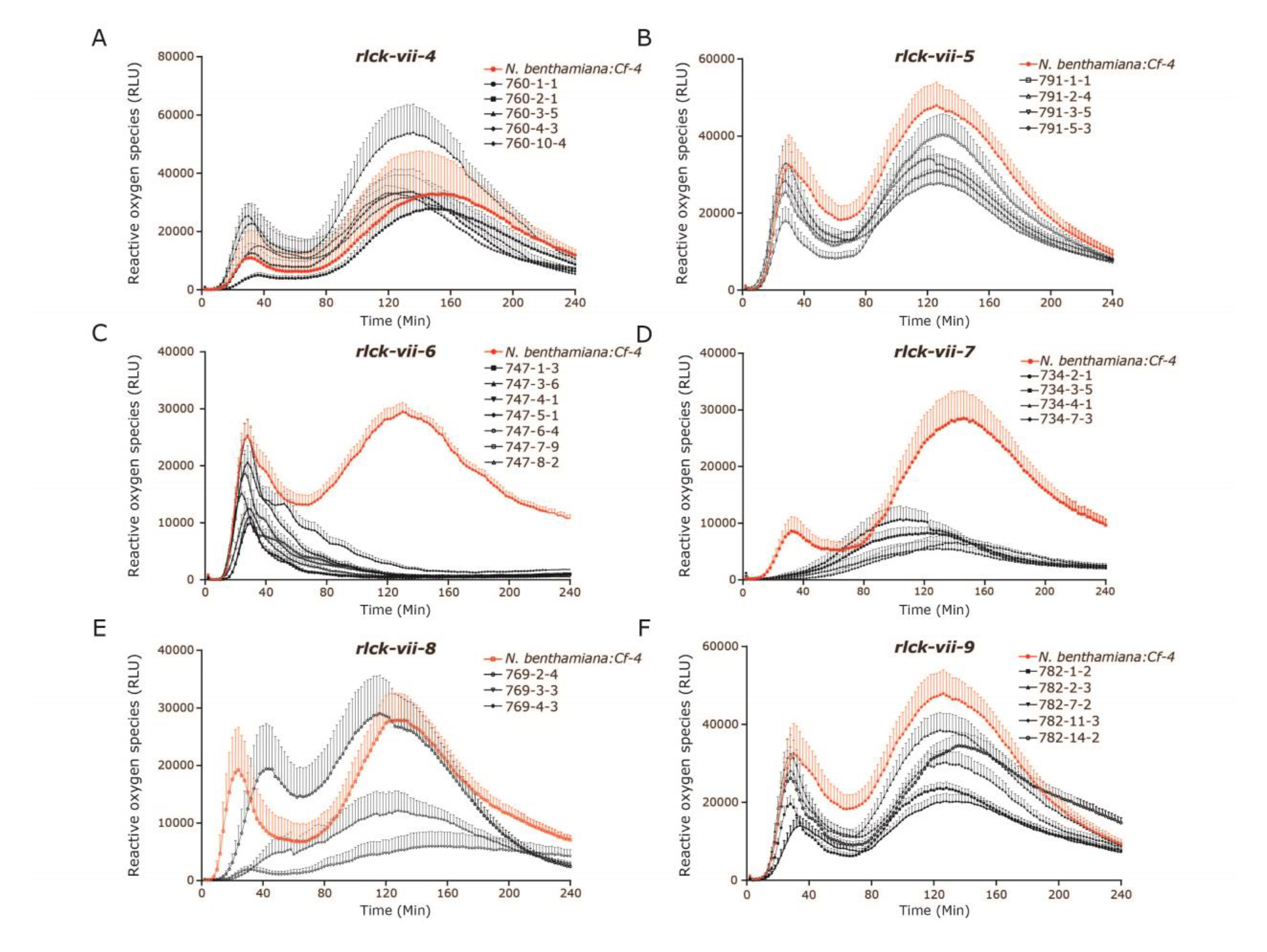
Various Subfamilies from RLCK Class VII Play an Important Role in the Avr4/Cf- 4/SOBIR1-Triggered ROS Burst in *N. benthamiana:Cf-4*. Selected members from each RLCK class VII subfamily, being 4, 5, 6, 7, 8 and 9, were targeted for knock-out in *N. benthamiana:Cf-4* by CRISPR/Cas technology. Subsequently, ROS accumulation induced upon Avr4 protein treatment of leaf discs obtained from five individual *rlck-vii-4* transformants **(A)**, four *rlck-vii-5* transformants **(B)**, seven *rlck-vii-6* transformants **(C)**, four *rlck-vii-7* transformants **(D)**, three *rlck-vii-8* transformants **(E)** and five *rlck-vii-9* transformants **(F)**, was determined. For this, leaf discs were treated with 0.1 μM Avr4 protein, and the generation of ROS was monitored. Note that all transformants were tested in the T1 generation. ROS production is expressed as RLUs, and the data are represented as mean + SEM (n≥6). The ROS profiles of the positive control (*N. benthamiana:Cf- 4*), included in all assays, are indicated in red in all the line charts. Similar results were obtained in replicates and data from one representative experiment are shown.

To further explore whether these RLCK-VII subfamilies in *N. benthamiana* also play a role in LRR-RLK-triggered immune signaling, we treated all the knock-out lines with the flg22 peptide to activate the endogenous FLS2 receptor of *N. benthamiana*. *N. benthamiana:Cf-4* was used as a positive control, which showed a rapid and typical monophasic FLS2- mediated ROS burst upon flg22 peptide treatment, reaching its peak at about 25 minutes after the addition of the peptide (Huang et al., 2021) (Figure S11). Similar to our previous observation when performing ROS assays with the pure Avr4 protein, the flg22-induced ROS accumulation was dampened to different levels in all the *rlck-vii-6* knock-out lines and was almost eliminated in some of the *rlck-vii-7* and *rlck-vii-8* knock-out lines (Figure S11C, S11D and S11E). Surprisingly, several *rlck-vii-4* knock-out lines displayed an increased ROS burst when challenged with flg22, which suggests that members of RLCK-VII-4 may negatively regulate the flg22/FLS2-triggered ROS production (Figure S11A). As observed previously upon treatment with Avr4, the remaining *rlck-vii* knock-out lines did not show obvious phenotypes in these flg22 assays. Our observations indicate that, in addition to a role in Avr4/Cf-4/SOBIR1-triggered immune signaling, the members of the RLCK-VII-6, −7 and −8 might also play a positive role in FLS2-mediated immune signaling.

### Members of *N. benthamiana* RLCK-VII-7 appear to play a crucial role in the Avr4/Cf-4/SOBIR1-triggered HR

These *rlck* knock-out plants were then subjected to HR assays to further identify the RLCK subfamilies that are important for Avr4/Cf-4-triggered HR activation. Infiltration of pure Avr4 protein in leaves of *N. benthamiana:Cf-4* induces HR activation, which can be detected and quantified by red light imaging (Landeo Villanueva et al., 2021). This HR intensity was significantly attenuated in all the *rlck-vii-7* knock-out lines, but not in the other *rlck-vii* lines (Figure 5), demonstrating that in addition to their role in ROS production, members from RLCK-VII-7 might also be pivotal for the Avr4/Cf-4-triggered HR.

**Figure 5.**
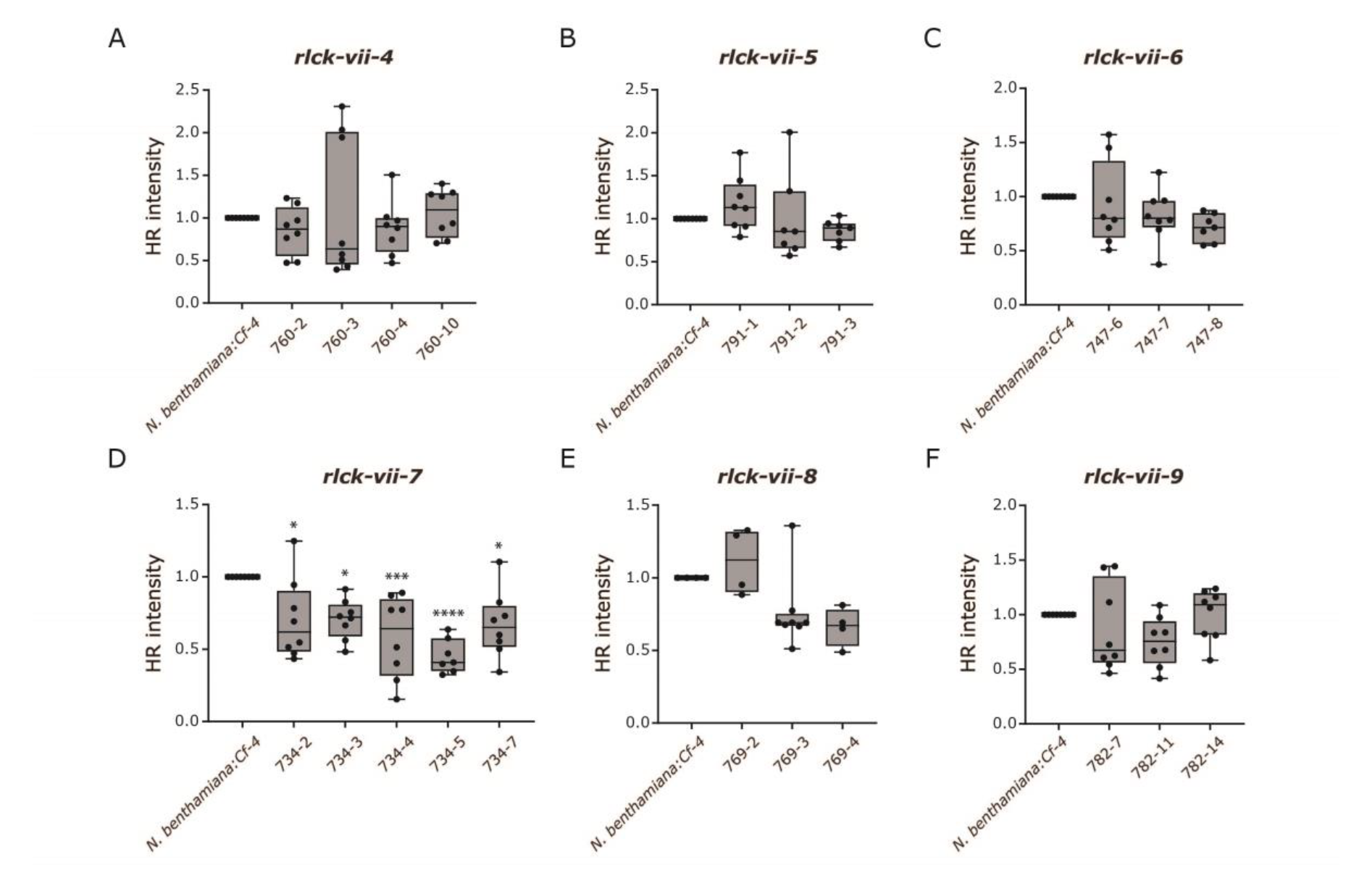
RLCK-VII-7 Is Required for the Avr4/Cf-4-Triggered HR in *N. benthamiana:Cf-4*. A solution of 5 μM pure Avr4 protein was infiltrated in leaves of the *N. benthamiana:Cf-4 rlck-vii-4* **(A)**, *rlck-vii-5* **(B)**, *rlck-vii-6* **(C)**, *rlck-vii-7* **(D)**, *rlck-vii-8* **(E)** and *rlck-vii-9* **(F)** transformants, and the Avr4/Cf-4-triggered HR was subsequently imaged using the ChemiDoc and quantified using Image Lab, at 2 dpi. All quantifications are shown as dots (n≥6) and the means as lines. Statistical analysis was performed with ANOVA/Dunnett’s multiple comparison test, compared with *N. benthamiana:Cf-4.* *p < 0.05; ***p < 0.001; ****p < 0.0001. Experiments were repeated at least three times with similar results, and representative results are shown.

### Members of RLCK-VII-6 positively regulate the production of ROS triggered by multiple ExIPs in *N. benthamiana,* but are not required for Avr4/Cf-4-triggered MAPK and HR activation

To knock out eight members from RLCK-VII-6 simultaneously, nine single-guide RNAs (sgRNAs) were designed (Table S1). The genotypes of four T1 knock-out plants, which originated from two independent transgene-free lines in which *rlck-vii-6* was targeted, were determined and the result was described by Stuttmann et al. (2021). By amplifying and sequencing the targeted gene regions, three independent homozygous knock-out lines, named 747-3-6-2, 747-3-6-7 and 747-7-9-6, in which all eight candidate genes were mutated, were identified (Figure 6A). Notably, these knock-out lines did not exhibit any significant changes in overall morphology, when compared to the original *N. benthamiana:Cf-4* plants in which the mutagenesis was performed (Figure 6B).

**Figure 6.**
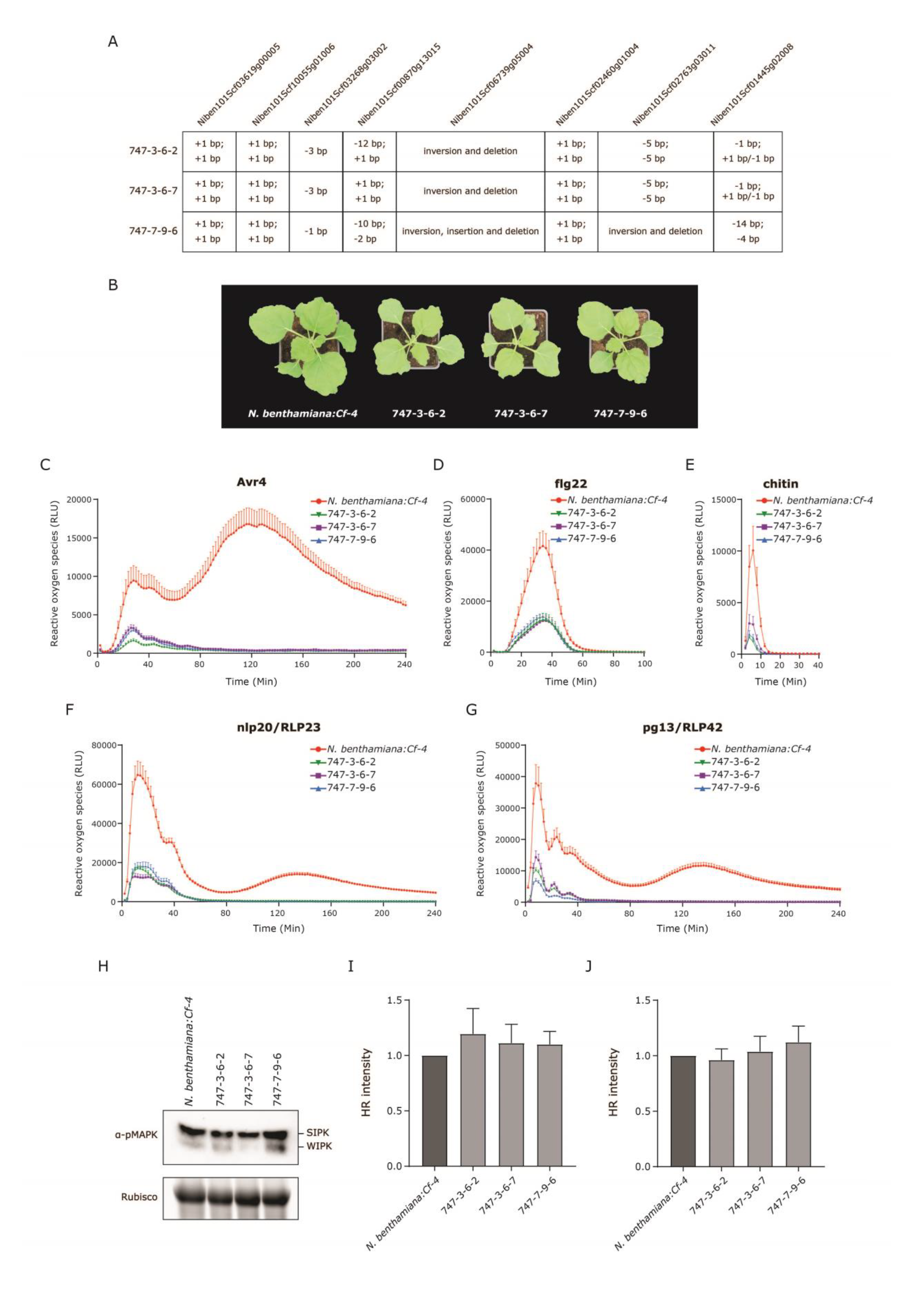
Members of *N. benthamiana* RLCK-VII-6 Are Required for ROS Production Triggered by Multiple ExIPs, But Dispensable for the Avr4/Cf-4-Triggered MAPK and HR Activation. **(A)** Overview of the types of mutations present in all the RLCK-VII-6 members in the three homozygous *N. benthamiana:Cf-4* knock-out lines. Note that two single-guide RNAs (sgRNAs) were designed to target each gene, except for *Niben101Scf03268g03002*, which was only targeted by one sgRNA (Table S1). **(B)** Morphological phenotypes of *N. benthamiana:Cf-4* and the three independent *rlck-vii-6* knock-out lines. All plants were grown in soil under the same conditions, at the same time and were photographed when they were four to five weeks old. Leaf discs obtained from three independent *rlck-vii-6* homozygous T2 knock-out lines, as well as from *N. benthamiana:Cf-4* (the positive control), were treated with 0.1 μM pure Avr4 protein **(C)**, 0.1 μM flg22 **(D)**, or with 10 μM chitohexaose **(E)**, and ROS production was monitored. Leaf discs taken from the independent *N. benthamiana:Cf-4 rlck-vii-6* lines, as well as from *N. benthamiana:Cf-4*, transiently expressing either *RLP23* (F) or *RLP42* (G), were treated with the corresponding elicitors at a 1 μM concentration and the accumulation of ROS was monitored. ROS production is expressed as RLUs, and the data are represented as mean + SEM (n≥6). The ROS profiles of the positive control (*N. benthamiana:Cf-4*) that was included in all the assays are indicated in red in all the line charts. **(H)** The three independent *rlck-vii-6* homozygous T2 knock-out lines do not exhibit an altered Avr4/Cf-4-triggered MAPK activation when compared to *N. benthamiana:Cf-4* plants. A solution of 5 μM pure Avr4 protein was infiltrated in the leaves of knock-out plants, as well as in *N. benthamiana:Cf-4* plants. Leaf samples were taken 15 min later and total protein extracts were run on SDS gel and subjected to immunoblotting employing a p42/p44-erk antibody specifically detecting MAPKs that are activated by phosphorylation (α-pMAPK). The intensity of the Rubisco band in the stain-free gel indicates equal loading. Experiments were repeated at least three times and similar results were obtained. Representative pictures are shown. **(I** and **J)** The intensity of the Avr4/Cf-4- triggered HR in leaves of the three independent *rlck-vii-6* knock-out plants is similar when compared to *N. benthamiana:Cf-4*. **(I)** A solution of 5 μM pure Avr4 protein was infiltrated in leaves of the different plants, which were subsequently imaged using the ChemiDoc, with the RFP channel, at 2 days post infiltration (dpi). (J) *Avr4* was transiently expressed in leaves of the different plants (at an OD600 of 0.5), which were subsequently imaged using the ChemiDoc at 4 dpi. The intensity of the HR was quantified by Image Lab. Statistical significance was determined by a one-way ANOVA/Dunnett’s multiple comparison test, compared with *N. benthamiana:Cf-4*. Data are represented as mean + SEM. Experiments were repeated at least three times with similar results, and representative results are shown.

Both the Avr4- and flg22-induced ROS burst was attenuated in the *rlck-vii-6* knock-out plants from the T1 generation (Figure 4C and S11C). To further verify these results, leaf discs taken from the homozygous knock-out lines were again treated with either the Avr4 protein or flg22 peptide, followed by monitoring ROS accumulation, with *N. benthamiana:Cf-4* plants taken along as a positive control. Consistent with the aforementioned results, ROS production triggered by the Avr4 protein and flg22 peptide was strongly compromised when compared to *N. benthamiana:Cf-4*, thereby confirming that the members of RLCK-VII-6 are indeed required for both Cf-4/SOBIR1- and FLS2- triggered immune signaling (Figure 6C and 6D). The importance of this RLCK-VII subfamily in positively regulating ROS accumulation led us to determine possible changes in the ROS burst induced by other ExIPs, such as chitin, in the 747-3-6-2, 747-3-6-7 and 747-7-9-6 lines. Strikingly, these three independent knock-out lines also exhibited an obvious reduction in chitin-triggered ROS production, when compared to *N. benthamiana:Cf-4* (Figure 6E).

The Arabidopsis LRR-RLPs RLP23 and RLP42 perceive a conserved 20-amino-acid peptide from necrosis and ethylene-inducing peptide 1 (NEP1)-like proteins (nlp20) and *Botrytis cinerea* endo-polygalacturonases (PGs and their derived peptide, pg13), respectively (Albert et al., 2015; Zhang et al., 2021; Zhang et al., 2014). Similar to Cf-4, both RLP23 and RLP42 require SOBIR1 and BAK1 to initiate immune signaling upon recognition of their matching elicitor (Albert et al., 2015; Liebrand et al., 2013; Zhang et al., 2021; Zhang et al., 2014). Interestingly, although *RLP23* and *RLP42* are Arabidopsis genes, *N. benthamiana* plants that overexpress either *RLP23* or *RLP42* show sensitivity to their corresponding ExIPs. This suggests that RLP23 and RLP42, like Cf-4, are able to employ the endogenous *N. benthamiana* SOBIR1 and BAK1 proteins and additional, probably functionally conserved, downstream partners to activate immune signaling in response to treatment with nlp20 and pg13, respectively (Albert et al., 2015; Zhang et al., 2021). To decipher whether the *N. benthamiana* RLCK-VII-6 family members also play a role downstream of RLP23 and RLP42, we transiently overexpressed them in the different *rlck-vii-6* knock-out lines, as well as in *N. benthamiana:Cf-4*, and collected leaf discs of the infiltrated area at 24 hours after infiltration. Subsequently, after adding the corresponding ligands, we monitored ROS production. Strikingly, similar to the situation with Avr4/Cf-4, both the nlp20/RLP23- and pg13/RLP42-mediated ROS burst in *N. benthamiana* plants was biphasic, of which the first burst was rapid with a relatively high amplitude, whereas the second burst was sustained, with relatively low amplitude (Figure 6F and 6G). Both the nlp20/RLP23- and the pg13/RLP42-triggered ROS production was significantly impaired in all *rlck-vii-6* knock-out plants, when compared to *N. benthamiana:Cf-4*. This was especially the case for the second burst, which was completely abolished (Figure 6F and 6G). Collectively, these results indicate that the members of RLCK-VII-6 are required for regulating the ROS production triggered by a broad spectrum of ExIPs.

In addition to ROS production, the activation of a downstream MAPK cascade is another key signaling event of ExTI, which generally can be detected within minutes upon ExIP recognition (Yu et al., 2017; Zhang et al., 2018). We observed that infiltration of the Avr4 protein in leaves of *N. benthamiana:Cf-4* induces the activation of a MAPK cascade within five minutes (Figure S12). To explore whether this Avr4/Cf-4-triggered MAPK activation is affected in the *rlck-vii-6* knock-out plants, we infiltrated pure Avr4 protein in leaves of the different knock-out lines as well as in leaves of *N. benthamiana:Cf-4* plants. The leaf samples were harvested at 15 min after Avr4 infiltration, followed by performing a western blot assay revealing MAPK activation. WOUND-INDUCED PROTEIN KINASE (WIPK) and SA-INDUCED PROTEIN KINASE (SIPK) are two MAPKs that are activated as a result of phosphorylation in *N. benthamiana* (Sharma et al., 2003) and that can be detected by specific antibodies upon their activation. Intriguingly, no obvious changes in MAPK activation in the three knock-out lines were observed at the tested time point when compared to *N. benthamiana:Cf-4*, suggesting that while the RLCK-VII-6 members are important for the ROS burst, these are not essential for the Avr4/Cf-4-triggered MAPK activation (Figure 6H).

Unlike some ExIPs such as flg22, elf18 or chitin that only induce weak ExTI responses in plants, Avr4 triggers a typical HR in leaves of *N. benthamiana:Cf-4* and MM-Cf-4 tomato (Cai et al., 2001; de Wit, 2016; Gabriëls et al., 2007; Gabriëls et al., 2006). To examine whether members of RLCK-VII-6 actually contribute to the Cf-4/Avr4-triggered HR in *N. benthamiana*, we infiltrated either pure Avr4 protein or *Agrobacterium tumefaciens* (Agrobacterium) harboring the binary constitutive *Avr4* expression construct in leaves of *rlck-vii-6* knock-out lines and of *N. benthamiana:Cf-4* as a control. Neither the Avr4 protein infiltration nor the agro-infiltration revealed significant changes in the capacity to mount an Avr4/Cf-4-triggered HR of the knock-out plants when compared to *N. benthamiana:Cf- 4* (Figure 6I and 6J). Altogether, these results demonstrate that the members of RLCK-VII-6 play no, or only a minor role, in regulating the Avr4/Cf-4-triggered MAPK and HR activation, whereas they are essential for Avr4/Cf-4-triggered ROS accumulation.

Of note, all these three *N. benthamiana:Cf-4 rlck-vii-6* knock-out lines show similar phenotypes, although some mutations might not inactivate the targeted RLCKs. Accordingly, these genes either do not contribute to the tested immune responses, or the mutations do actually inactivate them.

### Members of RLCK-VII-7 positively regulate the production of ROS triggered by multiple ExIPs and HR activation triggered by Avr4/Cf-4, but are not required for Avr4/Cf-4-triggered MAPK activation in *N. benthamiana*

Nine sgRNAs were designed to target seven genes of RLCK-VII-7 (Table S1), and two independent homozygous knock-out lines were selected by amplifying and sequencing the target gene regions, named 734-3-5-5-1 and 734-5-3-2-3 (Figure 7A). Of note, these knock-out plants did not display any obvious changes in plant overall morphology, when compared to *N. benthamiana:Cf-4* (Figure 7B).

**Figure 7.**
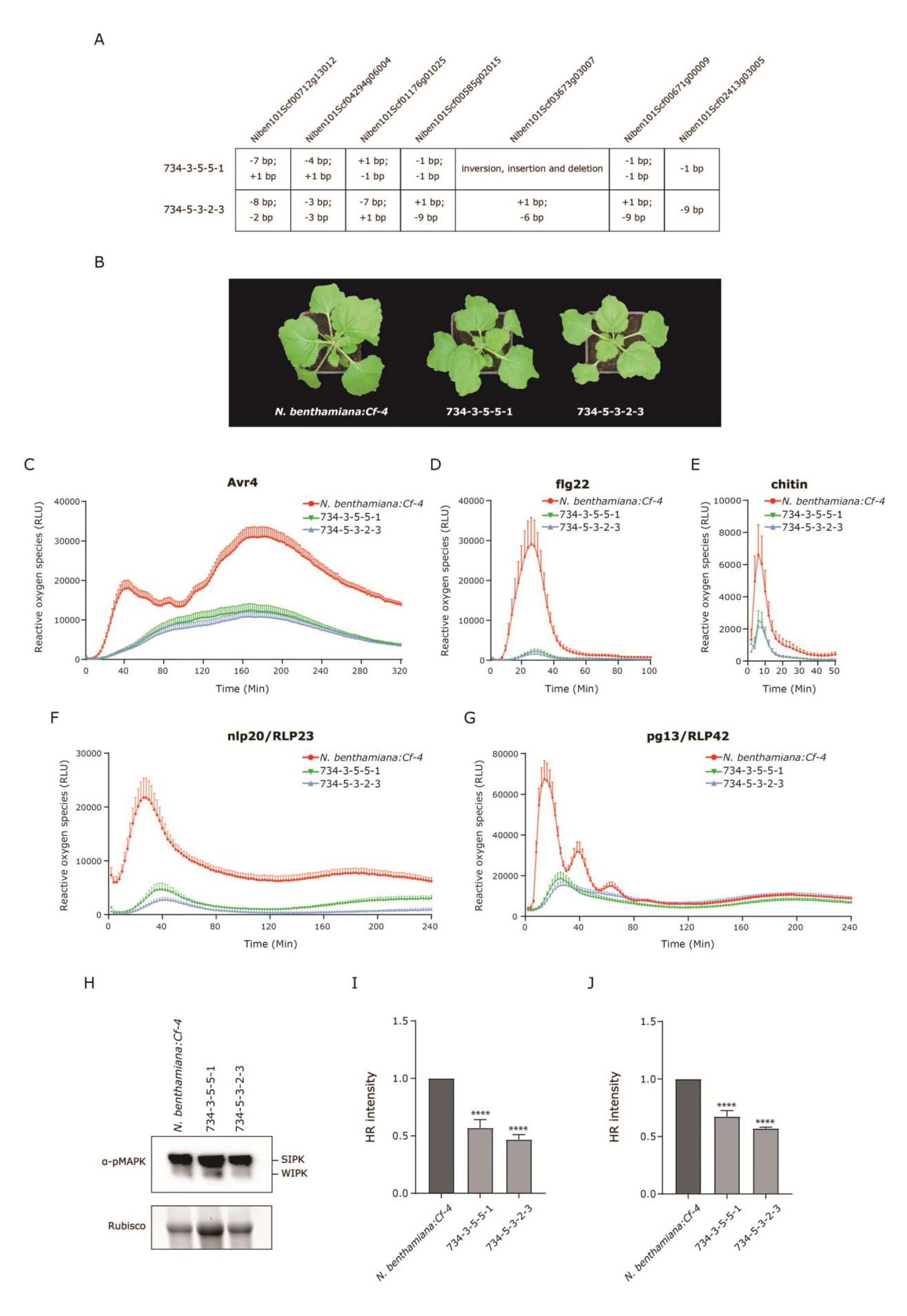
Members of RLCK-VII-7 Are Pivotal for ROS Production Triggered by Multiple ExIPs and for Avr4/Cf-4-Induced HR, But Are Dispensable for the Avr4/Cf-4-Triggered MAPK Activation. **(A)** Overview of the types of mutations present in all the RLCK-VII-7 members in the two independent homozygous *N. benthamiana:Cf-4* knock-out lines, 734-3-5-5-1 and 734-5- 3-2-3. Note that two sgRNAs were designed to target each gene, except for *Niben101Scf02413g03005*, which was only targeted by one sgRNA. **(B)** Morphological phenotypes of *N. benthamiana:Cf-4* and the two independent *rlck-vii-7* knock-out lines. All plants were grown in soil under the same conditions and were photographed when they were four to five weeks old. ROS production triggered by Avr4 **(C)**, flg22 **(D)**, chitin **(E)**, nlp20/RLP23 **(F)**, and pg13/RLP42 **(G)** in *N. benthamiana:Cf-4* (the positive control) and two independent *N. benthamiana:Cf-4 rlck-vii-7* knock-out plants. ROS production is expressed as RLUs, and the data are represented as mean + SEM (n≥6). The ROS profiles of the positive control (*N. benthamiana:Cf-4*) that was included in all the assays are indicated in red in all the line charts. **(H)** Avr4/Cf-4-triggered MAPK activation is not altered in *rlck-vii-7* knock-out plants, when compared to that in *N. benthamiana:Cf-4* plants. Pure Avr4 protein (5 μM) was infiltrated in the leaves of knock-out and *N. benthamiana:Cf-4* plants. Leaf samples were collected 15 min later, followed by total protein extraction and immunoblotting using a p42/p44-erk antibody (α-pMAPK). The intensity of the Rubisco band in the stain-free gel serves as a loading control. Similar results were obtained at least three times and representative images are shown. **(I** and **J)** The HR intensity triggered by Avr4/Cf-4 in leaves of two independent *rlck-vii-7* knock-out lines is significantly reduced, when compared to that in *N. benthamiana:Cf-4* plants. **(I)** Pure Avr4 protein (5 μM) was infiltrated in leaves of knock-out plants, as well as *N. benthamiana:Cf- 4* plants. The HR was imaged by the Chemidoc and the intensity was quantified by Image Lab at 2 dpi, as described above. **(J)** Agrobacterium harboring *Avr4* was infiltrated in the leaves of *N. benthamiana:Cf-4* and two knock-out lines (at an OD600 of 0.5). Avr4/Cf-4-induced HR was then detected and quantified at 4 dpi, as described above. Statistical significance was determined by a one-way ANOVA/Dunnett’s multiple comparison test, compared with *N. benthamiana:Cf-4*, ****p < 0.0001. Data are represented as mean + SEM. Experiments were repeated at least three times with similar results, and representative results are shown.

The knock-out plants of *rlck-vii-7* of the T1 generation already showed impaired Avr4/Cf- 4- and flg22/FLS2-induced ROS accumulation (Figure 4D and S11D). To validate these results and to further elucidate the importance of this particular subfamily in plant immunity, Avr4-, flg22-, chitin-, nlp20/RLP23-, and pg13/RLP42-triggered ROS production in the *N. benthamiana:Cf-4* control and in two selected *rlck-vii-7* knock-out lines was measured. Consistently, the marked decrease in ROS accumulation was again observed in the knock-out plants after the addition of Avr4 protein and flg22 peptide (Figure 7C and 7D), which confirms that members of RLCK-VII-7 indeed play a crucial role in both Avr4/Cf- 4- and flg22/FLS2-mediated immune signaling. Interestingly, unlike the *rlck-vii-6* knock-out plants, in which the first peak of the ROS burst induced by Avr4 was severely dampened and the second ROS burst was eliminated (Figure 6C), *rlck-vii-7* knock-out plants displayed an overall reduction of the Avr4/Cf-4-triggered ROS burst, and instead of the typical biphasic ROS, only a weak and sustained ROS burst was observed (Figure 7C). These observations point to the different roles that RLCK-VII-6 and RLCK-VII-7 have in regulating the Avr4/Cf-4-stimulated ROS production. Moreover, the ROS burst induced by chitin, nlp20/RLP23, and pg13/RLP42 was also strongly impaired in the *rlck-vii-7* knock-out plants when compared to *N. benthamiana:Cf-4* (Figure 7E, 7F and 7G). Therefore, in addition to members of RLCK-VII-6, members of RLCK-VII-7 also play a crucial role in regulating the production of ROS induced by a broad spectrum of ExIPs.

We next asked whether this particular subfamily is also involved in Avr4/Cf-4-mediated MAPK activation. A solution of pure Avr4 protein was infiltrated in leaves of *N. benthamiana:Cf-4* and two independent knock-out lines, after which leaf samples were then collected 15 min later, followed by total protein extraction, SDS-PAGE and western blotting. Notably, the two knock-out lines did not exhibit altered MAPK activation at the tested time point when compared to *N. benthamiana:Cf-4* (Figure 7H), suggesting that members of RLCK-VII-7 are not required for Avr4/Cf-4-triggered MAPK activation in *N. benthamiana*.

We observed that all *rlck-vii-7* knock-out plants displayed a significant reduction in the Avr4/Cf-4-triggered HR (Figure 5D). To confirm this result, the selected homozygous knock-out lines were subjected to an HR assay. Either pure Avr4 protein or Agrobacterium harboring an *Avr4* overexpression construct was infiltrated in leaves of the *N. benthamiana:Cf-4* control and two independent knock-out lines. Again, we observed that the intensity of the HR was strongly reduced in *N. benthamiana*, upon both treatments, when members of RLCK-VII-7 were knocked out (Figure 7I and 7J). Taken together, these results indicate that members of RLCK-VII-7 positively regulate ROS production mediated by Avr4, flg22, chitin, nlp20/RLP23, and pg13/RLP42 in *N. benthamiana*. Furthermore, although they seem not to be required for Avr4/Cf-4-triggered MAPK activation, the RLCK-VII-7 members indeed play an essential role in the Avr4/Cf-4-triggered HR.

It is worth noting that two *N. benthamiana:Cf-4 rlck-vii-7* knock-out lines show similar phenotypes, despite the fact that some targeted RLCKs from this subfamily might not be actually knocked out. This observation implies that these RLCKs either do not play a role in regulating plant immunity in *N. benthamiana* plants, or they are indeed non-functional anymore due to the mutations.

### Members of RLCK-VII-8 positively regulate the production of ROS triggered by multiple ExIPs, but are dispensable for Avr/Cf-4-triggered MAPK and HR activation in *N. benthamiana*

Since the members of RLCK-VII-8 showed a possible importance in regulating Avr4/Cf-4- and flg22/FLS2-mediated ROS accumulation in *N. benthamiana*, their homozygous knock-out lines were selected. Nine genes were targeted by an array of eight different sgRNAs (Table S1), and Sanger-sequencing of the targeted gene regions led to the identification of two independent homozygous knock-out lines, named 769-4-4-20-3 and 769-4-4-20-8 (Figure 8A). It is worth noting that knocking out RLCK-VII-8 members did not alter the overall plant morphology, when compared to *N. benthamiana:Cf-4* (Figure 8B).

**Figure 8.**
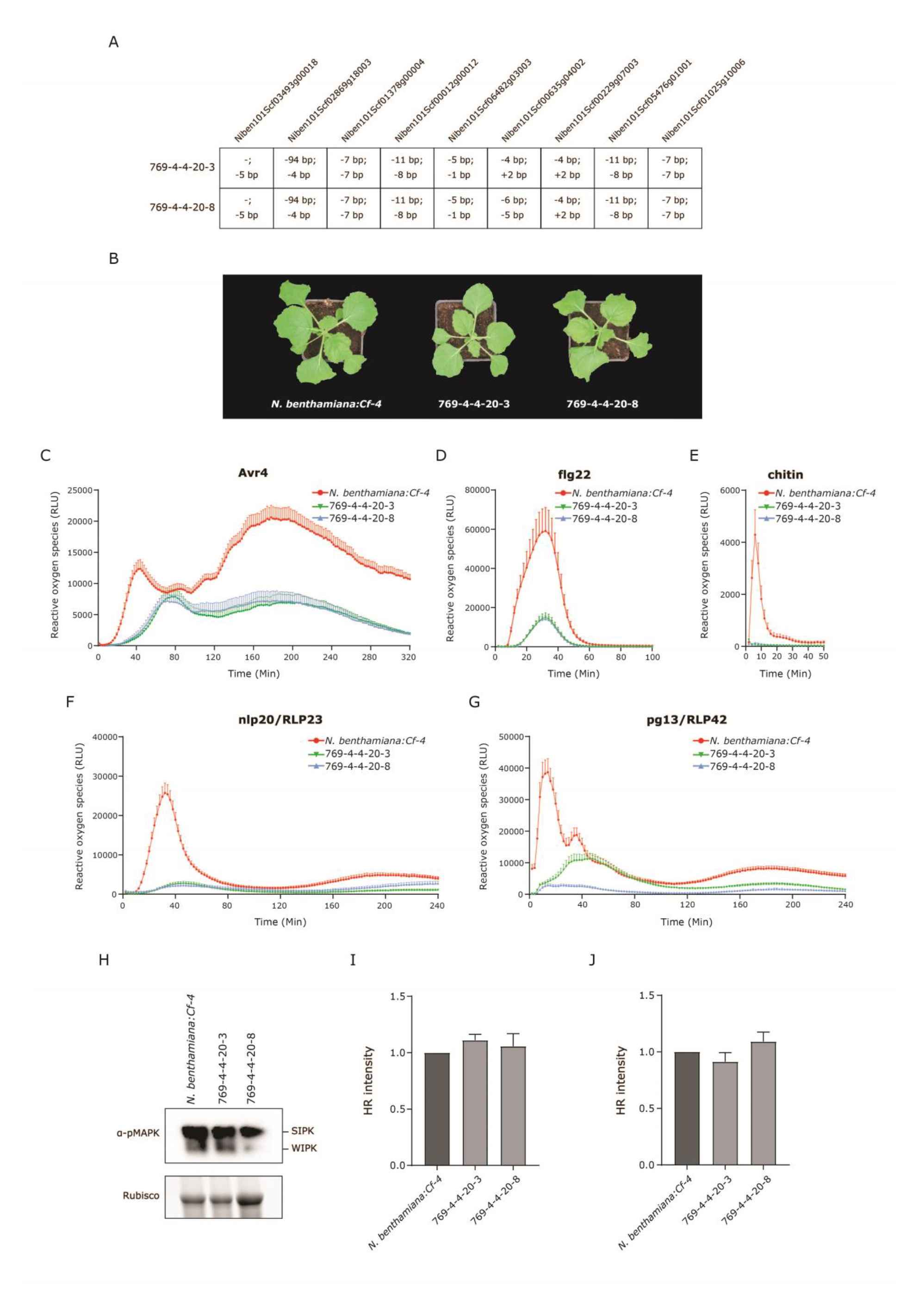
Members of *N. benthamiana* RLCK-VII-8 Play an Essential Role in ROS Production Triggered by Multiple ExIPs, But not in the Avr4/Cf-4-Triggered MAPK and HR Activation. **(A)** Overview of the types of mutations present in all the RLCK-VII-8 members in the two independent homozygous *N. benthamiana:Cf-4* knock-out lines, 769-4-4-20-3 and 769-4-4-20-8. Note that two sgRNAs were designed to target each gene. **(B)** Morphological phenotypes of *N. benthamiana:Cf-4* and the two independent *rlck-vii-8* knock-out lines. All plants were grown in soil under the same conditions and were photographed when they were four to five weeks old. Avr4- **(C)**, flg22- **(D)**, chitin- **(E)**, nlp20/RLP23- **(F)**, and pg13/RLP42- **(G)** stimulated ROS accumulation in two independent *N. benthamiana:Cf-4 rlck-vii-8* knock-out plants, as well as in *N. benthamiana:Cf- 4* (the positive control). ROS production is expressed as RLUs, and the data are represented as mean + SEM (n≥6). The ROS profiles of the positive control (*N. benthamiana:Cf-4*) that was included in all the assays are indicated in red in all the line charts. **(H)** Avr4/Cf-4-triggered MAPK activation in *rlck-vii-8* knock-out plants is similar to that in *N. benthamiana:Cf-4* plants. A solution of 5 μM pure Avr4 protein was infiltrated in the leaves of knock-out and *N. benthamiana:Cf-4* plants (the positive control). Leaf samples were harvested 15 min later, and total protein extracts were run on SDS gel and subjected to immunoblotting using a p42/p44-erk antibody (α-pMAPK). The intensity of the Rubisco band in the stain-free gel serves as a loading control. **(I** and **J)** The two independent *rlck-vii-8* knock-out lines do not exhibit altered Avr4/Cf-4-triggered HR when compared to *N. benthamiana:Cf-4* plants. Pure Avr4 protein (**I**) or Agrobacterium harboring *Avr4* (J) was infiltrated in leaves of *N. benthamiana:Cf-4* and two independent knock-out lines, which were then imaged by Chemidoc at 2dpi and 4 dpi, respectively. The HR intensity was quantified and analyzed as described above. Experiments were performed in triplicate with similar results.

Leaf disks of *N. benthamiana:Cf-4* and the two independent *rlck-vii-8* knock-out lines were subsequently treated with Avr4 protein, followed by measuring the production of ROS. As expected, Avr4/Cf-4-induced ROS production was again severely compromised in the two knock-out lines, when compared to the *N. benthamiana:Cf-4* (Figure 8C), which provides solid evidence that members of RLCK-VII-8 are indeed vital for Avr4/Cf-4-triggered ROS production. More interestingly, even though the first and second ROS peaks were reduced in size in the knock-out plants, they were still distinguishable, and additionally, the first ROS peak displayed an obvious delay. These results are different from what we observed with the *rlck-vii-6* and *rlck-vii-7* knock-out plants (Figure 6C and 7C), which again reveals that different RLCK-VII subfamilies are employed in different ways to regulate the Avr4/Cf- 4-triggered biphasic ROS burst in *N. benthamiana*. Besides, flg22-, chitin-, nlp20/RLP23-, and pg13/RLP42-induced ROS accumulation was also strongly inhibited in these knock-out plants when compared to *N. benthamiana:Cf-4* (Figure 8D, 8E, 8F and 8G), especially concerning the chitin-triggered ROS production, which was almost abolished in knock-out plants.

To further investigate the role of RLCK-VII-8 in MAPK signaling, a MAPK activation assay was performed with *N. benthamiana:Cf-4* and the two *rlck-vii-8* knock-out lines, as described above. Despite their importance in ROS signaling, members of RLCK-VII-8 seem to not be required for Avr4/Cf-4-triggered MAPK activation, as the knock-out plants displayed a MAPK activation similar to *N. benthamiana:Cf-4*, at 15 min after infiltration of the Avr4 protein (Figure 8H). Furthermore, in agreement with the HR activation data obtained with the knock-out plants of *rlck-vii-8* in the T1 generation, the Avr4/Cf-4-induced HR remained unaltered in the two independent homozygous knock-out lines, when compared to *N. benthamiana:Cf-4*, after infiltration with either Avr4 protein or with Agrobacterium harboring an *Avr4* overexpression construct (Figure 8I and 8J). Collectively, these results suggest that even though members of RLCK-VII-8 play no role, or only a minor role, in regulating the Avr4/Cf-4-triggered MAPK activation and HR, they are indeed essential for ROS production triggered by a broad spectrum of ExIPs, including Avr4.

Notably, mutations introduced in *Niben101Scf06482g03003* in both *N. benthamiana:Cf-4 rlck-vii-8* knock-out lines might actually not be inactivating this RLCK. Therefore, we could not draw a conclusion about whether this particular RLCK plays a role in regulating plant immunity.

## DISCUSSION

SOBIR1 and BAK1 are well-known co-receptors for LRR-RLPs, such as Cf-4, RLP23, and RLP42 (Albert et al., 2015; Liebrand et al., 2013; Liebrand et al., 2014; Postma et al., 2016; Zhang et al., 2021). It has been reported that an RLP/SOBIR1 complex forms heterodimers with BAK1 upon its elicitation and that subsequent trans-phosphorylation events between the kinase domains of SOBIR1 and BAK1 are required for initiating downstream immune signaling (van der Burgh et al., 2019). Both SOBIR1 and BAK1 are RD kinases, suggesting the possibility of auto-phosphorylation in their activation segment. Here, our *in vitro* studies demonstrate that, similar to their Arabidopsis homologs, *Nb*SOBIR1, *Sl*SOBIR1, *Sl*SOBIR1-like, *Nb*BAK1, and *Sl*BAK1 exhibit auto-phosphorylation activity (Figure 2A and S4) (Benjamin Schwessinger, 2011; Jia Li, 2002; Leslie et al., 2010). Furthermore, SOBIR1 indeed can directly phosphorylate BAK1, and the intrinsic kinase activity of SOBIR1 is required for this trans-phosphorylation event (Figure 2B, S4C and S4D). Accordingly, BAK1 is also able to directly phosphorylate SOBIR1, which again depends on the kinase activity of BAK1 (Figure 2C, S4E and S4F). These trans-phosphorylation events are proposed to eventually result in the full activation of SOBIR1/BAK1-containing immune complexes. As many LRR-RLPs that are involved in plant immunity require SOBIR1 and BAK1 for their function, the model that we proposed earlier (van der Burgh et al., 2019), and is now further supported by this study and by the work of Wei and colleagues (2022), probably also applies to immune signaling triggered by additional RLP/SOBIR1 complexes. Moreover, increasing evidence has indicated that BAK1 promotes the activation of the receptor complex upon the perception of an ExIP by the LRRs of the matching primary receptor. However, the signaling specificity is determined by the kinase domain of the primary receptor, which is either an RLK or the constitutive RLP/SOBIR1 complex, whereas BAK1 merely acts as a general complex activator (Hohmann et al., 2020). Therefore, after being trans-phosphorylated by BAK1, SOBIR1 is proposed to initiate the trans-phosphorylation events with downstream cytoplasmic signaling components. Such components are for example particular RLCKs, thereby triggering a specific type of immune signaling, irrespective of the RLP that is involved in the RLP/SOBIR1 complex.

Furthermore, in this study, we show that *Nb*SOBIR1 Thr522 and its analogous residues in tomato SOBIR1s (*Sl*SOBIR1 Thr513 and *Sl*SOBIR1-like Thr526), present in the activation segments of SOBIR1, are essential for their intrinsic kinase activity, and thus the Avr4/Cf- 4-mediated HR, the ROS burst and downstream MAPK activation (Figure 1, 2B, S2, S4A and S4B). These results are further supported by a recent study, which shows that *At*SOBIR1 T529A does not exhibit intrinsic kinase activity, thereby resulting in a complete loss of cell death in *N. benthamiana* upon its overexpression (Wei et al., 2022). Interestingly, the equivalent Thr residue has been proven essential for many other RD kinases. A good example is Arabidopsis RLK CHITIN ELICITOR RECEPTOR KINASE 1 (CERK1) Thr479 (Figure S1), which is indispensable for the activation of the *At*CERK1 kinase (Suzuki et al., 2016; Suzuki et al., 2019). Furthermore, the Arabidopsis LRR-RLK NUCLEAR SHUTTLE PROTEIN (NSP)-INTERACTING KINASE1 (NIK1) is a virulence target of the begomovirus NSP and is involved in plant antiviral immunity (Mariano et al., 2004; Santos et al., 2010). A mutation at Thr474, which locates at the activation segment of the kinase domain of NIK1 (Figure S1), attenuates the auto-phosphorylation of NIK1 and enhances susceptibility to the Cabbage leaf curl virus (Santos et al., 2009). BRASSINOSTEROID-INSENSITIVE 1 (BRI1) is one of the best-characterized LRR-RLKs, with essential roles in plant growth and development (Belkhadir et al., 2014; Chinchilla et al., 2009). Biochemical and genetic analyses have revealed that T1049, present in the activation segment of the kinase domain of BRI1 (Figure S1), is vital for the intrinsic kinase function of Arabidopsis BRI1, both *in vitro* and *in planta* (Wang et al., 2008). Therefore, this particular Thr residue might play a general role in regulating the intrinsic kinase activity of RD kinases.

In addition to Thr522, Tyr469 (as well as its analogous residues in *Sl*SOBIR1 and *Sl*SOBIR1- like) in the kinase domain of *Nb*SOBIR1 has also been identified to be crucial for the Avr4/Cf-4-induced HR and MAPK activation, but not for its intrinsic kinase activity and Avr4/Cf-4-triggered ROS accumulation (Figure 3, S6 and S8). Strikingly, a vital role in regulating plant immunity has recently been assigned to this particular Tyr residue present in the kinase domain of several well-known RLKs. For instance, upon the perception of elf18, the Arabidopsis LRR-RLK EFR phosphorylates at Tyr836, which is equivalent to *Nb*SOBIR1 Tyr469, and this phosphorylation is required for the activation of EFR itself and the initiation of sequential downstream immune responses. The Arabidopsis CERK1 is the co-receptor of the fungal cell wall component chitin (Miya et al., 2007; Wan et al., 2008). Phosphorylation of CERK1 Tyr428, which is also analogous to *Nb*SOBIR1 Tyr469, is essential for chitin-triggered CERK1 activation, ROS production, MAPK activation, downstream RLCK phosphorylation, and resistance to the fungal pathogen *B. cinerea* (Liu et al., 2018). Recently, the Arabidopsis lectin RLK LIPO-OLIGOSACCHARIDE-SPECIFIC REDUCED ELICITATION (LORE) has been reported to phosphorylate at Y600, which is also the analogous residue of *Nb*SOBIR1 Tyr469, upon its activation. A mutation at LORE Tyr600 results in a highly compromised LORE-mediated ROS accumulation and reduced resistance to *Pseudomonas syringae* pv. *tomato* DC3000 (Luo et al., 2020). These results collectively demonstrate the importance of this particular Tyr residue in the kinase domain of cell-surface RLKs that are involved in plant immunity. Although *At*SOBIR1 has been reported to auto-phosphorylate at Tyr residues (Leslie et al., 2010), no phosphorylated Tyr residues have been detected by MS in the kinase domain of *At*SOBIR1, when auto-phosphorylated, or after being trans-phosphorylated by BAK1 *in vitro* (Wei et al., 2022), or when produced *in planta* (van der Burgh et al., 2018). This is in contrast to the aforementioned RLKs, which can phosphorylate at this Tyr residue. Nonetheless, based on the structure of the *At*SOBIR1 kinase domain, which has been determined recently (Wei et al., 2022), *At*SOBIR1 Tyr476 (equivalent Tyr residue of *Nb*SOBIR1 Tyr469) is solvent-exposed, therefore, it might be easier to access by downstream components or other regulatory proteins (Figure S13). Therefore, this important Tyr residue in the kinase domain of SOBIR1 might not regulate plant immune responses by phosphorylation, but by interacting with specific downstream signaling partners, such as members of RLCK-VII-7 that are required for Avr4/Cf-4-induced HR (Figure 7I and 7J).

We observed that the kinetics of the Avr4-induced biphasic ROS burst displayed differently in *N. benthamiana:Cf-4 rlck-vii-6*, *rlck-vii-7* and *rlck-vii-8* (Figure 6C, 7C and 8C). For the *rlck-vii-6* knock-out lines, the second burst is specifically and completely inhibited (Figure 6C); for the *rlck-vii-7* knock-out lines, the overall ROS production is strongly attenuated, and only a weak and sustained ROS burst is triggered (Figure 7C); whereas, for the *rlck-vii-8* knock-out lines, both ROS bursts are strongly compromised, with the first burst being delayed (Figure 8C). This raises the possibility that there are different downstream ROS regulatory mechanisms in *N. benthamiana*, which together determine the ROS profile. In Arabidopsis, RESPIRATORY BURST OXIDASE HOMOLOG D (RBOHD) is engaged in extracellular ROS production, and growing evidence has suggested that RLCKs differentially regulate RBOHD activation through the differential phosphorylation of various sites in the RBOHD enzyme (Kadota et al., 2019). For instance, BIK1 directly phosphorylates the N-terminus of RBOHD to positively regulate ROS production in Arabidopsis, whereas PBL13, which negatively regulates RBOHD activation, directly phosphorylates the C-terminus of RBOHD. In *N. benthamiana*, the RBOHB homolog is responsible for the fast apoplastic ROS production during the establishment of immunity, and therefore we speculate that members from RLCK-VII-6, RLCK-VII-7 and RLCK-VII-8 phosphorylate RBOHB at different sites, thereby causing different ROS kinetics (Yoshioka et al., 2003). Furthermore, it has been reported that in *N. benthamiana* the first burst of the biphasic apoplastic ROS burst is mediated by swift RBOHB phosphorylation, whereas the second burst is the result of transcriptional upregulation of the *RBOHB* gene, a process that is mediated by activated WRKY transcription factors (Adachi et al., 2015). Hence, we hypothesize that members from RLCK-VII-6, upon their activation by the upstream cell-surface complex, phosphorylate RBOHB at specific sites for the swift ROS burst, and meanwhile directly or indirectly phosphorylating certain transcription factors to regulate the later phase of the ROS burst.

## MATERIALS AND METHODS

### Plant materials

*N. benthamiana:Cf-4 sobir1*/*sobir1-like* knock-out plants were used in this study. As *SOBIR1-like* is not functional in *N. benthamiana*, we further refer to these knock-out plants as *N. benthamiana:Cf-4 sobir1* (Huang et al., 2021).

A highly efficient multiplex editing technique employed to knock out multiple RLCK-VII subfamily members in *N. benthamiana:Cf-4* has been described previously (Stuttmann et al., 2021). To screen for homozygous transformants, genomic DNA from each mutant plant was isolated by using the Phire Tissue Direct PCR Master Mix (Thermo Fisher Scientific), followed by amplifying the sgRNA-targeted regions and subsequent Sanger sequencing of the obtained PCR fragments. Primers used for genotyping can be found in Table S2.

### Plant growth conditions

All *N. benthamiana* plants used in this study were cultivated in a climate chamber under 15 h of light at 21 °C and 9 h of darkness at 19 °C, with a relative humidity of ∼70%.

### Binary vectors for *Agrobacterium tumefaciens*-mediated transient transformation

The constructs pBIN-KS-35S::*Nb*SOBIR1-eGFP (SOL2911), pBIN-KS- 35S::*Nb*SOBIR1^D482N^-eGFP (kinase-dead mutant) (SOL7928), pBIN-KS-35S::*Sl*SOBIR1- eGFP (SOL2774), pBIN-KS-35S::*Sl*SOBIR1^D473N^-eGFP (kinase-dead mutant) (SOL2875), pBIN-KS-35S::*Sl*SOBIR1-like-eGFP (SOL2773), pBIN-KS-35S::*Sl*SOBIR1-like ^D486N^-eGFP (kinase-dead mutant) (SOL2876) and pMOG800-Avr4 have been described previously (Liebrand et al., 2013). The codon change, resulting in a Ser/Thr-to-Ala or Tyr-to-Phe amino acid change in the kinase domain of SOBIR1s, was introduced by performing overlap extension PCR using the plasmids pENTR/D-Topo:*Nb*SOBIR1 (SOL4064), pENTR/D-Topo:*Sl*SOBIR1 (SOL2746), and pENTR/D-Topo:*Sl*SOBIR1-like (SOL2745) as templates (Liebrand et al., 2013; Liu and Naismith, 2008). Phusion Hot Start II DNA Polymerase (Thermo Scientific) was used for the overlap extension PCR and the primers that were used are listed in Table S2. The methylated template plasmids remaining in the PCR products were digested by DpnI (NEB), and after transformation to *E. coli* DH5α, the required SOBIR1 mutants carrying individual mutations were selected by Sanger sequencing, and then introduced into pBIN-KS-35S::GWY-eGFP (SOL2095; for C-terminally tagging with eGFP), by using Gateway LR Clonase II (Invitrogen). Thereby, the binary vectors pBIN-KS-35S::*Nb*SOBIR1^T512A^-eGFP (SOL7909), pBIN-KS-35S::*Nb*SOBIR1^T515A^-eGFP (SOL7910), pBIN-KS-35S::*Nb*SOBIR1^T516A^-eGFP (SOL7911), pBIN-KS-35S::*Nb*SOBIR1^S517A^-eGFP (SOL7912), pBIN-KS-35S::*Nb*SOBIR1^T522A^-eGFP (SOL7913), pBIN-KS- 35S::*Sl*SOBIR1^T503A^-eGFP (SOL7969), pBIN-KS-35S::*Sl*SOBIR1^T506A^-eGFP (SOL7970), pBIN-KS-35S::*Sl*SOBIR1^T507A^-eGFP (SOL7971), pBIN-KS-35S::*Sl*SOBIR1^S508A^-eGFP (SOL7972), pBIN-KS-35S::*Sl*SOBIR1^T513A^-eGFP (SOL7973), pBIN-KS-35S::*Sl*SOBIR1- like^T516A^-eGFP (SOL7950), pBIN-KS-35S::*Sl*SOBIR1-like^T519A^-eGFP (SOL7951), pBIN-KS- 35S::*Sl*SOBIR1-like^T520A^-eGFP (SOL7952), pBIN-KS-35S::*Sl*SOBIR1-like^S521A^-eGFP (SOL7953), pBIN-KS-35S::*Sl*SOBIR1-like^T526A^-eGFP (SOL7954), pBIN-KS- 35S::*Nb*SOBIR1^Y355F^-eGFP (SOL7914), pBIN-KS-35S::*Nb*SOBIR1^Y426F^-eGFP (SOL7915), pBIN-KS-35S::*Nb*SOBIR1^Y429F^-eGFP (SOL7916), pBIN-KS-35S::*Nb*SOBIR1^Y431F^-eGFP (SOL7917), pBIN-KS-35S::*Nb*SOBIR1^Y469F^-eGFP (SOL7918), pBIN-KS- 35S::*Nb*SOBIR1^Y525F^-eGFP (SOL7919), pBIN-KS-35S::*Nb*SOBIR1^Y530F^-eGFP (SOL7920), pBIN-KS-35S::*Nb*SOBIR1^Y543F^-eGFP (SOL7921), pBIN-KS-35S::*Sl*SOBIR1^Y346F^-eGFP (SOL7974), pBIN-KS-35S::*Sl*SOBIR1^Y417F^-eGFP (SOL7975), pBIN-KS-35S::*Sl*SOBIR1^Y420F^- eGFP (SOL7976), pBIN-KS-35S::*Sl*SOBIR1^Y422F^-eGFP (SOL7977), pBIN-KS- 35S::*Sl*SOBIR1^Y460F^-eGFP (SOL7978), pBIN-KS-35S::*Sl*SOBIR1^Y521F^-eGFP (SOL7979), pBIN-KS-35S::*Sl*SOBIR1^Y522F^-eGFP (SOL7980), pBIN-KS-35S::*Sl*SOBIR1^Y534F^-eGFP (SOL7981), pBIN-KS-35S::*Sl*SOBIR1^Y588F^-eGFP (SOL7982), pBIN-KS-35S::*Sl*SOBIR1- like^Y359F^-eGFP (SOL7942), pBIN-KS-35S::*Sl*SOBIR1-like^Y430F^-eGFP (SOL7943), pBIN-KS- 35S::*Sl*SOBIR1-like^Y433F^-eGFP (SOL7944), pBIN-KS-35S::*Sl*SOBIR1-like^Y435F^-eGFP (SOL7945), pBIN-KS-35S::*Sl*SOBIR1-like^Y473F^-eGFP (SOL7946), pBIN-KS-35S::*Sl*SOBIR1- like^Y529F^-eGFP (SOL7947), pBIN-KS-35S::*Sl*SOBIR1-like^Y534F^-eGFP (SOL7948) and pBIN-KS-35S::*Sl*SOBIR1-like^Y547F^-eGFP (SOL7949), for *in planta* expression were obtained.

### *A. tumefaciens*-mediated transient transformation

All binary plasmids were transformed into *A. tumefaciens* (further referred to as Agrobacterium) strain C58C1, carrying the helper plasmid pCH32. Agrobacterium strains harboring the transient expression constructs of RLP23 and RLP42 were received from Lisha Zhang and Thorsten Nürnberger (Albert et al., 2015; Zhang et al., 2021). Infiltration of Agrobacterium into leaves of *N. benthamiana* (agro-infiltration) was performed as described before, at an OD_600_ of 0.8 (Hoorn et al., 2000).

The development of cell death was photographed and quantified by red light imaging using the ChemiDoc (Bio-Rad) (Landeo Villanueva et al., 2021). Statistical analysis was performed using one-way ANOVA by GraphPad Prism 9.

For checking the protein accumulation level of SOBIR1 mutants after their transient expression in leaves of *N. benthamiana*, a protein immunoprecipitation assay, followed by immunoblotting, was performed as described before (Thomas W.H. Liebrand, 2012).

### Reactive oxygen species (ROS) assay

For ROS burst assays, leaf discs were taken from 4-week-old *N. benthamiana:Cf-4* and *rlck* knock-out plants, while for plants transiently expressing SOBIR1 mutants for the complementation studies, *RLP23* or *RLP42*, leaf discs were collected at 24 h after ago-infiltration. Leaf discs were then floated on 80 µL of sterile water in a 96-wells plate overnight and hereafter, the water in each well was replaced carefully by 50 µL of fresh sterile water. After another 1 h of incubation, 50 µL of the reaction solution, containing 100 µM of luminol (L-012, Fujifilm, Japan), 20 µg/mL horseradish peroxidase (Sigma), and the elicitor to be tested (being 0.2 µM Avr4 protein, 0.2 μM flg22, 20 μM chitohexaose, 2 μM nlp20 or 2 μM pg13), was added to each well. Subsequently, the production of luminescence was monitored with a CLARIOstar plate reader (BMG Labtech). The line charts showing the detected values of ROS were created using GraphPad Prism 9.

### MAPK activation assay

Each SOBIR1 mutant was transiently co-expressed with *Avr4* (OD_600_=0.8 for each binary vector) in the first fully expanded leaves of *N. benthamiana:Cf-4 sobir1* plants. Leaf samples were harvested at 2 dpi, and leaf samples from the various *N. benthamiana:Cf-4 rlck* knock-out lines, infiltrated with a solution of 5 µM Avr4 protein, were harvested at 15 min after treatment.

Total protein was extracted and subjected to SDS-PAGE, after which an anti-p42/p44-erk antibody (NEB) was employed to detect the activated MAPKs on western blots.

### Recombinant protein expression

To produce the recombinant cytoplasmic kinase domain (KD) of SOBIR1 and BAK1 with a GST or 6×His tag in *E. coli*, the vectors pET-GST (Addgene No. 42049) and pET-15b were employed, respectively. Both vectors were linearized by PCR amplification with the primer pairs pET-GST_fw/rev and pET-15b_fw/rev (Table S2). Meanwhile, the coding sequence of *Nb*SOBIR1 KD, *Sl*SOBIR1 KD, and *Sl*SOBIR1-like KD, as well as all their corresponding mutants, were PCR-amplified from the corresponding pENTR/D-Topo plasmids, using the primers containing the homologous sequence of either pET-GST or pET-15b (Table S2). Hereafter, the linearized vector and amplified insert were recombined by using the ClonExpress II One Step Cloning kit (Vazyme, China). After transformation to *E. coli* DH5α, the correct expression constructs were selected by performing colony PCRs and Sanger sequencing.

For recombinant protein expression, the *E. coli* strain BL21 was used. Bacteria harboring the correct expression construct were cultured at 37 °C overnight in LB liquid medium and subsequently inoculated into fresh LB medium at a ratio of 1:200 (v/v). After culturing at 37 °C for around 3 h, the bacterial population became in an exponential growth phase, with an OD_600_ between 0.6 and 0.8, at which IPTG was added to a final concentration of 0.5 mM, followed by incubating the culture at 22 °C overnight for protein production.

### *In vitro* phosphorylation assays

*In vitro* phosphorylation assays were performed as previously described (Taylor et al., 2013). In brief, cells from 100 µL of the *E. coli* cultures that were started for protein expression were collected by centrifugation and then resuspended in 100 µL of SDS sample buffer (200 mM Tris-HCl, 8% SDS, 40% glycerol, 400 mM DTT, and 0.2% bromophenol blue), followed by boiling for 10 min. Hereafter, the samples were centrifuged for 2 min at maximum speed in an Eppendorf centrifuge, and 8 µL of the supernatant were loaded onto a precast mini-PROTEIN TGX Polyacrylamide Gel (BIORAD). After running for around 100 min at 160 V, the gel was incubated in fixation solution (50% methanol, 10% acetic acid in H_2_O), overnight. Next, the gel was washed in deionized water for 30 min twice, and the phosphorylated proteins were stained using a Pro-Q Diamond solution (Invitrogen). Subsequently, the staining solution was removed by washing the gel in de-staining solution (20% acetonitrile, 50 mM sodium acetate), and proteins were stained with Coomassie brilliant blue (CBB).

### Phylogenetic analysis of the RLCKs from Arabidopsis, *N. benthamiana,* and tomato

To conduct a phylogenetic analysis of the *At*BIK1 homologs in *N. benthamiana*, tomato, and Arabidopsis, their predicted proteomes were obtained from www.solgenomics.net and www.arabidopsis.org. Hereafter, the three predicted proteomes were independently queried for Pfam domains by using HMMER (v3.1b2; gathering cut-off) (Eddy, 1998). Sequences that contain annotated Pfam domains aside from cytoplasmic kinases (PF00069 or PF07714) were removed, and the sequences of the annotated kinase domains were extracted. Then, we took the domain PF07714 as a lead and removed the sequences that deviated in length from the kinase domain of *At*BIK1. The remaining 1,455 kinase domain sequences were aligned using MAFFT (v7.271) (Katoh and Standley, 2013), the alignment was subsequently trimmed using ClipKIT (v1.3.0; smart-gap) (Steenwyk et al., 2020) and a neighbor-joining phylogenetic tree was built using QuickTree (with 1,000 bootstrap replicates) (Howe et al., 2002) (Figure S9A). Next, a well-supported (>92% bootstrap support) sub-clade of putative BIK1 homologs, which comprised 123 sequences including *At*BIK1, was extracted from this guide tree. Subsequently, a refined phylogenetic tree was generated with these sequences by using the maximum-likelihood (ML) phylogeny as implemented in IQ-Tree (v2.2.0) (Nguyen et al., 2015). The 123 extracted kinase domains were re-aligned using MAFFT and trimmed as described above, and the ML phylogeny was constructed in IQ-Tree, using automatic amino acid substitution model selection (optimal model: Q.plant with five categories of rate heterogeneity) (Kalyaanamoorthy et al., 2017; Minh et al., 2021). Branch support for the phylogenetic tree was obtained using ultrafast bootstrap, as well as SH-aLRT, as implemented in IQ-tree (Hoang et al., 2018).

## Supporting information

Supplemental information Huang et al.

## ACKNOWLEDGMENTS

The authors thank Bert Essenstam from Unifarm for excellent plant care. Laurens Deurhof and Gabriel Lorencini Fiorin are acknowledged for technical assistance. We thank Xiaoqian Shi-Kunne for her help with retrieving the protein sequences of the SOBIR1 homologs from different plant species. Lisha Zhang and Thorsten Nürnberger (Eberhard-Karls-University, Tübingen, Germany) are acknowledged for kindly sharing the RLP23 and RLP42 binary constructs, and nlp20 and pg13 peptides. Wen R.H. Huang is supported by the China Scholarship Council (CSC). Sergio Landeo Villanueva is supported by the Peruvian Council for Science, Technology and Technological Innovation (CONCYTEC) and its executive unit FONDECYT.

## AUTHOR CONTRIBUTIONS

### Wen R.H. Huang

Took part in conceiving the project, performed experiments and wrote the manuscript.

### Ciska Braam

Generated various constructs used in this study, performed SOBIR1 complementation studies and selected *N. benthamiana:Cf-4 rlck-vii-6* homozygous knock-out lines.

### Carola Kretschmer

Generated all the *rlck* knock-out transformants in *N. benthamiana*.

### Huan Liu

Assisted with HR screening of *N. benthamiana:Cf-4 rlck* knock-out transformants, selected *N. benthamiana:Cf-4 rlck-vii-7* homozygous knock-out lines and helped with mutant phenotyping.

### Filiz Ferik

Helped with the generation of *N. benthamiana:Cf-4 rlck* knock-out transformants.

### Aranka M. van der Burgh

Initiated the studies on SOBIR1 phosphorylation and provided input during the writing process.

### Max Pluis

Selected *N. benthamiana:Cf-4 rlck-vii-8* homozygous knock-out lines and helped with mutant phenotyping.

### Agnes Omabour Hagan

Generated *SlSOBIR1* gene mutants and assisted with the complementation studies.

### Sergio Landeo Villanueva

Assisted with selecting *RLCK* genes for our knock-out studies.

### Jinbin Wu

Assisted with selecting *RLCK* genes for our knock-out studies.

### Amber van Loosbroek

Assisted with the selection of the *N. benthamiana:Cf-4 rlck-vii-8* homozygous knock-out lines and with mutant phenotyping.

### Yulu Wang

Provided the three-dimensional structure of the kinase domain of *At*SOBIR1.

### Michael F. Seidl

Performed the bioinformatics analysis of the RLCK families and contributed to writing the manuscript.

### Johannes Stuttmann

Designed and supervised the CRISPR/Cas *RLCK* gene knock-out experiments in *N. benthamiana* and contributed to writing the manuscript.

### Matthieu H.A.J. Joosten

Conceived and guided the project and assisted in writing the manuscript.

## REFERENCES

Adachi, H., Nakano, T., Miyagawa, N., Ishihama, N., Yoshioka, M., Katou, Y., Yaeno, T., Shirasu, K., and Yoshioka, H. (2015). WRKY Transcription Factors Phosphorylated by MAPK Regulate a Plant Immune NADPH Oxidase in Nicotiana benthamiana. Plant Cell. 27(9), 2645–2663. Published online 2015/09/17 DOI: 10.1105/tpc.15.00213.

Albert, I., Bohm, H., Albert, M., Feiler, C.E., Imkampe, J., Wallmeroth, N., Brancato, C., Raaymakers, T.M., Oome, S., Zhang, H., et al. (2015). An RLP23-SOBIR1-BAK1 complex mediates NLP-triggered immunity. Nat Plants. 1, 15140. Published online 2015/01/01 DOI: 10.1038/nplants.2015.140.

Albert, I., Hua, C., Nurnberger, T., Pruitt, R.N., and Zhang, L. (2020). Surface Sensor Systems in Plant Immunity. Plant Physiol. 182(4), 1582–1596. Published online 2019/12/12 DOI: 10.1104/pp.19.01299.

Bailey, T.L., Boden, M., Buske, F.A., Frith, M., Grant, C.E., Clementi, L., Ren, J., Li, W.W., and Noble, W.S. (2009). MEME SUITE: tools for motif discovery and searching. Nucleic Acids Res. 37(Web Server issue), W202-208. Published online 2009/05/22 DOI: 10.1093/nar/gkp335.

Belkhadir, Y., Yang, L., Hetzel, J., Dangl, J.L., and Chory, J. (2014). The growth-defense pivot: crisis management in plants mediated by LRR-RK surface receptors. Trends Biochem Sci. 39(10), 447–456. Published online 2014/08/05 DOI: 10.1016/j.tibs.2014.06.006.

Benjamin Schwessinger, M.R., Yasuhiro Kadota, Vardis Ntoukakis, Jan Sklenar, Alexandra Jones, Cyril Zipfel (2011). Phosphorylation-Dependent Differential Regulation of Plant Growth, Cell Death, and Innate Immunity by the Regulatory Receptor-Like Kinase BAK1. PLoS Genetics. 7(4), e1002046. DOI: 10.1371/journal.pgen.1002046.

Bi, G., Zhou, Z., Wang, W., Li, L., Rao, S., Wu, Y., Zhang, X., Menke, F.L.H., Chen, S., and Zhou, J.M. (2018). Receptor-Like Cytoplasmic Kinases Directly Link Diverse Pattern Recognition Receptors to the Activation of Mitogen-Activated Protein Kinase Cascades in Arabidopsis. Plant Cell. 30(7), 1543–1561. Published online 2018/06/07 DOI: 10.1105/tpc.17.00981.

Bleecker, S.-H.S.a.A.B. (2001). Receptor-like kinases from Arabidopsis form a monophyletic gene family related to animal receptor kinases. PNAS. 98.

Bombarely, A., Rosli, H.G., Vrebalov, J., Moffett, P., Mueller, L.A., and Martin, G.B. (2012). A draft genome sequence of Nicotiana benthamiana to enhance molecular plant-microbe biology research. Mol Plant Microbe Interact. 25(12), 1523–1530. Published online 2012/08/11 DOI: 10.1094/MPMI-06-12-0148-TA.

Burgh, A.M.v.d., and Joosten, M.H.A.J. (2019). Plant Immunity: Thinking Outside and Inside the Box. Trends in Plant Science.

Burgh, A.M.v.d., Postma, J., Robatzek, S., and Joosten, M.H.A.J. (2019). Kinase activity of SOBIR1 and BAK1 is required for immune signalling. Molecular Plant Pathology. 20(3), 410–422. Published online 2018/11/09 DOI: 10.1111/mpp.12767.

Burgh, A.M.v.d., Thomma, B.P.H.J.P.d., Joosten, M.H.A.J.D., Burgh, A.M.v.d., Thomma, B.P.H.J.P.d., and Joosten, M.H.A.J.D., 2018. SOBIR1-containing immune complexes at the plant cell surface: partners and signalling. Wageningen University, Wageningen.

Cai, X., Takken, F.L.W., Joosten, M.H.A.J., and Wit, P.J.G.M.d. (2001). Specific recognition of AVR4 and AVR9 results in distinct patterns of hypersensitive cell death in tomato, but similar patterns of defence-related gene expression. MOLECULAR PLANT PATHOLOGY. 2, 77–86.

Chinchilla, D., Shan, L., He, P., de Vries, S., and Kemmerling, B. (2009). One for all: the receptor-associated kinase BAK1. Trends Plant Sci. 14(10), 535–541. Published online 2009/09/15 DOI: 10.1016/j.tplants.2009.08.002.

Chinchilla, D., Zipfel, C., Robatzek, S., Kemmerling, B., Nürnberger, T., Jones, J.D.G., Felix, G., and Boller, T. (2007). A flagellin-induced complex of the receptor FLS2 and BAK1 initiates plant defence. Nature. 448(7152), 497–500. Published online 2007/07/13 DOI: 10.1038/nature05999.

Chisholm, S.T., Coaker, G., Day, B., and Staskawicz, B.J. (2006). Host-Microbe Interactions: Shaping the Evolution of the Plant Immune Response. Cel. 124. Published online 814 DOI: 10.1016/j.cell.2006.02.008.

Dardick, C., Schwessinger, B., and Ronald, P. (2012). Non-arginine-aspartate (non-RD) kinases are associated with innate immune receptors that recognize conserved microbial signatures. Curr Opin Plant Biol. 15(4), 358–366. Published online 2012/06/05 DOI: 10.1016/j.pbi.2012.05.002.

de Wit, P.J. (2016). Cladosporium fulvum Effectors: Weapons in the Arms Race with Tomato. Annu Rev Phytopathol. 54, 1–23. Published online 2016/05/25 DOI: 10.1146/annurev-phyto-011516-040249.

DeFalco, T.A., and Zipfel, C. (2021). Molecular mechanisms of early plant pattern-triggered immune signaling. Mol Cell. Published online 2021/08/18 DOI: 10.1016/j.molcel.2021.07.029.

Eddy, S.R. (1998). Profile hidden Markov models. Bioinformatics. 14, 755–763.

G. Manning, D.B.W., 1 R. Martinez, 1 T. Hunter, 2 S. Sudarsanam 1,3 (2002). The Protein Kinase Complement of the Human Genome. SCIENCE. 298.

Gabriëls, S.H., Vossen, J.H., Ekengren, S.K., van Ooijen, G., Abd-El-Haliem, A.M., van den Berg, G.C., Rainey, D.Y., Martin, G.B., Takken, F.L., de Wit, P.J., et al. (2007). An NB-LRR protein required for HR signalling mediated by both extra-and intracellular resistance proteins. Plant J. 50(1), 14–28. Published online 2007/03/10 DOI: 10.1111/j.1365-313X.2007.03027.x.

Gabriëls, S.H.E.J., Takken, F.L.W., Vossen, J.H., Jong, C.F.d., Liu, Q., Turk, S.C.H.J., Wachowski, L.K., Peters, J., Witsenboer, H.M.A., Wit, P.J.G.M.d., et al. (2006). cDNA-AFLP combined with functional analysis reveals novel genes involved in the hypersensitive response. Mol Plant Microbe Interact. 19.

Gao, M., Wang, X., Wang, D., Xu, F., Ding, X., Zhang, Z., Bi, D., Cheng, Y.T., Chen, S., Li, X., et al. (2009). Regulation of cell death and innate immunity by two receptor-like kinases in Arabidopsis. Cell Host & Microbe. 6(1), 34–44.

Gómez-Gómez, L., Bauer, Z., and Boller, T. (2001). Both the Extracellular Leucine-Rich Repeat Domain and the Kinase Activity of FLS2 Are Required for Flagellin Binding and Signaling in Arabidopsis. The Plant Cell. 13.

Gómez-Gómez, L., and Boller, T. (2000). FLS2: An LRR Receptor–like Kinase Involved in the Perception of the Bacterial Elicitor Flagellin in Arabidopsis. Molecular Cell. 5.

Goodin, M.M., Zaitlin, D., Naidu, R.A., and Lomme, S.A. (2008). Nicotiana benthamiana: Its History and Future as a Model for Plant–Pathogen Interactions. MPMI. 21.

Heese, A., Hann, D.R., Gimenez-Ibanez, S., Jones, A.M.E., He, K., Li, J., Schroeder, J.I., Peck, S.C., and Rathjen, J.P. (2007). The receptor-like kinase SERK3/BAK1 is a central regulator of innate immunity in plants. Proceedings of the National Academy of Sciences of the United States of America. 104(29).

Hoang, D.T., Chernomor, O., von Haeseler, A., Minh, B.Q., and Vinh, L.S. (2018). UFBoot2: Improving the Ultrafast Bootstrap Approximation. Mol Biol Evol. 35(2), 518–522. Published online 2017/10/28 DOI: 10.1093/molbev/msx281.

Hohmann, U., Ramakrishna, P., Wang, K., Lorenzo-Orts, L., Nicolet, J., Henschen, A., Barberon, M., Bayer, M., and Hothorn, M. (2020). Constitutive Activation of Leucine-Rich Repeat Receptor Kinase Signaling Pathways by BAK1-Interacting Receptor-Like Kinase 3 Chimera. Plant Cell. Published online 2020/08/17 DOI: 10.1105/tpc.20.00138.

Hoorn, R.A.L.V.d., Laurent, F., Roth, R., and Wit, P.J.G.M.D. (2000). Agroinfiltration Is a Versatile Tool That Facilitates Comparative Analyses of Avr9/Cf-9-Induced and Avr4/Cf-4-Induced Necrosis. Molecular Plant-Microbe Interactions. 13.

Howe, K., Bateman, A., and Durbin, R. (2002). QuickTree: building huge Neighbour-Joining trees of protein sequences. Bioinformatics. 18, 1546–1547.

Huang, W.R.H., Schol, C., Villanueva, S.L., Heidstra, R., and Joosten, M.H.A.J. (2021). Knocking out SOBIR1 in Nicotiana benthamiana abolishes functionality of transgenic receptor-like protein Cf-4. Plant Physiology. DOI: 10.1093/plphys/kiaa047.

Jia Li, J.W., Kevin A. Lease, Jason T. Doke, Frans E. Tax, and John C. Walker (2002). BAK1, an Arabidopsis LRR Receptor-like Protein Kinase, Interacts with BRI1 and Modulates Brassinosteroid Signaling. Cell. 110.

Johnson, L.N., Noble, M.E.M., and Owen, D.J. (1996). Active and Inactive Protein Kinases: Structural Basis for Regulation. Cell. 85, 10. Published online 158.

Joosten, M.H.A.J., J.Cozijnsen, T., and Wil, P.J.G.M.d. (1994). Host resistance to a fungal tomato pathogen lost by a single base-pair change in an avirulence gene. NATURE. 367(27).

Kadota, Y., Liebrand, T.W.H., Goto, Y., Sklenar, J., Derbyshire, P., Menke, F.L.H., Torres, M.A., Molina, A., Zipfel, C., Coaker, G., et al. (2019). Quantitative phosphoproteomic analysis reveals common regulatory mechanisms between effector-and PAMP-triggered immunity in plants. New Phytol. 221(4), 2160–2175. Published online 2018/10/10 DOI: 10.1111/nph.15523.

Kalyaanamoorthy, S., Minh, B.Q., Wong, T.K.F., von Haeseler, A., and Jermiin, L.S. (2017). ModelFinder: fast model selection for accurate phylogenetic estimates. Nat Methods. 14(6), 587–589. Published online 2017/05/10 o.

Katoh, K., and Standley, D.M. (2013). MAFFT multiple sequence alignment software version 7: improvements in performance and usability. Mol Biol Evol. 30(4), 772–780. Published online 2013/01/19 DOI: 10.1093/molbev/mst010.

Kong, L., Rodrigues, B., Kim, J.H., He, P., and Shan, L. (2021). More than an on-and-off switch: Post-translational modifications of plant pattern recognition receptor complexes. Curr Opin Plant Biol. 63, 102051. Published online 2021/05/23 DOI: 10.1016/j.pbi.2021.102051.

Kourelis, J., Kaschani, F., Grosse-Holz, F.M., Homma, F., Kaiser, M., and van der Hoorn, R.A.L. (2019). A homology-guided, genome-based proteome for improved proteomics in the alloploid Nicotiana benthamiana. BMC Genomics. 20(1), 722. Published online 2019/10/06 DOI: 10.1186/s12864-019-6058-6.

Landeo Villanueva, S., Malvestiti, M.C., van Ieperen, W., Joosten, M., and van Kan, J.A.L. (2021). Red light imaging for programmed cell death visualization and quantification in plant-pathogen interactions. Mol Plant Pathol. Published online 2021/01/27 DOI: 10.1111/mpp.13027.

Lee, D.H., Lee, H.S., and Belkhadir, Y. (2021). Coding of plant immune signals by surface receptors. Curr Opin Plant Biol. 62, 102044. Published online 2021/05/13 DOI: 10.1016/j.pbi.2021.102044.

Leslie, M.E., Lewis, M.W., Youn, J.-Y., Daniels, M.J., and Liljegren, S.J. (2010). The EVERSHED receptor-like kinase modulates floral organ shedding in Arabidopsis. Development. 137. Published online 476 DOI: 10.1242/dev.041335.

Li, P., Zhao, L., Qi, F., Htwe, N., Li, Q., Zhang, D., Lin, F., Shang-Guan, K., and Liang, Y. (2021). The receptor-like cytoplasmic kinase RIPK regulates broad-spectrum ROS signaling in multiple layers of plant immune system. Mol Plant. Published online 2021/06/16 DOI: 10.1016/j.molp.2021.06.010.

Liang, X., and Zhou, J.M. (2018). Receptor-Like Cytoplasmic Kinases: Central Players in Plant Receptor Kinase-Mediated Signaling. Annu Rev Plant Biol. 69, 267–299. Published online 2018/05/03 DOI: 10.1146/annurev-arplant-042817-040540.

Liebrand, T.W.H., Berg, G.C.M.v.d., Zhang, Z., Smit, P., Cordewener, J.H.G., America, A.H.P., Sklenar, J., Jones, A.M.E., Tameling, W.I.L., Robatzek, S., et al. (2013). Receptor-like kinase SOBIR1/EVR interacts with receptor-like proteins in plant immunity against fungal infection. Proceedings of the National Academy of Sciences of the United States of America. 110(24). Published online 10015 DOI: 10.1073/pnas.1313401110.

Liebrand, T.W.H., Burg, H.A.v.d., and Joosten, M.H.A.J. (2014). Two for all: receptor-associated kinases SOBIR1 and BAK1. Trends in Plant Science. 19(2), 123–132.

Lin, W., BoLi, Lu, D., Chen, S., Zhu, N., He, P., and Shan, L. (2014). Tyrosine phosphorylation of protein kinase complex BAK1/BIK1 mediates Arabidopsis innate immunity. pnas. 111(9). DOI: 10.1073/pnas.1318817111.

Lin, W., Ma, X., Shan, L., and He, P. (2013). Big roles of small kinases: the complex functions of receptor-like cytoplasmic kinases in plant immunity and development. J Integr Plant Biol. 55(12), 1188–1197. Published online 2013/05/29 DOI: 10.1111/jipb.12071.

Lin, Z.J., Liebrand, T.W., Yadeta, K.A., and Coaker, G. (2015). PBL13 Is a Serine/Threonine Protein Kinase That Negatively Regulates Arabidopsis Immune Responses. Plant Physiol. 169(4), 2950–2962. Published online 2015/10/04 DOI: 10.1104/pp.15.01391.

Liu, H., and Naismith, J.H. (2008). An efficient one-step site-directed deletion, insertion, single and multiple-site plasmid mutagenesis protocol. BMC Biotechnol. 8, 91. Published online 2008/12/06 DOI: 10.1186/1472-6750-8-91.

Liu, J., Liu, B., Chen, S., Gong, B.-Q., Chen, L., Zhou, Q., Xiong, F., Wang, M., Feng, D., Li, J.-F., et al. (2018). A Tyrosine Phosphorylation Cycle Regulates Fungal Activation of a Plant Receptor Ser/Thr Kinase. Cell Host & Microbe 23, 1. DOI: 10.1016/j.chom.2017.12.005.

Lu, D., Wu, S., Gao, X., Zhang, Y., Shan, L., and He, P. (2010). A receptor-like cytoplasmic kinase, BIK1, associates with a flagellin receptor complex to initiate plant innate immunity. Proceedings of the National Academy of Sciences of the United States of America. 107(1), 496-501. Published online 2009/12/19 DOI: 10.1073/pnas.0909705107.

Lu, Y., and Tsuda, K. (2021). Intimate Association of PRR-and NLR-Mediated Signaling in Plant Immunity. Mol Plant Microbe Interact. 34(1), 3–14. Published online 2020/10/14 DOI: 10.1094/MPMI-08-20-0239-IA.

Luo, X., Wu, W., Liang, Y., Xu, N., Wang, Z., Zou, H., and Liu, J. (2020). Tyrosine phosphorylation of the lectin receptor-like kinase LORE regulates plant immunity. EMBO J. e102856. Published online 2020/01/11 DOI: 10.15252/embj.2019102856.

Macho, A.P., and Zipfel, C. (2014). Plant PRRs and the Activation of Innate Immune Signaling. Molecular Cell. 54. DOI: 10.1016/j.molcel.2014.03.028.

Mariano, A.C., Andrade, M.O., Santos, A.A., Carolino, S.M., Oliveira, M.L., Baracat-Pereira, M.C., Brommonshenkel, S.H., and Fontes, E.P. (2004). Identification of a novel receptor-like protein kinase that interacts with a geminivirus nuclear shuttle protein. Virology. 318(1), 24–31. Published online 2004/02/20 DOI: 10.1016/j.virol.2003.09.038.

Minh, B.Q., Dang, C.C., Vinh, L.S., and Lanfear, R. (2021). QMaker: Fast and Accurate Method to Estimate Empirical Models of Protein Evolution. Syst Biol. 70(5), 1046–1060. Published online 2021/02/23 DOI: 10.1093/sysbio/syab010.

Mithoe, S.C., and Menke, F.L.H. (2018). Regulation of pattern recognition receptor signalling by phosphorylation and ubiquitination. Current Opinion in Plant Biology. 45, 162–170. DOI: 10.1016/j.pbi.2018.07.008.

Miya, A., Albert, P., Shinya, T., Desaki, Y., Ichimura, K., Shirasu, K., Narusaka, Y., Kawakami, N., Kaku, H., and Shibuya, N. (2007). CERK1, a LysM receptor kinase, is essential for chitin elicitor signaling in Arabidopsis. PNAS. 104(49). Published online 19618.

Monaghan, J., and Zipfel, C. (2012). Plant pattern recognition receptor complexes at the plasma membrane. Current Opinion in Plant Biology. 15. DOI: 10.1016/j.pbi.2012.05.006.

Nguyen, L.T., Schmidt, H.A., von Haeseler, A., and Minh, B.Q. (2015). IQ-TREE: a fast and effective stochastic algorithm for estimating maximum-likelihood phylogenies. Mol Biol Evol. 32(1), 268–274. Published online 2014/11/06 DOI: 10.1093/molbev/msu300.

Nolen, B., Taylor, S., and Ghosh, G. (2004). Regulation of protein kinases; controlling activity through activation segment conformation. Mol Cell. 15(5), 661–675. Published online 2004/09/08 DOI: 10.1016/j.molcel.2004.08.024.

Oliver, A.W., Knapp, S., and Pearl, L.H. (2007). Activation segment exchange: a common mechanism of kinase autophosphorylation? Trends Biochem Sci. 32(8), 351–356. Published online 2007/07/14 DOI: 10.1016/j.tibs.2007.06.004.

Perraki, A., DeFalco, T.A., Derbyshire, P., Avila, J., Sere, D., Sklenar, J., Qi, X., Stransfeld, L., Schwessinger, B., Kadota, Y., et al. (2018). Phosphocode-dependent functional dichotomy of a common co-receptor in plant signalling. Nature. 561(7722), 248–252. Published online 2018/09/05 DOI: 10.1038/s41586-018-0471-x.

Postma, J., Liebrand, T.W.H., Bi, G., Evrard, A., Bye, R.R., Mbengue, M., Kuhn, H., Joosten, M.H.A.J., and Robatzek, S. (2016). Avr4 promotes Cf-4 receptor-like protein association with the BAK1/SERK3 receptor-like kinase to initiate receptor endocytosis and plant immunity. New Phytologist. DOI: 10.1111/nph.13802.

Pruitt, R.N., Locci, F., Wanke, F., Zhang, L., Saile, S.C., Joe, A., Karelina, D., Hua, C., Frohlich, K., Wan, W.L., et al. (2021). The EDS1-PAD4-ADR1 node mediates Arabidopsis pattern-triggered immunity. Nature. Published online 2021/09/10 DOI: 10.1038/s41586-021-03829-0.

Qi, J., Wang, J., Gong, Z., and Zhou, J.M. (2017). Apoplastic ROS signaling in plant immunity. Curr Opin Plant Biol. 38, 92–100. Published online 2017/05/17 DOI: 10.1016/j.pbi.2017.04.022.

Ranf, S. (2017). Sensing of molecular patterns through cell surface immune receptors. Curr Opin Plant Biol. 38, 68–77. Published online 2017/05/14 DOI: 10.1016/j.pbi.2017.04.011.

Rao, S., Zhou, Z., Miao, P., Bi, G., Hu, M., Wu, Y., Feng, F., Zhang, X., and Zhou, J.M. (2018). Roles of Receptor-Like Cytoplasmic Kinase VII Members in Pattern-Triggered Immune Signaling. Plant Physiol. 177(4), 1679–1690. Published online 2018/06/17 DOI: 10.1104/pp.18.00486.

Santos, A.A., Carvalho, C.M., Florentino, L.H., Ramos, H.J., and Fontes, E.P. (2009). Conserved threonine residues within the A-loop of the receptor NIK differentially regulate the kinase function required for antiviral signaling. PLoS One. 4(6), e5781. Published online 2009/06/06 DOI: 10.1371/journal.pone.0005781.

Santos, A.A., Lopes, K.V., Apfata, J.A., and Fontes, E.P. (2010). NSP-interacting kinase, NIK: a transducer of plant defence signalling. J Exp Bot. 61(14), 3839–3845. Published online 2010/07/14 DOI: 10.1093/jxb/erq219.

Sharma, P.C., Ito, A., Shimizu, T., Terauchi, R., Kamoun, S., and Saitoh, H. (2003). Virus-induced silencing of WIPK and SIPK genes reduces resistance to a bacterial pathogen, but has no effect on the INF1-induced hypersensitive response (HR) in Nicotiana benthamiana. Mol Genet Genomics. 269(5), 583–591. Published online 2003/07/03 DOI: 10.1007/s00438-003-0872-9.

Shinya, T., Yamaguchi, K., Desaki, Y., Yamada, K., Narisawa, T., Kobayashi, Y., Maeda, K., Suzuki, M., Tanimoto, T., Takeda, J., et al. (2014). Selective regulation of the chitin-induced defense response by the Arabidopsis receptor-like cytoplasmic kinase PBL27. Plant J. 79(1), 56–66. Published online 2014/04/23 DOI: 10.1111/tpj.12535.

Shiu, S.-H., and Bleecker, A.B. (2001). Receptor-like kinases from Arabidopsis form a monophyletic gene family related to animal receptor kinases. PNAS. 98(19), 10763–10768.

Shiu, S.H., Karlowski, W.M., Pan, R., Tzeng, Y.H., Mayer, K.F., and Li, W.H. (2004). Comparative analysis of the receptor-like kinase family in Arabidopsis and rice. Plant Cell. 16(5), 1220–1234. Published online 2004/04/24 DOI: 10.1105/tpc.020834.

Steenwyk, J.L., Buida, T.J., 3rd, Li, Y., Shen, X.X., and Rokas, A. (2020). ClipKIT: A multiple sequence alignment trimming software for accurate phylogenomic inference. PLoS Biol. 18(12), e3001007. Published online 2020/12/03 DOI: 10.1371/journal.pbio.3001007.

Steinbrenner, A.D. (2020). The evolving landscape of cell surface pattern recognition across plant immune networks. Curr Opin Plant Biol. 56, 135–146. Published online 2020/07/03 DOI: 10.1016/j.pbi.2020.05.001.

Stuttmann, J., Barthel, K., Martin, P., Ordon, J., Erickson, J.L., Herr, R., Ferik, F., Kretschmer, C., Berner, T., Keilwagen, J., et al. (2021). Highly efficient multiplex editing: one-shot generation of 8x Nicotiana benthamiana and 12x Arabidopsis mutants. Plant J. 106(1), 8–22. Published online 2021/02/13 DOI: 10.1111/tpj.15197.

Sun, Y., Li, L., Macho, A.P., Han, Z., Hu, Z., Zipfel, C., Zhou, J.-M., and Chai, J. (2013). Structural Basis for flg22-Induced Activation of the Arabidopsis FLS2-BAK1 Immune Complex. SCIENCE. 342.

Suzuki, M., Shibuya, M., Shimada, H., Motoyama, N., Nakashima, M., Takahashi, S., Suto, K., Yoshida, I., Matsui, S., Tsujimoto, N., et al. (2016). Autophosphorylation of Specific Threonine and Tyrosine Residues in Arabidopsis CERK1 is Essential for the Activation of Chitin-Induced Immune Signaling. Plant Cell Physiol. 57(11), 2312–2322. Published online 2016/08/28 DOI: 10.1093/pcp/pcw150.

Suzuki, M., Yoshida, I., Suto, K., Desaki, Y., Shibuya, N., and Kaku, H. (2019). AtCERK1 phosphorylation site S493 contributes to the transphosphorylation of downstream components for chitin-induced immune signaling. Plant Cell Physiol. Published online 2019/05/24 DOI: 10.1093/pcp/pcz096.

Tameling, W.I., Nooijen, C., Ludwig, N., Boter, M., Slootweg, E., Goverse, A., Shirasu, K., and Joosten, M.H. (2010). RanGAP2 mediates nucleocytoplasmic partitioning of the NB-LRR immune receptor Rx in the Solanaceae, thereby dictating Rx function. Plant Cell. 22(12), 4176–4194. Published online 2010/12/21 DOI: 10.1105/tpc.110.077461.

Tang, D., Wang, G., and Zhou, J.M. (2017). Receptor Kinases in Plant-Pathogen Interactions: More Than Pattern Recognition. Plant Cell. 29(4), 618–637. Published online 2017/03/18 DOI: 10.1105/tpc.16.00891.

Taylor, I., Seitz, K., Bennewitz, S., and Walker, J.C. (2013). A simple in vitro method to measure autophosphorylation of protein kinases. Plant Methods. 9(22).

Thomas, C.M., Jones, D.A., Parniske, M., Harrison, K., Balint-Kurti, P.J., Hatzixanthis, K., and Jones, J.D.G. (1997). Characterization of the Tomato Cf-4 Gene for Resistance to Cladosporium fulvum ldentifies Sequences That Determine Recognitional Specificity in Cf-4 and Cf-9. The Plant Cell. 9.

Thomas W.H. Liebrand, P.S., Ahmed Abd-El-Haliem 2, Ronnie de Jonge, Jan H.G. Cordewener, Antoine H.P. America, Jan Sklenar, Alexandra M.E. Jones, Silke Robatzek, Bart P.H.J. Thomma, Wladimir I.L. Tameling, and Matthieu H.A.J. Joosten* (2012). Endoplasmic Reticulum-Quality Control Chaperones Facilitate the Biogenesis of Cf Receptor-Like Proteins Involved in Pathogen Resistance of Tomato1[C][W]. Plant Physiology. 159. Published online 1833 DOI: 10.1104/pp.112.196741.

Tsuda, K., and Katagiri, F. (2010). Comparing signaling mechanisms engaged in pattern-triggered and effector-triggered immunity. Curr Opin Plant Biol. 13(4), 459–465. Published online 2010/05/18 DOI: 10.1016/j.pbi.2010.04.006.

Wan, J., Zhang, X.C., Neece, D., Ramonell, K.M., Clough, S., Kim, S.Y., Stacey, M.G., and Stacey, G. (2008). A LysM receptor-like kinase plays a critical role in chitin signaling and fungal resistance in Arabidopsis. Plant Cell. 20(2), 471–481. Published online 2008/02/12 DOI: 10.1105/tpc.107.056754.

Wan, W.-L., Fröhlich, K., Pruitt, R.N., Nürnberger, T., and Zhang, L. (2019). Plant cell surface immune receptor complex signaling. Current Opinion in Plant Biology. 50, 18–28. DOI: 10.1016/j.pbi.2019.02.001.

Wan, W.L., Zhang, L., Pruitt, R., Zaidem, M., Brugman, R., Ma, X., Krol, E., Perraki, A., Kilian, J., and Grossmann, G. (2019). Comparing Arabidopsis receptor kinase and receptor protein-mediated immune signaling reveals BIK1-dependent differences. New Phytologist. 221(4), 2080–2095.

Wang, J., and Chai, J. (2020). Structural Insights into the Plant Immune Receptors PRRs and NLRs. Plant Physiol. 182(4), 1566–1581. Published online 2020/02/13 DOI: 10.1104/pp.19.01252.

Wang, X., Goshe, M.B., Soderblom, E.J., Phinney, B.S., Kuchar, J.A., Li, J., Asami, T., Yoshida, S., Huber, S.C., and Clouse, S.D. (2005). Identification and functional analysis of in vivo phosphorylation sites of the Arabidopsis BRASSINOSTEROID-INSENSITIVE1 receptor kinase. Plant Cell. 17(6), 1685–1703. Published online 2005/05/17 DOI: 10.1105/tpc.105.031393.

Wang, X., Kota, U., He, K., Blackburn, K., Li, J., Goshe, M.B., Huber, S.C., and Clouse, S.D. (2008). Sequential transphosphorylation of the BRI1Bak1 receptor kinase complex impacts early events in Brassinosteroid signaling. Developmental Cell. 15. DOI: 10.1016/j.devcel.2008.06.011.

Waszczak, C., Carmody, M., and Kangasjarvi, J. (2018). Reactive Oxygen Species in Plant Signaling. Annu Rev Plant Biol. 69, 209–236. Published online 2018/03/01 DOI: 10.1146/annurev-arplant-042817-040322.

Wei, X., Wang, Y., Zhang, S., Gu, T., Steinmetz, G., Yu, H., Guo, G., Liu, X., Fan, S., Wang, F., et al. (2022). Structural analysis of receptor-like kinase SOBIR1 revealed mechanisms that regulate its phosphorylation-dependent activation. Plant Communications. DOI: 10.1016/j.xplc.2022.100301.

Wendell A. Lim, T.P. (2010). Phosphotyrosine Signaling: Evolving a New Cellular Communication System. CELL. 142. DOI: 10.1016/j.cell.2010.08.023.

Yamada, K., Yamaguchi, K., Shirakawa, T., Nakagami, H., Mine, A., Ishikawa, K., Fujiwara, M., Narusaka, M., Narusaka, Y., Ichimura, K., et al. (2016). The Arabidopsis CERK1-associated kinase PBL27 connects chitin perception to MAPK activation. EMBO J. 35(22), 2468–2483. Published online 2016/09/30 DOI: 10.15252/embj.201694248.

Yoshioka, H., Numata, N., Nakajima, K., Katou, S., Kawakita, K., Rowland, O., Jones, J.D., and Doke, N. (2003). Nicotiana benthamiana gp91phox homologs NbrbohA and NbrbohB participate in H2O2 accumulation and resistance to Phytophthora infestans. Plant Cell. 15(3), 706–718. Published online 2003/03/05 DOI: 10.1105/tpc.008680.

Yu, X., Feng, B., He, P., and Shan, L. (2017). From Chaos to Harmony: Responses and Signaling upon Microbial Pattern Recognition. Annu Rev Phytopathol. 55, 109–137. Published online 2017/05/20 DOI: 10.1146/annurev-phyto-080516-035649.

Yuan, M., Ngou, B.P.M., Ding, P., and Xin, X.F. (2021). PTI-ETI crosstalk: an integrative view of plant immunity. Curr Opin Plant Biol. 62, 102030. Published online 2021/03/09 DOI: 10.1016/j.pbi.2021.102030.

Zhang, J., Li, W., Xiang, T., Liu, Z., Laluk, K., Ding, X., Zou, Y., Gao, M., Zhang, X., Chen, S., et al. (2010). Receptor-like cytoplasmic kinases integrate signaling from multiple plant immune receptors and are targeted by a Pseudomonas syringae effector. Cell Host Microbe. 7(4), 290–301. Published online 2010/04/24 DOI: 10.1016/j.chom.2010.03.007.

Zhang, L., Hua, C., Pruitt, R.N., Qin, S., Wang, L., Albert, I., Albert, M., van Kan, J.A.L., and Nurnberger, T. (2021). Distinct immune sensor systems for fungal endopolygalacturonases in closely related Brassicaceae. Nat Plants. Published online 2021/07/31 DOI: 10.1038/s41477-021-00982-2.

Zhang, L., Kars, I., Essenstam, B., Liebrand, T.W., Wagemakers, L., Elberse, J., Tagkalaki, P., Tjoitang, D., van den Ackerveken, G., and van Kan, J.A. (2014). Fungal endopolygalacturonases are recognized as microbe-associated molecular patterns by the arabidopsis receptor-like protein RESPONSIVENESS TO BOTRYTIS POLYGALACTURONASES1. Plant Physiol. 164(1), 352–364. Published online 2013/11/22 DOI: 10.1104/pp.113.230698.

Zhang, M., Su, J., Zhang, Y., Xu, J., and Zhang, S. (2018). Conveying endogenous and exogenous signals: MAPK cascades in plant growth and defense. Curr Opin Plant Biol. 45(Pt A), 1–10. Published online 2018/05/13 DOI: 10.1016/j.pbi.2018.04.012.

Zhou, J.-M., and Zhang, Y. (2020). Plant Immunity: Danger Perception and Signaling. Cell. 181(5), 978–989. Published online 2020/05/23 DOI: 10.1016/j.cell.2020.04.028.

Zipfel, C., Kunze, G., Chinchilla, D., Caniard, A., Jones, J.D., Boller, T., and Felix, G. (2006). Perception of the bacterial PAMP EF-Tu by the receptor EFR restricts Agrobacterium-mediated transformation. Cell. 125(4), 749–760. Published online 2006/05/23 DOI: 10.1016/j.cell.2006.03.037.

