## Supplemental information Huang et al. for "Receptor-like cytoplasmic kinases belonging to different subfamilies mediate immune responses downstream of the Cf-4 resistance protein in *Nicotiana benthamiana*"

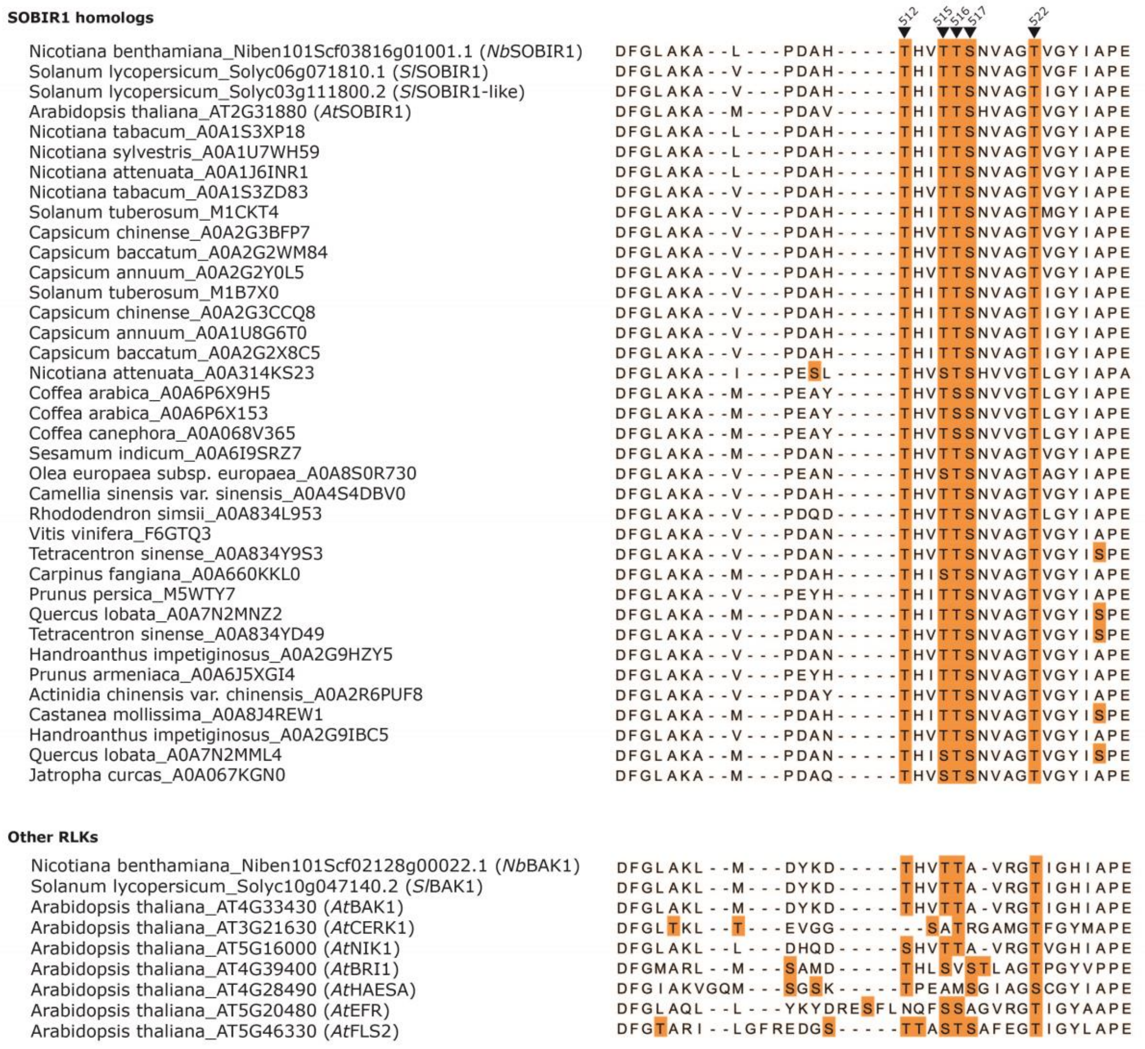

**Figure S1. Five potential phosphorylation sites in the activation segment of *Nb*SOBIR1 are highly conserved in the kinase domain of various RLKs from different plant species.** Protein sequences of the SOBIR1 homologs from various plant species were obtained from the UniProt database (https://www.uniprot.org/), while the protein sequences of other RLKs were retrieved from TAIR (https://www.arabidopsis.org/index.jsp) and the Sol Genomics Network (https://solgenomics.net/). The alignment was visualized using JalView and only the amino acid sequences of the activation segment of the kinase domain are shown. All the putatively phosphorylatable Ser (S) and Thr (T) residues are highlighted in orange. The positions of the five Ser/Thr residues of *Nb*SOBIR1 that are subjected to this study are indicated on top.

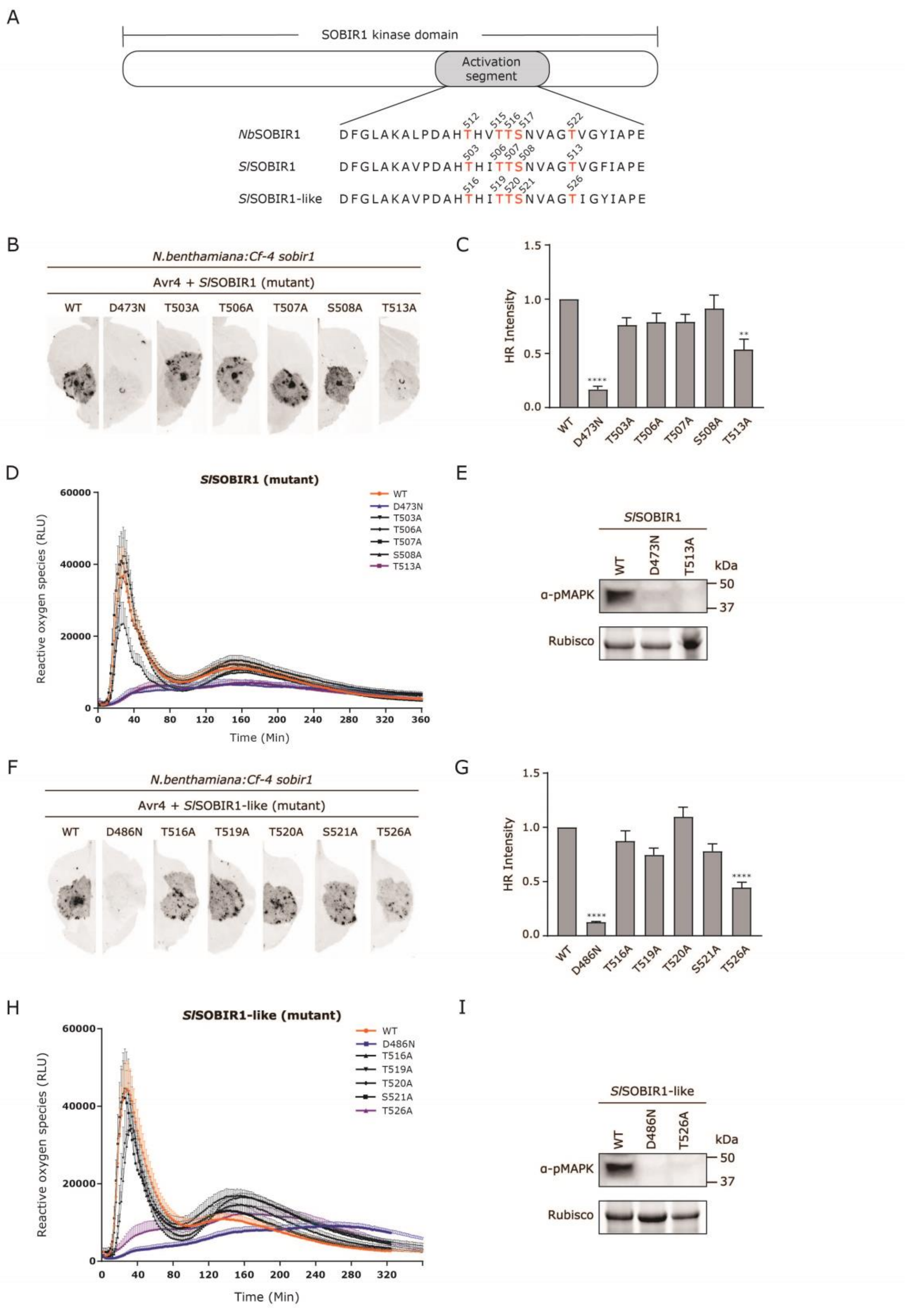

**Figure S2. The analogous residues of *Nb*SOBIR1 Thr522 in both tomato SOBIR1s play a crucial role in mounting Avr4/Cf-4-triggered immune responses. (A)** Schematic diagram of the kinase domain of SOBIR1, with the activation segment indicated. The amino acid sequences of the activation segments of *Nb*SOBIR1, *Sl*SOBIR1 and *Sl*SOBIR1-like are aligned and are shown below the diagram. Conserved residues acting as potential phosphorylation sites are indicated in red. **(B** to

**E)** Mutagenesis screen of five potential phosphorylation sites in the activation segment of *Sl*SOBIR1 based on the Avr4/Cf-4-triggered HR activation **(B** and **C)**, ROS burst **(D)**, and MAPK activation **(E)**. **(F** to **I)** Mutagenesis screen of five potential phosphorylation sites in the activation segment of *Sl*SOBIR1-like based on the Avr4/Cf-4-triggered HR activation **(F** and **G)**, ROS burst **(H)**, and MAPK activation **(I)**. All the generated *Sl*SOBIR1 and *Sl*SOBIR1-like mutants were individually co-expressed with *Avr4* in leaves of *N. benthamiana:Cf-4 sobir1* knock-out plants ($OD_{600}$ = 0.8), with their corresponding WT as a positive control and kinase-dead mutant (*Sl*SOBIR1 D473N or *Sl*SOBIR1-like D486N) as a negative control. The formation of HR was imaged at 5 dpi by Chemidoc **(B** and **F)** and the HR intensity was quantified by Image Lab **(C** and **G).** The HR intensity of leaves that transiently co-express *SlSOBIR1 WT* with *Avr4* **(C)** or *SlSOBIR1-like WT* with *Avr4* **(D)** was set as 1. Statistical analysis was performed with ANOVA/Dunnett's multiple comparison test, compared with their corresponding WT, $^{**}p < 0.01$; $^{****}p < 0.0001$. **(D** and **H)** Leaf discs of *N. benthamiana:Cf-4 sobir1* knock-out plants that transiently express individual *Sl*SOBIR1 and *Sl*SOBIR1-like mutants, as well as their corresponding WTs and kinase-dead mutants, were treated with 0.1 μM Avr4 protein, followed by measuring the production of ROS. ROS production is expressed as RLUs, and the data are represented as mean + SEM. **(E)** *SlSOBIR1 WT*, *T513A* and *D473N* and **(I)** *SlSOBIR1-like WT*, *T526A* and *D486N* were transiently co-expressed with *Avr4* in leaves of *N. benthamiana:Cf-4 sobir1* knock-out plants ($OD_{600}$ = 0.8), leaf samples were collected at 2 dpi, followed by being subjected to a MAPK activation assay. Experiments were repeated at least three times with similar results.

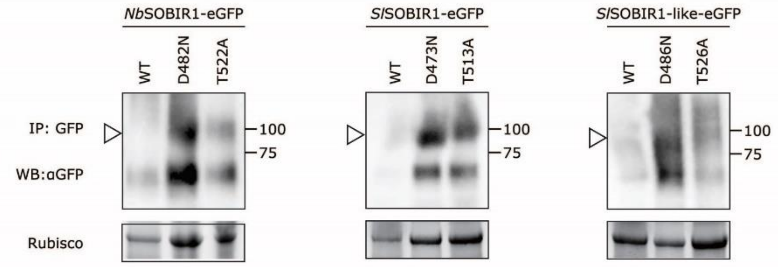

**Figure S3. Accumulation levels of *Nb*SOBIR1 T522A, *Sl*SOBIR1 T513A and *Sl*SOBIR1-like T526A *in planta*, in comparison to their WT and kinase-dead ("RN") versions.** *NbSOBIR1 T522A*, *SlSOBIR1 T513A* and *SlSOBIR1-like T526A*, which fail to restore the Avr4-triggered HR in *N. benthamiana:Cf-4 sobir1* knock-out plants, were transiently co-expressed in *benthamiana:Cf-4 sobir1* plants with *Avr4* (both at an $OD_{600}$ of 0.8). Transient expression of their respective WT and kinase-dead D to N version, in combination with *Avr4*, was included as positive and negative controls, respectively. The leaf samples were collected at 2 dpi. Total protein extracts were subjected to immuno-purification (IP) using GFP-affinity beads, followed by western blotting (WB) with αGFP antibody (upper panels). The amount of total protein that was used for the IP is reflected by the Rubisco band present in the total protein extracts loaded on SDS gel (lower panels). Arrowheads indicate the band corresponding to SOBIR1-eGFP. Note that transient expression of *WT SOBIR1* in combination with *Avr4* results in cell death, thereby causing very low WT SOBIR1 accumulation levels.

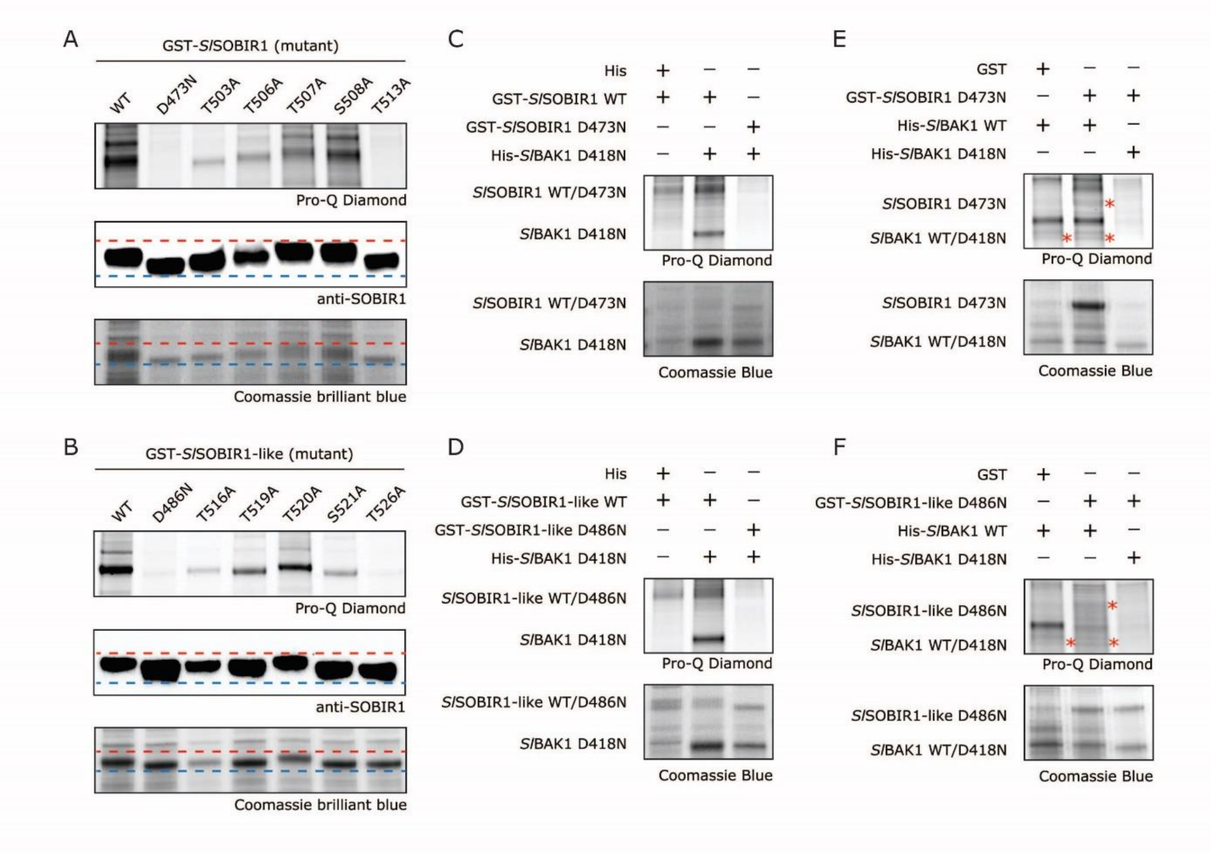

**Figure S4. SOBIR1 and BAK1 trans-phosphorylate each other *in vitro*. (A** and **B)** *Sl*SOBIR1 Thr513 and *Sl*SOBIR1-like Thr526 are essential for their intrinsic kinase activity. The N-terminally GST-tagged cytoplasmic kinase domains of **(A)** *Sl*SOBIR1 WT, kinase-dead mutant D473N, and five Ser/Thr-to Ala mutants, and **(B)** *Sl*SOBIR1-like WT, kinase-dead mutant D486N, and five Ser/Thr-to-Ala mutants were produced in *E. coli*, followed by being subjected to western blotting and *in vitro* phosphorylation assay. The phosphorylation status of the recombinant proteins was determined by using the Pro-Q Diamond stain, which specifically stains the phosphorylated proteins (top panels). The production of SOBIR1 kinase domains was confirmed by western blotting, using SOBIR1 antibodies (middle panels). And all recombinant proteins were stained by Coomassie brilliant blue (bottom panels). **(C** and **D)** *Sl*SOBIR1 WT and *Sl*SOBIR1-like WT directly phosphorylate the kinase-dead mutant of *Sl*BAK1 D418N. The N-terminally GST-tagged cytoplasmic domains of *SlSOBIR1 WT* or *D473N* **(C)** and *SlSOBIR1-like WT* or *D486N* **(D)** were co-expressed with N-terminally His-tagged cytoplasmic domain of *SlBAK1 D418N* in *E. coli*. After the SDS-PAGE of the *E. coli* lysates, the Pro-Q Diamond stain was employed to detect the phosphorylated recombinant proteins (top panels), while Coomassie brilliant blue was used to visualize all proteins (bottom panels). **(E** and **F)** *Sl*BAK1 WT directly phosphorylates the kinase-dead mutant of *Sl*SOBIR1 D473N and *Sl*SOBIR1-like D486N. The cytoplasmic domain of *SlBAK1 WT* or *D418N*, which was fused to a His tag in its N-terminus, was co-expressed with the cytoplasmic domain of *SlSOBIR1 D473N* **(E)** or *SlSOBIR1-like D486N* **(F)**, which was fused to a GST tag in its N-terminus, in *E. coli*. The recombinant proteins were then subjected to SDS-PAGE, followed by being stained by Pro-Q Diamond stain (top panels) and Coomassie brilliant blue (bottom panels). Bands with the expected sizes are indicated with an asterisk. Experiments were repeated at least three times with similar results, and representative pictures are shown.

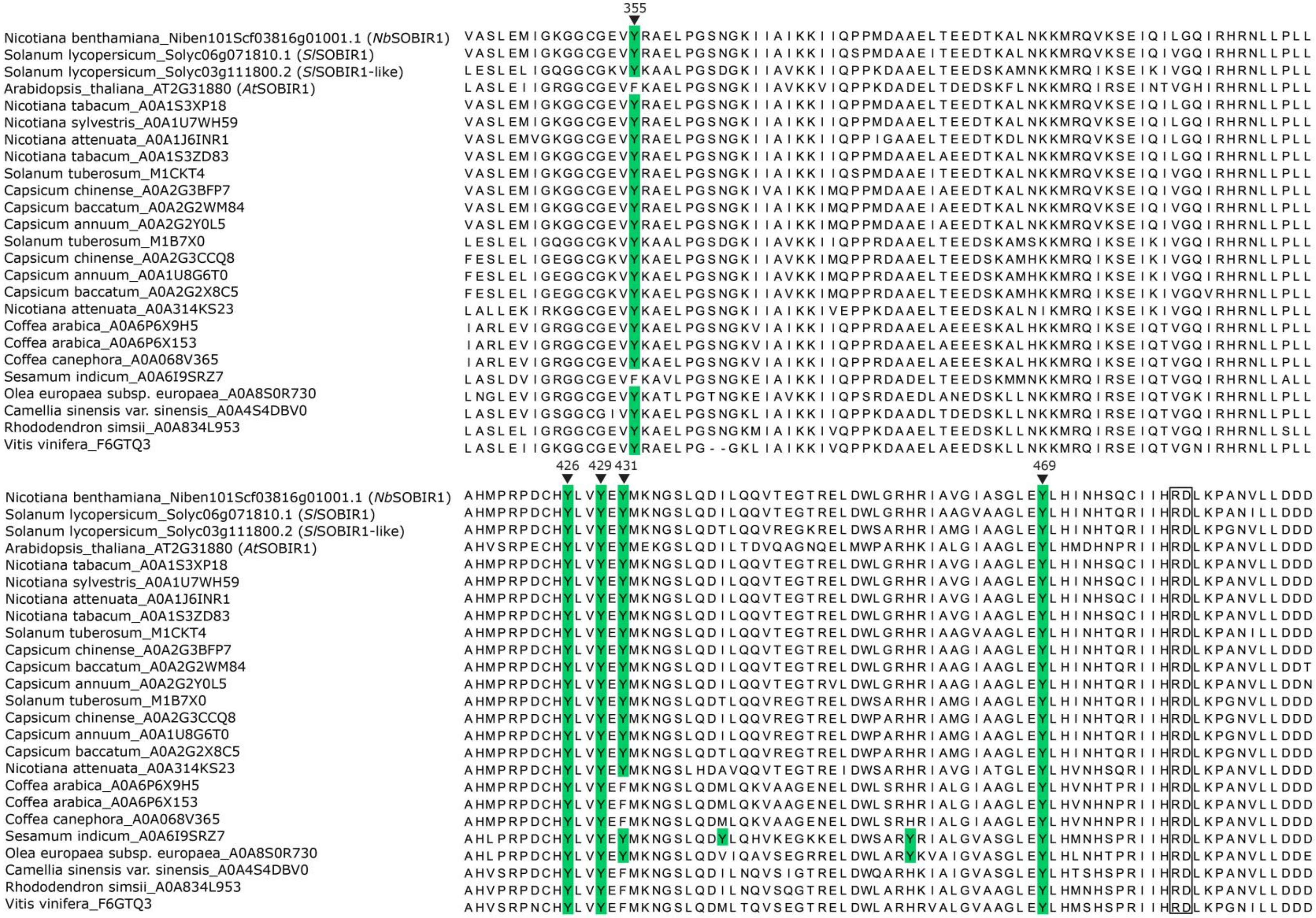
355
Nicotiana benthamiana_Niben101Scf03816g01001.1 (NbSOBIR1)
Solanum lycopersicum_Solyc06g071810.1 (SlSOBIR1)
Solanum lycopersicum_Solyc03g111800.2 (SlSOBIR1-like)
Arabidopsis_thaliana_AT2G31880 (AtSOBIR1)
Nicotiana tabacum_A0A1S3XP18
Nicotiana sylvestris_A0A1U7WH59
Nicotiana attenuata_A0A1J6INR1
Nicotiana tabacum_A0A1S3ZD83
Solanum tuberosum_M1CKT4
Capsicum chinense_A0A2G3BFP7
Capsicum baccatum_A0A2G2WM84
Capsicum annuum_A0A2G2Y0L5
Solanum tuberosum_M1B7X0
Capsicum chinense_A0A2G3CCQ8
Capsicum annuum_A0A1U8G6T0
Capsicum baccatum_A0A2G2X8C5
Nicotiana attenuata_A0A314KS23
Coffea arabica_A0A6P6X9H5
Coffea arabica_A0A6P6X153
Coffea canephora_A0A068V365
Sesamum indicum_A0A6I9SRZ7
Olea europaea subsp. europaea_A0A8S0R730
Camellia sinensis var. sinensis_A0A4S4DBV0
Rhododendron simsii_A0A834L953
Vitis vinifera_F6GTQ3
426
429
431
469

525 530 543

| | |
|---|---|
| Nicotiana benthamiana_Niben101Scf03816g01001.1 (*Nb*SOBIR1) | MEARIADFGLAKALPDAHTHVTTSNVAGTVGYIAPEYHQTLKFTGKCDIYSFGVVLAVLVIGKLPSDEFFQHTPEMS |
| Solanum lycopersicum_Solyc06g071810.1 (*Sl*SOBIR1) | MEARVADFGLAKAVPDAHTHITTSNVAGTVGFIAPEYYQTLKFTDKCDIYSFGVVLAVLVIGKGPSDDFFQHTSEMS |
| Solanum lycopersicum_Solyc03g111800.2 (*Sl*SOBIR1-like) | MEARIADFGLAKAVPDAHTHITTSNVAGTIGYIAPEYHQTLKFTDKCDIYSFGVLLGVLVMGKLPSDEFFQNTSEMS |
| Arabidopsis_thaliana_AT2G31880 (*At*SOBIR1) | MEARISDFGLAKAMPDAVTHITTSHVAGTVGYIAPEFYQTHKFTDKCDIYSFGVILGILVIGKLPSDEFFQHTDEMS |
| Nicotiana tabacum_A0A1S3XP18 | MEARIADFGLAKALPDAHTHITTSNVAGTVGYIAPEYHQTLKFTDKCDIYSFGVVLAVLVIGKLPSDEFFQHTPEMS |
| Nicotiana sylvestris_A0A1U7WH59 | MEARIADFGLAKALPDAHTHITTSNVAGTVGYIAPEYHQTLKFTDKCDIYSFGVVLAVLVIGKLPSDEFFQHTPEMS |
| Nicotiana attenuata_A0A1J6INR1 | MEARIADFGLAKALPDAHTHITTSNVAGTVGYIAPEYHQTLKFTDKCDIYSFGVVLAVLVIGKLPSDEFFQHTPEMS |
| Nicotiana tabacum_A0A1S3ZD83 | MEARIADFGLAKAVPDAHTHVTTSNVAGTVGYIAPEYHQTLKFTDKCDIYSFGVVLAVLVIGKLPSDEFFQHTPEMS |
| Solanum tuberosum_M1CKT4 | MEARIADFGLAKAVPDAHTHITTSNVAGTMGYIAPEYYQTLKFTDKCDIYSFGVVLAVLVIGKGPSDEYFQHTSEMS |
| Capsicum chinense_A0A2G3BFP7 | MEPRIADFGLAKAVPDAHTHVTTSNVAGTVGYIAPEYHQTLKFTDKCDIYSFGVVLAVLVVGKLPSDDFFQHTSEMS |
| Capsicum baccatum_A0A2G2WM84 | MEARIADFGLAKAVPDAHTHVTTSNVAGTVGYIAPEYHQTLKFTDKCDIYSFGVVLAVLVIGKLPSDDFFQHTSEMS |
| Capsicum annuum_A0A2G2Y0L5 | MEPRIADFGLAKAVPDAHTHVTTSNVAGTVGYIAPEYHQTLKFTDKCDIYSFGVVLAVLVVGKLPSDDFFQHTSEMS |
| Solanum tuberosum_M1B7X0 | MEARIADFGLAKAVPDAHTHITTSNVAGTIGYIAPEYHQTLKFTDKCDIYSFGVLLGVLVMGKLPSDEFFQNTSEMS |
| Capsicum chinense_A0A2G3CCQ8 | LEARIADFGLAKAVPDAHTHITTSNVAGTIGYIAPEYHQTLKFTDKCDIYSFGVLLGVLVMGKLPSDEFFQTTSEMS |
| Capsicum annuum_A0A1U8G6T0 | LEGRIADFGLAKAVPDAHTHITTSNVAGTIGYIAPEYYQTLKFTDKCDIYSFGVLLGVLVMGKLPSDEFFQTTSEMS |
| Capsicum baccatum_A0A2G2X8C5 | LEPRIADFGLAKAVPDAHTHITTSNVAGTIGYIAPEYHQTLKFTDKCDIYSFGVLLGVLVMGKLPSDEFFQTTSEMS |
| Nicotiana attenuata_A0A314KS23 | MEARIADFGLAKAIPESLTHVSTSHVVGTLGYIAPAYYQTVKFTDKCDIYSFGVLLGVLVMGKFPSDELFQPASGMG |
| Coffea arabica_A0A6P6X9H5 | MEARIADFGLAKAMPEAYTHVTSSNVVGTLGYIAPEYHQTLKFTDKCDIYSFGVLLASLVMGKLPSDEFFQETDEMN |
| Coffea arabica_A0A6P6X153 | MEARIADFGLAKAMPEAYTHVTSSNVVGTLGYIAPEYHQTLKFTDKCDIYSFGVLLASLVMGKLPSDEFFQETDEMN |
| Coffea canephora_A0A068V365 | MEARIADFGLAKAMPEAYTHVTSSNVVGTLGYIAPEYHQTLKFTDKCDIYSFGVLLASLVMGKLPSDEFFQETDEMN |
| Sesamum indicum_A0A6I9SRZ7 | MEARIADFGLAKAMPDANTHVTTSNVAGTVGYIAPEYYQTFKFTDKCDIYSFGVVLGVLVMGKLPSDDFFQHTDELS |
| Olea europaea subsp. europaea_A0A8S0R730 | MEARIADFGLAKAVPEANTHVSTSNVAGTAGYIAPEYHQTFKFTEKCDIYSFGVVLGALTMGKFPSDEFFQHTEEMH |
| Camellia sinensis var. sinensis_A0A4S4DBV0 | MEARIADFGLAKAVPDAHTHVTTSNVAGTVGYIAPEYHQTLKFTDKCDIFSFGVLLGVLVIGKLPSDDFFQHTSEMS |
| Rhododendron simsii_A0A834L953 | MEARIADFGLAKAVPDQDTHVTTSNVAGTLGYIAPEYHQTMKFTDKCDIYSFGVLLGVLVIGRLPSDNFFQDTSEMS |
| Vitis vinifera_F6GTQ3 | MEARIADFGLAKAVPDANTHVTTSNVAGTVGYIAPEYHQTLKFTDKCDIYSFGVLLGVLVVGKLPSDDFFQHTAEMS |

| | |
|---|---|
| Nicotiana benthamiana_Niben101Scf03816g01001.1 (*Nb*SOBIR1) | LVKWLRNVMTSEDPKRAIDSKLIGNGFEEQMLLVLKIACFCTLENPKERPNSKDVRCMLTQIKH------ |
| Solanum lycopersicum_Solyc06g071810.1 (*Sl*SOBIR1) | LVKWLRNVMTSDDPKIAIDPKLIGNGYDEQMLLVLKIACFCTLDNPKERPNSKDVRCMLTQIKH------ |
| Solanum lycopersicum_Solyc03g111800.2 (*Sl*SOBIR1-like) | LVKWMRNVMTSEDPNRAIDPKLMGNGNEDQMLLVLKIACFCTMENPKERPNSKDVRCMLMQIKH------ |
| Arabidopsis_thaliana_AT2G31880 (*At*SOBIR1) | LIKWMRNIITSENPSLAIDPKLMDQGFDEQMLLVLKIACYCTLDDPKQRPNSKDVRTMLSQIKH------ |
| Nicotiana tabacum_A0A1S3XP18 | LVKWLRNVMTSEDPKRAIDPKLIGTGFEEQMLLVLKIACFCTLENPKERPNSKDVRCMLTQIKH------ |
| Nicotiana sylvestris_A0A1U7WH59 | LVKWLKNVMTSEDPKRAIDPKLIGTGFEEQMLLVLKIACFCTLENPKERPNSKDVRCMLTQIKH------ |
| Nicotiana attenuata_A0A1J6INR1 | LVKWLRNVMTSEDPKRAIDQKLIGNGFEEQMLLVLKIACFCTLENPKERPNSKDVRCMLTQIKH------ |
| Nicotiana tabacum_A0A1S3ZD83 | LVKWLRNVMTSEDPKRAIDPKLIGSGFEEQMLLVLKIACFCTLENPKERPNSKDVRCMLTQIKH------ |
| Solanum tuberosum_M1CKT4 | LVKWLRNVMTSDDPKIAIDPKLRGNGYEEQMLLVLKIACFCTLDNPKERPNSKDVRCMLTQIKH------ |
| Capsicum chinense_A0A2G3BFP7 | LVKWLRNVMTSDDPKRAIDPNLIGNGYEEQMLLVLKIACFCTMDNPKERPNSKDVRCMLTQIKH------ |
| Capsicum baccatum_A0A2G2WM84 | LVKWLRNVMTSDDPKRAIDPNLMGNGYEEQMLLVLKIACFCTMDNPKERPNSKDVRCMLTQIKH------ |
| Capsicum annuum_A0A2G2Y0L5 | LVKWLRNVMTSDDPKRAIDPNLIGNGYEEQMLLVLKIACFCTMDNPKERPNSKDVRCMLTQIKH------ |
| Solanum tuberosum_M1B7X0 | LVKWMRNVMTSEDPNRAIDPKLMGNGNEDQMLLVLKIACFCTLENPKERPNSKDVRCMLMQIKH------ |
| Capsicum chinense_A0A2G3CCQ8 | LVKWMRNVMTSEDPNRAIDPKLMGNGNEEQMLLVLKIACFCTLENPKERPDSKDVRCMLMQIKH------ |
| Capsicum annuum_A0A1U8G6T0 | LVKWMRNVMTSEDPNRAIDPKLMGNGNEEQMLLVLKIACFCTLENPKERPNSKEVRCMLMQIKH------ |
| Capsicum baccatum_A0A2G2X8C5 | LVKWMRNVMTSEDPNRAIDPKLMGNGNEEQMLLVLKIACFCTLENPKERPNSKEARCMLMQIKH------ |
| Nicotiana attenuata_A0A314KS23 | LVKWMRNVMTSENPKRAIDPKLMGNGYEEQMLLVLKIACFCTLDNPKERPNSKDVRCMLTQIKP------ |
| Coffea arabica_A0A6P6X9H5 | LVHWMRNVMTSEDPKRAIDPKLLGNGYEEQMLLVLKIACFCTLENPKERPNSIDIRAMLFQIKYEKRQVM |
| Coffea arabica_A0A6P6X153 | LVLWMRNVMTSEDPKRAIDPKLLGNGYEEQMLLVLKIACFCTLENPKERPNSIDIRAMLFQIKYEKR--- |
| Coffea canephora_A0A068V365 | LVLWMRNVMTSEDPKRAIDPKLLGNGYEEQMLLVLKIACFCTLENPKERPNSIDIRAMLFQIKYEKR--- |
| Sesamum indicum_A0A6I9SRZ7 | LVKWMRNVMTSEDPKRAIDPKLLGNGYEEQMLLVLKVACFCTLDNPKERPNSKDARCMLSQIRH------ |
| Olea europaea subsp. europaea_A0A8S0R730 | LVKWMRNVMTSEDPKRAIDPRLLGNGYEEQILLVLKIACFCTLDNPKERPDSKEIRCMLTQIKH------ |
| Camellia sinensis var. sinensis_A0A4S4DBV0 | LVKWMRNVMTSEDPTRAIDPRLMGNGDHENMLLVLKIACFCTLDDPKQRPNSKDVRCMLSQIKN------ |
| Rhododendron simsii_A0A834L953 | LVKWMRNVMTSEDPNRAIDRKLIGNGHEEKMLLVLKIACFCTLEDPKQRPNSKDVRCMLDQITH------ |
| Vitis vinifera_F6GTQ3 | LVKWMANIRTSDDPSRAIDRKLMGNGFEEQMLLVLKIACFCTVDDAKIRPNSKDVRTMLSQIKH------ |

**Figure S5. Alignment of the protein sequences of the SOBIR1 kinase domain from various plant species.** Amino acid sequences of the SOBIR1 homologs from various plant species were obtained from the UniProt database (https://www.uniprot.org/). The alignment was visualized using JalView and only the amino acid sequences of the kinase domain are shown, with all the Tyr (Y) residues highlighted in green. The Tyr residues that are subjected to this study are marked with arrowheads on the top, and their position in *Nb*SOBIR1 is indicated on the top. The RD motif (in which a conserved arginine (R) precedes the highly conserved catalytic aspartate (D)), present in all SOBIR1 kinase domains, is indicated by a box.

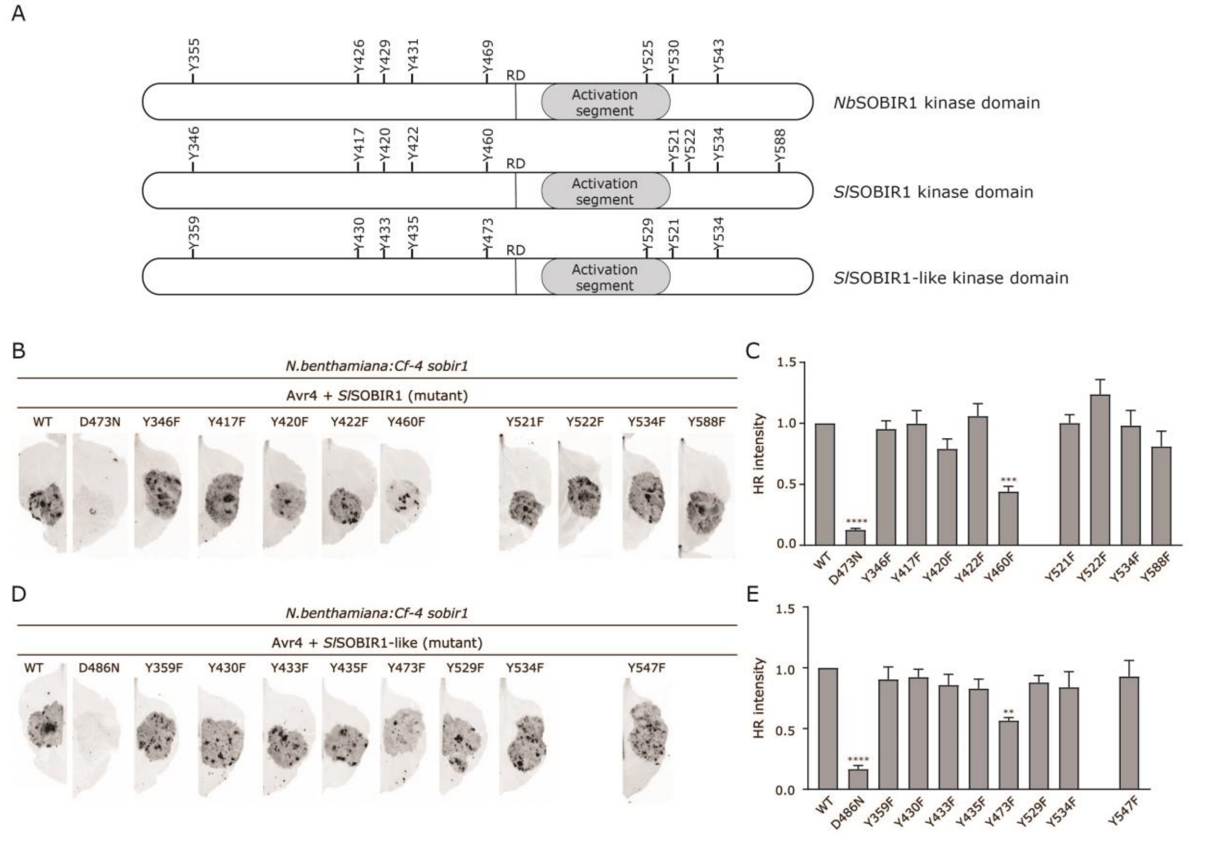

**Figure S6. *Sl*SOBIR1 Tyr460 and *Sl*SOBIR1-like Tyr473, which are the analogous residues of *Nb*SOBIR1 Tyr469, are essential for the Avr4/Cf-4-triggered HR in *N. benthamiana*. (A)** Schematic diagrams of the kinase domains of *Nb*SOBIR1, *Sl*SOBIR1 and *Sl*SOBIR1-like, with the location of the activation segment, the RD motif, and all Tyr residues indicated. **(B** to **E)** Mutagenesis screen of all putative Tyr phosphorylation sites in *Sl*SOBIR1 **(B** and **C)** and *Sl*SOBIR1-like **(D** and **E)**, as described in Figure 3, to determine their importance in immune signaling by complementation. Statistical analysis was performed with ANOVA/Dunnett's multiple comparison test, compared with their corresponding WT. $^{**}p < 0.01$; $^{***}p < 0.001$; $^{****}p < 0.0001$. Experiments were repeated at least three times with similar results, and representative pictures are shown.

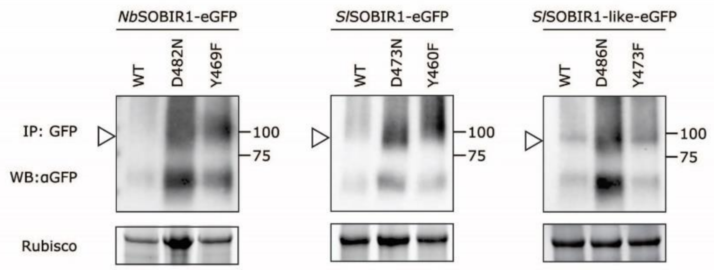

**Figure S7. Accumulation levels of *Nb*SOBIR1 Y469F, *Sl*SOBIR1 Y460F and *Sl*SOBIR1-like Y473F, as well as their corresponding WT and kinase-dead (D to N) versions, *in planta*.** *NbSOBIR1 Y469F*, *SlSOBIR1 Y460F* and *SlSOBIR1-like Y473F*, which fail to fully restore the Avr4-triggered HR in *N. benthamiana:Cf-4 sobir1* knock-out plants, were transiently expressed in *N. benthamiana:Cf-4 sobir1* plants in combination with *Avr4* (both at an $OD_{600}$ of 0.8), next to their respective *WT* that was combined with *Avr4* as a positive control, and their kinase-dead D to N version that was combined with *Avr4* as a negative control. Leaf samples were collected at 2 dpi. Total protein extracts were subjected to IP of the GFP-tagged SOBIR1 mutants using GFP-affinity beads, followed by WB with αGFP antibody (upper panels). The amount of total protein that was used for the IP is reflected by the Rubisco band present in the stain-free gel (lower panels). Arrowheads indicate the band representing SOBIR1-eGFP. Note that transient expression of *SOBIR1 WT* in combination with *Avr4* triggers an HR in *N. benthamiana:Cf-4 sobir1* plants, which explains the low accumulation levels of SOBIR1 WT.

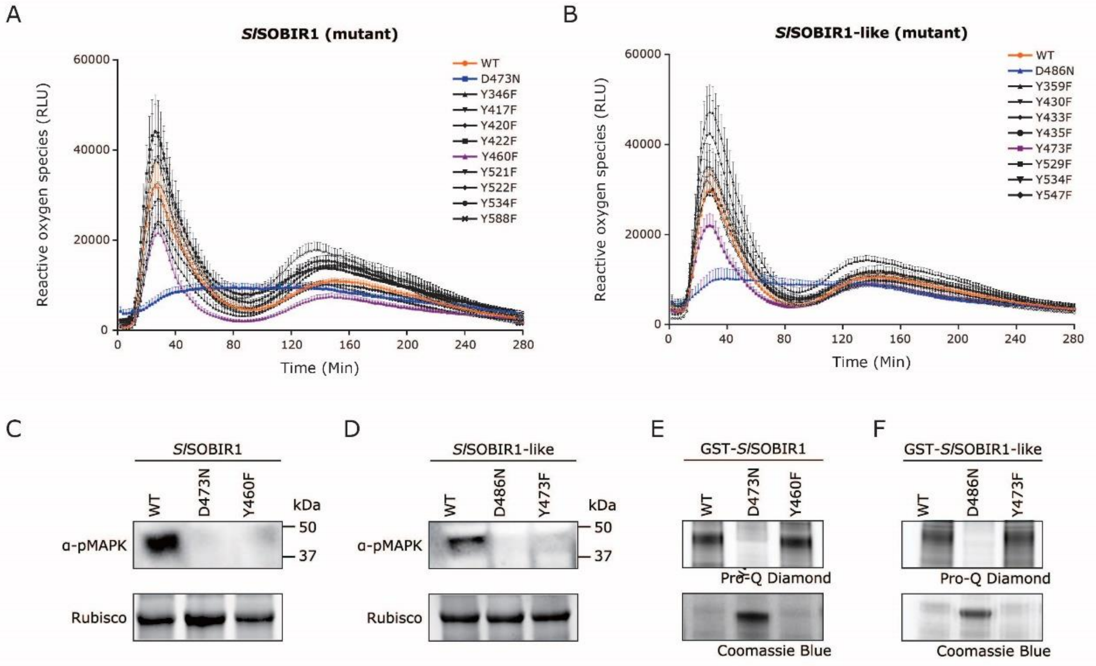

**Figure S8. *Sl*SOBIR1 Tyr460 and *Sl*SOBIR1-like Tyr473 are required for the Avr4/Cf-4-induced MAPK activation, but not for ROS accumulation and their intrinsic kinase activity. (A and B)** The different Tyr mutants of *Sl*SOBIR1 and *Sl*SOBIR1-like were transiently expressed in leaves of the *N. benthamiana:Cf-4 sobir1* knock-out line, with their corresponding WTs as positive controls and kinase-dead mutants as negative controls. Leaf discs were taken from these plants 24 hours after agro-infiltration, followed by adding 0.1 µM Avr4 protein and measuring ROS accumulation over time. ROS production is expressed as RLUs, and the data are represented as mean + SEM. **(C and D)** *SlSOBIR1 Y460F* and *SlSOBIR1-like Y473F*, as well as their corresponding WTs and kinase-dead mutants, were transiently co-expressed with *Avr4* in leaves of *N. benthamiana:Cf-4 sobir1* knock-out plants. The leaf samples were collected at 2 dpi. Hereafter, total protein extracts were subjected to immunoblotting with a p42/p44-erk antibody to determine the activation of downstream MAPKs by phosphorylation. **(E and F)** The N-terminally GST-tagged cytoplasmic kinase domains of *Sl*SOBIR1 Y460F and *Sl*SOBIR1-like Y473F were produced in *E. coli*, with their corresponding WTs as positive controls and their kinase-dead mutants as negative controls. After SDS-PAGE of the *E. coli* lysates, the recombinant proteins were stained with Coomassie brilliant blue (lower panels), whereas the phosphorylation status of the kinase domains was determined by performing a Pro-Q Diamond stain (upper panels). Experiments were repeated at least three times and similar results were obtained. Representative pictures are shown.

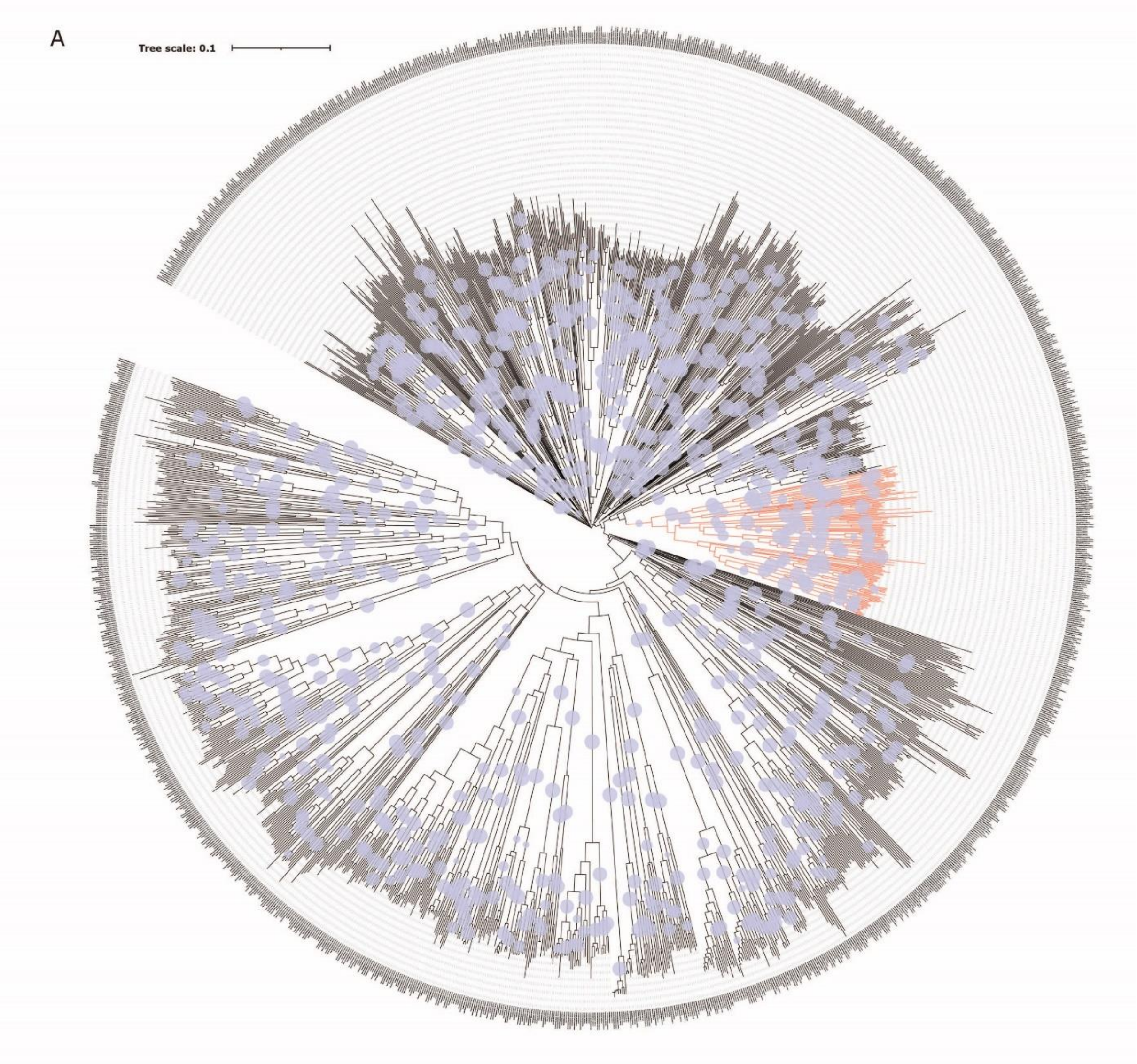
A
Tree scale: 0.1

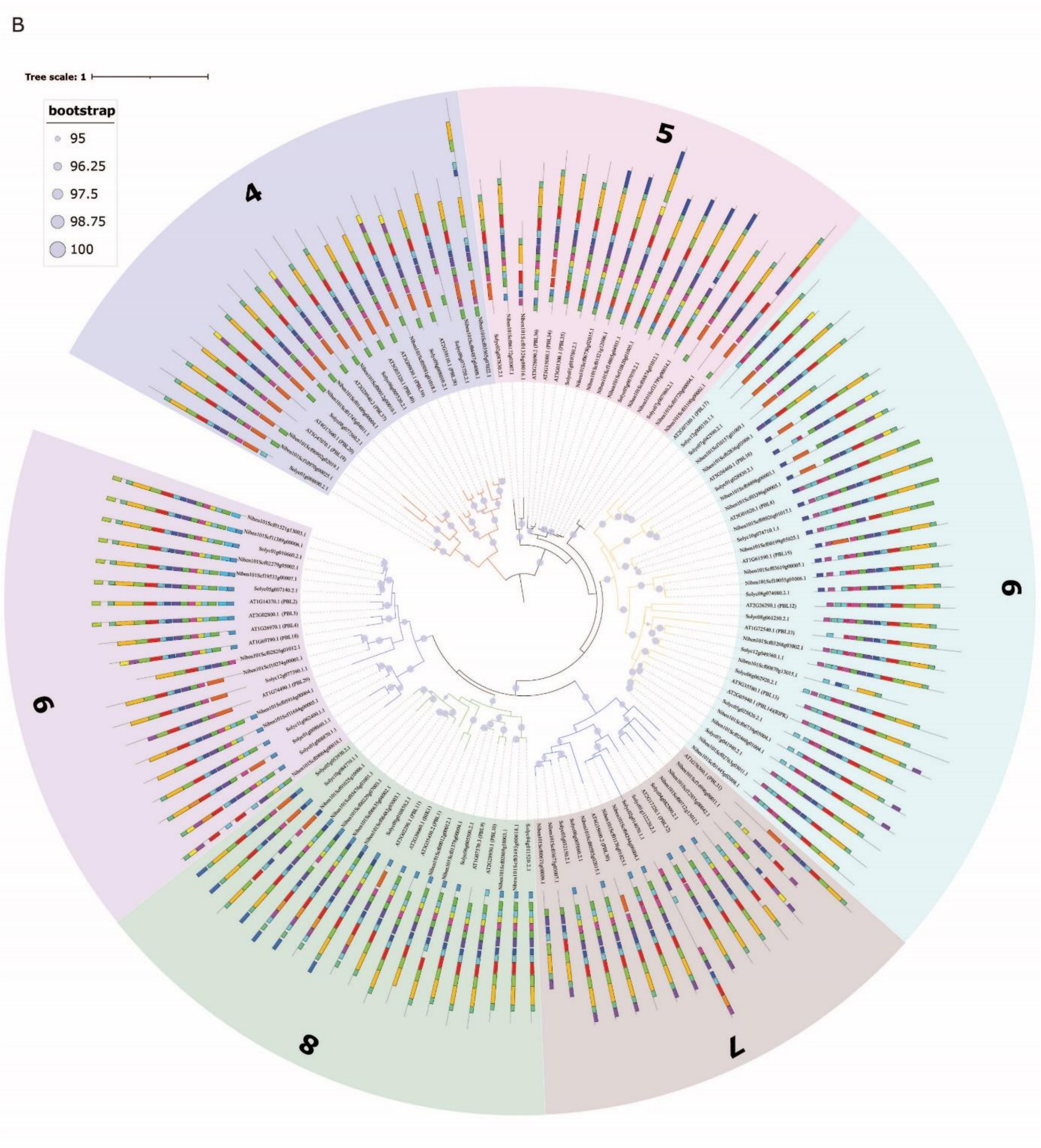

**Figure S9. Phylogenetic analysis of BIK1 homologs from Arabidopsis, tomato, and *N. benthamiana*. (A)** The amino acid sequences of only the kinase domain were extracted from all RLCK members from Arabidopsis, tomato, and *N. benthamiana* and aligned to subsequently generate a neighbor-joining phylogenetic tree using QuickTree (Howe et al., 2002). The sub-clade of putative BIK1 homologs, which comprised 123 sequences including *At*BIK1 (bootstrap support higher than 90%) is shown in red. **(B)** Phylogenetic analysis of the RLCK-VII subfamily members from Arabidopsis, tomato, and *N. benthamiana*. Amino acid motifs identified in the complete protein sequence using MEME are shown (Bailey et al., 2009). All the members present in this tree were further assigned to 6 subfamilies, which are depicted in different colors. These subfamilies are referred to as subfamily 4, 5, 6, 7, 8, and 9, according to the RLCK-VII subfamilies in Arabidopsis reported previously by Rao et al. (2018).

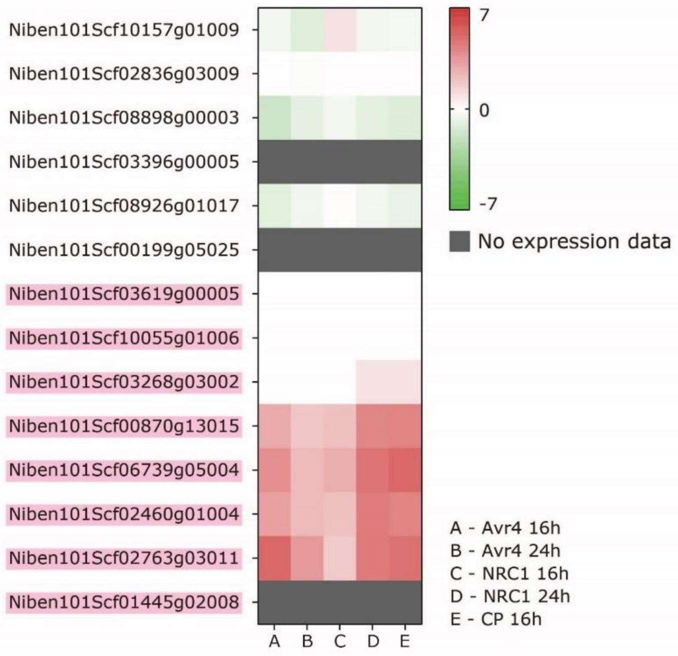

**Figure S10. Gene expression of the 14 members from *N. benthamiana* RLCK-VII-6.** Heat map of the relative expression (log2) of all 14 RLCK members of subfamily 6, as determined upon transient expression of *Avr4*, constitutively active *NRC1* (Gabriëls et al., 2007), or *CP* of potato virus X (Tameling et al., 2010), in leaves of both *Cf-4*- and *Rx*-transgenic *N. benthamiana* plants. The genes that were selected to be knocked out are highlighted in pink. NRC1, NB-LRR PROTEIN REQUIRED FOR HR-ASSOCIATED CELL DEATH 1; CP, coat protein.

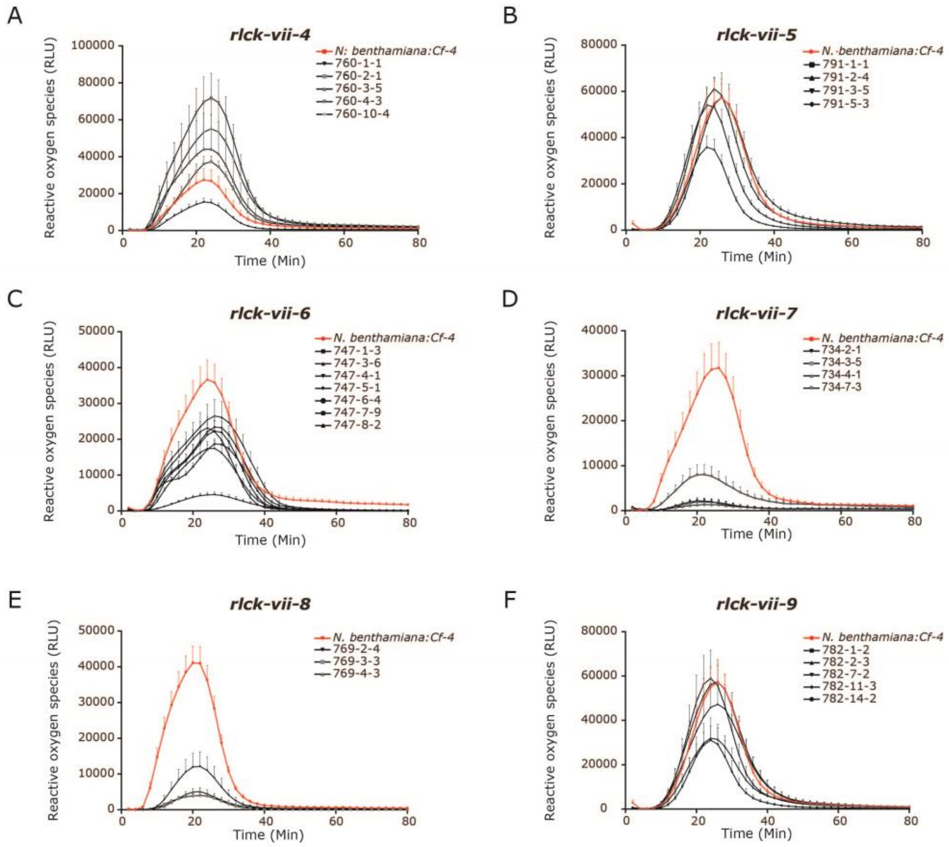

**Figure S11. RLCK-VII-6, -7 and -8 from *N. benthamiana* also play a positive role in the flg22/FLS2-triggered ROS burst.** ROS production, triggered upon treatment with flg22, by discs taken from leaves of *rlck-vii-4* **(A)**, *rlck-vii-5* **(B)**, *rlck-vii-6* **(C)**, *rlck-vii-7* **(D)**, *rlck-vii-8* **(E)** and *rlck-vii-9* **(F)** *N. benthamiana:Cf-4* knock-out plants from the T1 generation, was measured. For this, leaf discs were taken from the different knock-out plants, as well as from *N. benthamiana:Cf-4* (the positive control), followed by treatment with a final concentration of 0.1 µM flg22 peptide and subsequent monitoring of the accumulation of ROS. ROS production is expressed as RLUs, and the data are represented as mean plus the standard error of the mean (SEM) (n≥6). The ROS traces of the positive control are indicated in red in all the line charts. All experiments were repeated at least three times and data from one representative experiment are shown.

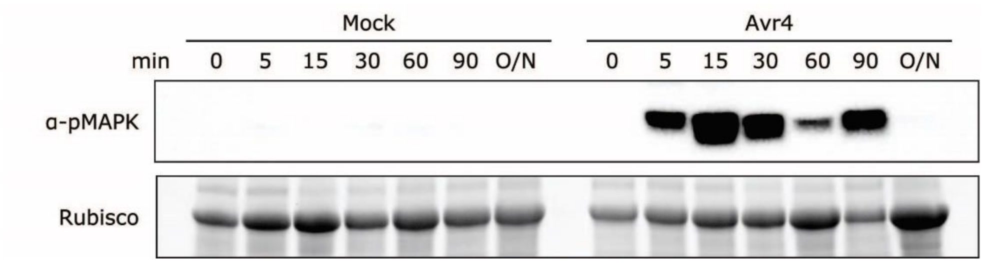

**Figure S12. The Avr4 protein triggers a swift MAPK activation in *N. benthamiana:Cf-4*.** Water (mock) or 5 μM of the pure Avr4 protein was infiltrated in leaves of *N. benthamiana:Cf-4*. Leaf samples were taken at the indicated time points after Avr4 infiltration, and total protein extracts were subjected to immunoblotting using a p42/p44-erk antibody specifically detecting MAPKs that are activated by phosphorylation (α-pMAPK). Rubisco is shown as a total protein loading control. O/N, overnight.

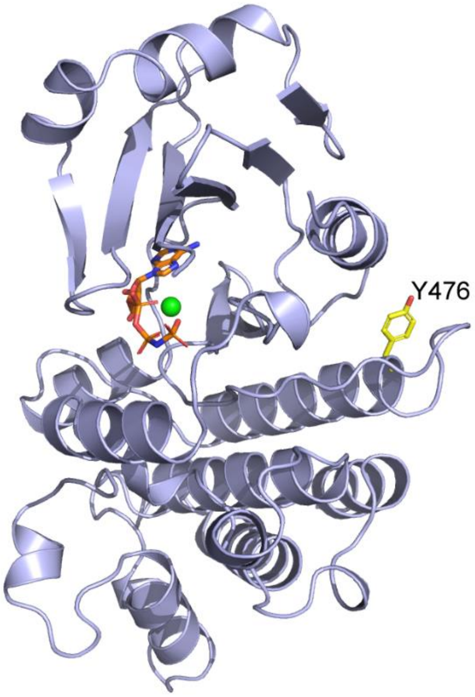

**Figure S13. The overall structure of the inactive *At*SOBIR1 kinase domain and position of the important Tyr residues.** The ribbon diagram of the SOBIR1 kinase domain is colored in light blue. The nonhydrolyzable ATP analog AMP-PNP and $Mg^{2+}$ are presented as an orange stick and a green sphere, respectively. *At*SOBIR1 Tyr476, which is analogous to *Nb*SOBIR1 Tyr469, is indicated as a yellow stick.

**Table S1.** ***RLCK*** **genes selected to be targeted by sgRNAs.**

| Subfamily | Gene name | Amount of sgRNAs | Target sequence (5′ - 3′) | Note |
|---|---|---|---|---|
| 4 | *Niben101Scf10970g00025* | 2 | TTACTTAAGATTGGAGAAGGAGG; ATGGGGAAAGAGGTATTCAGCGG | |
| | *Niben101Scf06902g02019* | | | |
| | *Niben101Scf01745g04011* | 3 | ATTAAGAAGCTCAATACACTTGG; GGAATGGCTTATTTGCACGACGG | |
| | *Niben101Scf01489g00004* | | | |
| | *Niben101Scf01071g03021* | | | Not present in the phylogeny, but clustered to this subfamily according to iTAK* |
| | *Niben101Scf00012g00010* | 2 | GTAGTGCCAGCGTTGATACGCGG; GTGCGGAGGATGATGAAAGGGGG | |
| | *Niben101Scf03565g07022* | 2 | GGATCGGCTAAAGTTCCTGGTGG; GCTCATAGGTTATTGCGCACAGG | |
| | *Niben101Scf04487g04008* | | | |
| | *Niben101Scf05081g01018* | | GTTCTACGTTGGCGACAAGAAGG; GATACCAGCACAGAGTCCAGAGG | |
| 5 | *Niben101Scf06172g03007* | 2 | GCTTTGGCCCTGTTTACAAAGGG; CTAATTTAACCAAATTCTGATGG | |
| | *Niben101Scf01326g08016* | | | Not expressed according to QUT |
| | *Niben101Scf06750g02015* | 2 | CCAGAATCCCTTCTTGGTGAAGG; ACCGCATCTAGCAGAATGTTGG | |
| | *Niben101Scf01521g12006* | | | |
| | *Niben101Scf08874g01012* | | | |
| | *Niben101Scf11795g00014* | | | |
| | *Niben101Scf14805g04011* | 2 | TGGAGTCTTGTGAATCAAAGGGG; GTTACTTTCCGCATTACTTGTGG | |
| | *Niben101Scf10820g01005* | | | |
| | *Niben101Scf05720g00004* | 2 | AAGAGCTAAAAGCTGCAACGGGG; GTTTATGAGTTCATGACCCGTGG | |
| | *Niben101Scf03100g00002* | | | |
| 6 | *Niben101Scf10157g01009* | | | Not selected, see Figure S10 |
| | *Niben101Scf02836g03009* | | | Not selected, see Figure S10 |
| | *Niben101Scf08898g00003* | | | Not selected, see Figure S10 |
| | *Niben101Scf03396g00005* | | | Not selected, see Figure S10 |
| | *Niben101Scf08926g01017* | | | Not selected, see Figure S10 |

| | | | | |
|---|---|---|---|---|
| | *Niben101Scf00199g05025* | | | Not selected, see Figure S10 |
| | *Niben101Scf03619g00005* | 2 | AATTACTTGCTTGGTGAAGGTGG; ATTGATTGGGTACTGTTGCGAGG | |
| | *Niben101Scf10055g01006* | | | |
| | *Niben101Scf03268g03002* | 1 | GGGTTTGGTCCTGTGCATAAGGG | |
| | *Niben101Scf00870g13015* | 2 | TTATCGTCGAACGCGATCATCGG; GATATTCAAGTTGCTTGCCATGG | |
| | *Niben101Scf06739g05004* | 2 | GTAATTTCTTGGGTGAAGGAGG; ATTGTTGTGAAGAGGAACACAGG | |
| | *Niben101Scf02460g01004* | | | |
| | *Niben101Scf02763g03011* | 2 | GGGTTTGGACCAGTTCATAAGGG; GTATATGAGTACATGCCCAGAGG | |
| | *Niben101Scf01445g02008* | | | |
| 7 | *Niben101Scf14996g00011* | | | Not expressed according to QUT |
| | *Niben101Scf12935g00042* | | | Not expressed according to QUT |
| | *Niben101Scf00712g13012* | 2 | GAGCTAGGAAGATTATCAACAGG; CCTAGTGGAGATGGAGGAGGTGG | |
| | *Niben101Scf04294g06004* | 2 | AAGACTTTCTCATCCTAACCTGG; CTTGCTTCTCCGATGCATGTAGG | |
| | *Niben101Scf01176g01025* | | | |
| | *Niben101Scf00585g02015* | 2 | CGCTCGCGGTAGCTGAAAAACGG; CTTGACACAAAACGACCTAGTGG | |
| | *Niben101Scf03673g03007* | 2 | ATTTATGCAGAAGGGAAGCTTGG; AGCCGCAAGTCCCAAGAAAGTGG | |
| | *Niben101Scf00671g00009* | | | |
| | *Niben101Scf02413g03005* | 1 | CTTGACACAAAACGACCTAGTGG | Not present in the phylogeny, but clustered to this subfamily according to iTAK |
| 8 | *Niben101Scf03493g00018* | 2 | ATGAACTCCGAGTAGCCACGAGG; GCTATAACCAAACCAGTTCCCGG | |
| | *Niben101Scf02869g18003* | | | |
| | *Niben101Scf01378g00004* | 2 | ATCGGGACGAAAATTCCTAGTGG; GATAGCAATCCAAGCTCGAGGG | |
| | *Niben101Scf00012g00012* | | | |
| | *Niben101Scf06482g03003* | 3 | GTCAGGACGGAAGTTTCTAGTGG; GTAGTAGGAGAAGGAGGTTTTGG | |
| | *Niben101Scf00635g04002* | | | |
| | *Niben101Scf00229g07003* | | | |
| | *Niben101Scf05476g01001* | 2 | TTAAACCAAGAAGGGTGGCAGGG; | |

| | | | |
|---|---|---|---|
| | *Niben101Scf01025 g10006* | | CTCCTACAACACTATCA GGACGG |
| 9 | *Niben101Scf09004 g00018* | 2 | GTTTATGAGGTCATGAC AAGAGG; GCCCATAGTACCTATAA CCCTGG |
| | *Niben101Scf11684 g00005* | 2 | TTAAGCCAGAAAGCTTT CAGGGG; GTATATGACCAAAGGAA GCTTGG |
| | *Niben101Scf05916 g00004* | | |
| | *Niben101Scf10274 g00001* | 2 | TTATGGCCCTGCAAACC TCGAGG; GTATATGAATTTATGCCA AGAGG |
| | *Niben101Scf02826 g01012* | | |
| | *Niben101Scf19533 g00007* | 2 | GTCAAGAAATTGAAGCC GGAAGG; GTGTATGAGTTCATGCC TAAAGG |
| | *Niben101Scf02279 g05002* | | |
| | *Niben101Scf11389 g00006* | 3 | GGTTCTGCTGGGAATCC TAGAGG; GCAACTTCGTCATCCAA ACCTGG; GTGTATGAGTTCATGCC TAAAGG |
| | *Niben101Scf01521 g13003* | | |

*iTAK (http://itak.feilab.net/); QUT (http://www.benthgenome.qut.edu.au/). The genes that are not expressed according to the QUT genome browser were not chosen to be targeted by CRISPR/Cas9. Some genes that are not present in the phylogenetic tree (Figure S9B), but were found to be clustered in a specific subfamily by using iTAK, were selected to be knocked out.

**Table S2. Nucleotide sequences of the primers used in this study.**

| Primer code | Primer name | Sequence (5′ - 3′) |
|---|---|---|
| Primers used for introducing point mutations in the SOBIR1 kinase domain (the introduced mutations are indicated with a capital) | | |
| ho113 | NbSOBIR1_T512A fw* | gcactcccagatgcccatGcacatgttacgacttc |
| ho114 | NbSOBIR1_T512A rev | gaagtcgtaacatgtgCatgggcatctgggagtgc |
| ho115 | NbSOBIR1_T515A fw | gatgcccatacacatgttGcgacttcaaatgttgcagg |
| ho116 | NbSOBIR1_T515A rev | cctgcaacatttgaagtcgCaacatgtgtatgggcatc |
| ho117 | NbSOBIR1_T516A fw | gcccatacacatgttacgGcttcaaatgttgcaggaac |
| ho118 | NbSOBIR1_T516A rev | gttcctgcaacatttgaagCcgtaacatgtgtatgggc |
| ho119 | NbSOBIR1_S517A fw | gcccatacacatgttacgactGcaaatgttgcagg |
| ho120 | NbSOBIR1_S517A rev | cctgcaacatttgCagtcgtaacatgtgtatgggc |
| ho121 | NbSOBIR1_T522A fw | cttcaaatgttgcaggaGctgtgggatatattgcacc |
| ho122 | NbSOBIR1_T522A rev | ggtgcaatatatcccacagCtcctgcaacatttgaag |
| ao1 | SlSOBIR1_T503A fw | gcagttccagatgctcatGcacatatcaccacttc |
| ao2 | SlSOBIR1_T503A rev | gaagtggtgatatgtgCatgagcatctggaactgc |
| ao3 | SlSOBIR1_T506A fw | gctcatacacatatcGccacttcaaatgttgcag |
| ao4 | SlSOBIR1_T506A rev | ctgcaacatttgaagtggCgatatgtgtatgagc |
| ao5 | SlSOBIR1_T507A fw | caccGcttcaaatgttgcaggaactgttggatttattg |
| ao6 | SlSOBIR1_T507A rev | caataaatccaacagttcctgcaacatttgaagCggtg |
| ao7 | SlSOBIR1_S508A fw | gctcatacacatatcaccactGcaaatgttgcagg |
| ao8 | SlSOBIR1_S508A rev | cctgcaacatttgCagtggtgatatgtgtatgagc |
| ao9 | SlSOBIR1_T513A fw | caaatgttgcaggaGctgttggatttattgc |
| ao10 | SlSOBIR1_T513A rev | gcaataaatccaacagCtcctgcaacatttg |
| ho166 | SlSOB-like_T516A_fw | gctgtcccagatgctcatGcacatattacaacttc |
| ho167 | SlSOB-like_T516A_rev | gaagttgtaatatgtgCatgagcatctgggacagc |
| ho168 | SlSOB-like_T519A_fw | cagatgctcatacacatattGcaacttcaaatgtggc |
| ho169 | SlSOB-like_T519A_rev | gccacatttgaagttgCaatatgtgtatgagcatctg |
| ho170 | SlSOB-like_T520A_fw | gatgctcatacacatattacaGcttcaaatgtggcag |
| ho171 | SlSOB-like_T520A_rev | ctgccacatttgaagCtgtaatatgtgtatgagcatc |
| ho172 | SlSOB-like_S521A_fw | ctcatacacatattacaactGcaaatgtggcaggaac |
| ho173 | SlSOB-like_S521A_rev | gttcctgccacatttgCagttgtaatatgtgtatgag |
| ho174 | SlSOB-like_T526A_fw | cttcaaatgtggcaggaGctataggatacatcgctc |
| ho175 | SlSOB-like_T526A_rev | gagcgatgtatcctatagCtcctgccacatttgaag |
| ho123 | NbSOBIR1_Y355F fw | ggtggatgcggagaagtttTtagagctgagttaccggg |
| ho124 | NbSOBIR1_Y355F rev | cccggtaactcagctctaAaaacttctccgcatccacc |
| ho125 | NbSOBIR1_Y426F fw | ctaggccagactgccattTcttggtatatgaatatatg |
| ho126 | NbSOBIR1_Y426F rev | catatattcatataccaagAaatggcagtctggcctag |
| ho127 | NbSOBIR1_Y429F fw | gactgccattacttggtatTtgaatatatgaaaaatgg |
| ho128 | NbSOBIR1_Y429F rev | ccatttttcatatattcaAataccaagtaatggcagtc |
| ho129 | NbSOBIR1_Y431F fw | gccattacttggtatatgaatTtatgaaaaatgggagc |
| ho130 | NbSOBIR1_Y431F rev | gctcccatttttcataAattcatataccaagtaatggc |
| ho131 | NbSOBIR1_Y469F fw | gatagcttctggacttgagtTtctccatataaaccac |
| ho132 | NbSOBIR1_Y469F rev | gtggtttatatggagaAactcaagtccagaagctatc |
| ho133 | NbSOBIR1_Y525F fw | gcaggaactgtgggatTtattgcaccagaataccatc |
| ho134 | NbSOBIR1_Y525F rev | gatggtattctggtgcaataAatcccacagttcctgc |
| ho135 | NbSOBIR1_Y530F fw | gatatattgcaccagaatTccatcagacactgaag |
| ho136 | NbSOBIR1_Y530F rev | cttcagtgtctgatggAattctggtgcaatatatc |

| | | |
|---|---|---|
| ho137 | NbSOBIR1_Y543F fw | cgggtaagtgtgatatatTcagctttggtgtggtg |
| ho138 | NbSOBIR1_Y543F rev | caccacaccaaagctgAatatatcacacttacccg |
| ao11 | SlSOBIR1_Y346F fw | gtggctgtggagaagtttTtagagcagagctac |
| ao12 | SlSOBIR1_Y346F rev | gtagctctgctctaAaaacttctccacagccac |
| ao13 | SlSOBIR1_Y417F fw | gccagactgtcattTcttggtctacgaatacatg |
| ao14 | SlSOBIR1_Y417F rev | catgtattcgtagaccaagAaatgacagtctggc |
| ao15 | SlSOBIR1_Y420F fw | cttggtctTcgaatacatgaaaaatgggagtttacag |
| ao16 | SlSOBIR1_Y420F rev | ctgtaaactcccatttttcatgtattcgAagaccaag |
| ao17 | SlSOBIR1_Y422F fw | cgaatTcatgaaaaatgggagtttacaggatatcc |
| ao18 | SlSOBIR1_Y422F rev | ggatatcctgtaaactcccatttttcatgAattcg |
| ao19 | SlSOBIR1_Y460F fw | gctgctggtctcgagtTtctccatataaaccatac |
| ao20 | SlSOBIR1_Y460F rev | gtatggtttatatggagaAactcgagaccagcagc |
| ao21 | SlSOBIR1_Y521F fw | gcaccagaatTttatcagacactgaagtttacag |
| ao22 | SlSOBIR1_Y521F rev | ctgtaaacttcagtgtctgataaAattctggtgc |
| ao23 | SlSOBIR1_Y522F fw | gcaccagaatattTcagacactgaagtttac |
| ao24 | SlSOBIR1_Y522F rev | gtaaacttcagtgtctgaAaatattctggtgc |
| ao25 | SlSOBIR1_Y534F fw | gataagtgtgatatatTcagctttggtgtggtgc |
| ao26 | SlSOBIR1_Y534F rev | gcaccacaccaaagctgAatatatcacacttatc |
| ao27 | SlSOBIR1_Y588F fw | gcttataggaaatggatTcgacgaacaaatgc |
| ao28 | SlSOBIR1_Y588F rev | gcatttgttcgtcgAatccatttcctataagc |
| ho142 | SlSOB-like_Y359F_fw | cattgggcaaggtggatgtggaaaagtttTtaaagctgc |
| ho143 | SlSOB-like_Y359F_rev | ctttccgtcacttccaggtaatgcagctttaAaaacttttcc |
| ho144 | SlSOB-like_Y430F_fw | gccaagaccagactgccactTcttggtctatgagtacatg |
| ho145 | SlSOB-like_Y430F_rev | catgtactcatagaccaagAagtggcagtctggtcttggc |
| ho146 | SlSOB-like_Y433F_fw | ctgccactacttggtctTtgagtacatgaaaaatgggagc |
| ho147 | SlSOB-like_Y433F_rev | gctcccatttttcatgtactcaAagaccaagtagtggcag |
| ho148 | SlSOB-like_Y435F_fw | gccactacttggtctatgagtTcatgaaaaatgggagc |
| ho149 | SlSOB-like_Y435F_rev | gctcccatttttcatgAactcatagaccaagtagtggc |
| ho150 | SlSOB-like_Y473F_fw | gctgctggactcgagtTtctccatataaatcatactcagcg |
| ho151 | SlSOB-like_Y473F_rev | cgctgagtatgatttatatggagaAactcgagtccagcagc |
| ho152 | SlSOB-like_Y529F_fw | ggcaggaactataggatTcatcgctccagaatatcatcag |
| ho153 | SlSOB-like_Y529F_rev | ctgatgatattctggagcgatgAatcctatagttcctgcc |
| ho154 | SlSOB-like_Y534F_fw | ggatacatcgctccagaatTtcatcagacactgaagttcactg |
| ho155 | SlSOB-like_Y534F_rev | cagtgaacttcagtgtctgatgaAattctggagcgatgtatcc |
| ho156 | SlSOB-like_Y547F_fw | ctgataagtgtgatatatTcagctttggggtgctg |
| ho157 | SlSOB-like_Y547F_rev | cagcaccccaaagctgAatatatcacacttatcag |
| Primers for genotyping the *rlck-vii-6* knock-out plants | | |
| ho228 | 747_geno_1-F_JS2419 | CCAAAACCAAAGGCAAAGGAG |
| ho229 | 747_geno_1-R1004_JS2420 | GATTAGCCAGGTAGATAGTGG |
| ho230 | 747_geno_1-R5004_JS2421 | GACTGAAGTTCCAAAAGCC |
| ho231 | 747_geno_2-F_JS2422 | GTGATGAAGATTGCATGGG |
| ho232 | 747_geno_2-R2008_JS2423 | GTTTACGAGTTATTGCACGAT |
| ho233 | 747_geno_2-R3011_JS2424 | CACCAACTCCCAAAAGCC |
| ho234 | 747_geno_3-F3002_JS2425 | GTGGTCATTCAAGGCTAGC |
| ho235 | 747_geno_3-R3002_JS2426 | TTTTCTTAAACTCCGTGCCG |
| ho236 | 747_geno_4-F13015_JS2427 | TCATCATCATAGTAGGCTAGCC |
| ho237 | 747_geno_4-R13015_JS2428 | TAGTCCTTTTGCAGCACCTA |
| ho238 | 747_geno_5-F_JS2429 | CGTAGGTTGTCATTCTCGGA |
| ho239 | 747_geno_5-R1006_JS2430 | TGTAACTTCTTGGGGTACGT |

| | | |
|---|---|---|
| ho240 | 747_geno_5-R0005_JS2431 | TGCGTACCATCAAAACAACC |
| ho241 | 747_R1-13015_JS2486 | TGTTTTCTCTTACACACCCC |
| ho242 | 747_F2-13015_JS2487 | TGAATACATGGCAAGGGGAA |
| Primers for genotyping the *rlck-vii-7* knock-out plants | | |
| LHo07 | 734_00009F_JS2556 | CCAGTTCTTCTGGTCTACG |
| LHo08 | 734_03007F_JS2555 | CCTGTTCTTCTGATCTATG |
| LHo09 | 734_03007R+00009R_JS2557 | CAACTAAGCAGACCTTACC |
| LHo10 | 734_01025F_JS2558 | TCCATTGTTCAGACACAGTG |
| LHo11 | 734_06004F_JS2559 | CATTGCTCAGAGATAGTGAG |
| LHo12 | 734_06004R+01025R_JS2560 | GATGCTCAGAAAGAAGAGG |
| LHo13 | 734_02015F_JS2551 | ACAAAAGGACCCAAATTGCT |
| LHo14 | 734_02015R_JS2552 | TGGAGTAAGCGTCTTCTCAT |
| LHo15 | 734_03005F_JS2553 | CTACGTACGCCTCTTACAGA |
| LHo16 | 734_03005R_JS2554 | CGGTGATTGCTTGTATCGAT |
| LHo17 | 734_13012F_JS2561 | TAGCAACAGCACAACTTGTT |
| LHo18 | 734_13012R_JS2562 | TCCTTGAGACCTTGTTCACT |
| Primers for genotyping the *rlck-vii-8* knock-out plants | | |
| Po05 | Scf00012 | AAATTAGAACACGGACAAT |
| Po01 | Scf00229_Fw | GCTGCAAAGATAGTAGCAATTGG |
| Po02 | Scf00229_Rv | GAAGCACGCAGATAGGCG |
| Po03 | Scf00635_Fw | GCAATTGGCTTTTGTATGTGC |
| Po04 | Scf00635_Rv | GCGACCTAATAATAGTTCAAAATTGC |
| Po36 | Scf1025_Fw | GACTAGTAATTCGAACAGC |
| Po37 | Scf1025_Rv | TCCCTAATCTGTATATGGGG |
| Po09 | Scf01378 | AAATTAGGACACGGGCAAC |
| Po08 | Scf02819 | TTTCGGTTGAACCCGTGGA |
| Po10 | Scf03493 | TTTCGGTTGAACTCATAGC |
| Po11 | Scf05476_FW | GTAATTGGTTTCTTTGGATGTGTATC |
| Po14 | Scf05476_Rv | GCGAATCAAGTCGCCTATTCATCA |
| Po15 | Scf6482_Fw | GGTGTCATGAGAAGTGCAGATTGC |
| Po16 | Scf6482_Fw | CCAATTTTACTACGAAGTACCATGCG |
| Po06 | Scf06482_Fw | GCAAAGATAGTAGTAATTGGCTTTTGTATGTGC |
| Po07 | Scf06482_Rv | GCAAATGAATGTAATTTTGAACTAGTATTACGTGC |
| Po12 | Scf0012-1378_Fw | ATGTTGCTATCTCTGGTTT |
| Po13 | Scf02869-3493_Rv | GCCTGTGTTCATCTTCTAAG |
| Primers used for generating *E. coli* expression constructs of the SOBIR1 kinase domain (sequences that are identical to the expression vectors are underlined) | | |
| ho201 | pET-GST_fw | aagcttgcggccgcactcgag |
| ho202 | pET-GST_rev | caggggcccctggaacagaacttc |
| ho203 | NbSOBIR1_KD+90bp_fw | gttctgttccaggggcccctgcgaaagggaaagactgatggaac |
| ho204 | NbSOBIR1_KD+90bp_rev | ctcgagtgcggccgcaagcttctaatgcttgatctgagttaacatac |
| ho215 | SlSOBIR1_KD+90bp_fw | gttctgttccaggggcccctgagaagagggaataacgattcaag |
| ho216 | SlSOBIR1_KD+90bp_rev | ctcgagtgcggccgcaagcttctaatgcttgatctgagttaacatgc |
| ho217 | SlSOBIR1-like_KD+90bp_fw | gttctgttccaggggcccctgagagggatcagaaatgatccagg |
| ho218 | SlSOBIR1-like_KD+90bp_rev | ctcgagtgcggccgcaagcttttaatgcttgatctgcatcaacatgc |
| ho219 | pET-15b_fw | cggatcctcgagcatatggctg |
| ho220 | pET-15b_rev | gctgctaacaaagcccgaaaggaag |
| ho221 | NbSO_KD+90bp_fw | gccatatgctcgaggatccgcgaaagggaaagactgatggaac |
| ho222 | NbSO_KD+90bp_rev | ctttcgggctttgttagcagcctaatgcttgatctgagttaacatac |

| | | |
|---|---|---|
| ho223 | SlSO_KD+90bp_fw | gccatatgctcgaggatccgagaagagggaataacgattcaag |
| ho224 | SlSO_KD+90bp_rev | ctttcgggctttgttagcagcctaatgcttgatctgagttaacatgc |
| ho225 | SlSO-like_KD+90bp_fw | gccatatgctcgaggatccgagagggatcagaaatgatccagg |
| ho226 | SlSO-like_KD+90bp_rev | ctttcgggctttgttagcagcttaatgcttgatctgcatcaacatgc |
| ho282 | NbSERK3b_kd_fw | gccatatgctcgaggatccgggacaactcaagaggttttccttg |
| ho283 | NbSERK3b_kd_rev | ctttcgggctttgttagcagctcatcttggccccgacaattcatc |
| ho286 | SlSERK3a_kd_fw | gccatatgctcgaggatccgggacaactcaaaaggttttccttgc |
| ho287 | SlSERK3a_kd_rev | ctttcgggctttgttagcagctcatcttggccctgacaactcatc |

* fw, forward; rev, reverse.
